# Three-dimensional envelope and subunit interactions of the plastid-encoded RNA polymerase from *Sinapis alba*

**DOI:** 10.1101/2022.05.04.490580

**Authors:** Rémi Ruedas, Yohann Couté, Sylvie Kieffer-Jacquinod, Soumiya Sankari Muthukumar, François-Xavier Gillet, Daphna Fenel, Grégory Effantin, Thomas Pfannschmidt, Robert Blanvillain, David Cobessi

## Abstract

RNA polymerases (RNAPs) involved in gene transcription are found in all living organisms with degrees of complexity ranging from single polypeptide chains to multimeric enzymes. In the chloroplasts, the nuclear-encoded RNA polymerase and the plastid-encoded RNA polymerase (PEP) are both involved in the selective transcription of the plastid genome. The PEP is a prokaryotic-type multimeric RNAP found in different states depending on light stimuli and cell identity. One of its active states requires the assembly of nuclear-encoded PEP-Associated Proteins (PAPs) on the catalytic core, producing a complex of more than 900 kDa regarded as essential for chloroplast biogenesis. A purification procedure compatible with structural analysis was used to enrich the native PEP from *Sinapis alba* chloroplasts. Mass spectrometry (MS)-based proteomic analysis identified the core components, the PAPs and additional members, and coupled to cross-linking (XL-MS) provided initial structural information about the relative position of PEP subunits. Sequence alignments of the catalytic core subunits across various chloroplasts of the green lineage and prokaryotes combined with structural data show that the overall shape of the catalytic core and the residues essential for the catalytic activity are conserved. However, variations are observed at the surface of the core, some of them corresponding to PAP binding sites as suggested by XL-MS experiments. Using negative stain electron microscopy, the PEP 3D envelope was calculated. 3D classification shows that the protrusions which we attribute to the PAPs are firmly associated with the catalytic core. Overall, the shape of the *S. alba* PEP envelope is different from that of RNAPs.

## Introduction

DNA-dependent RNA polymerases (RNAPs) are central enzymes of gene expression which transcribe the genetic information encoded in DNA into single-stranded RNAs; some of which suitable for translation. RNAPs exist in highly varying degrees of complexity ranging from single subunit enzymes in T3/T7 phages to highly multimeric enzymes in eukaryotes. Eubacterial multimeric RNAPs share a common catalytic core composed of two large subunits called β and β’ a dimer of α subunits and a monomer of the ω subunit (Cramer, 2002; Hirata *et al*., 2008; Murakami, 2015). For specific transcriptional activity, RNAPs require additional proteins such as σ factors that mediate recognition of gene promoters and are essential to initiate transcription. The three-dimensional structures of RNAPs have been solved for eukaryotic and prokaryotic RNAPs in several states (Murakami, 2015; Lee & Borukhov, 2016; Hanske *et al*., 2018). Structure comparisons and sequence alignments have shown that, although sequence identities can be low, the overall shape of the five core subunits is largely conserved among the RNAPs (Murakami, 2015). Furthermore, several homologous regions have been identified between the bacterial and eukaryotic RNAPs suggesting that the fold is better conserved than the amino acid sequences. Essential residues and regions for effective transcription are, however, conserved among RNAPs, indicating that the enzymes share a common transcription mechanism (Cramer, 2002). In eukaryotes, several RNAPs are involved in transcription of nuclear genes (RNAPs I, II and III) while a specific phage-type RNAP transcribes the mitochondrial DNA. Plant cells are unique among eukaryotes since they possess a third genetic compartment, plastids. These endosymbiotic organelles originate from the engulfment of an ancient cyanobacterium into a mitochondriate proto-eukaryote around 1.5 billion years ago (Bobik & Burch-Smith, 2015) which was followed by a massive transfer of the cyanobacterial genes into the nucleus of the host cell (Martin *et al*., 2002). As a result, most plastome (chloroplastic DNA (cpDNA)) of today’s plastids contains only about 120 genes (Sugiura, 1992) encoding (i) components of the plastid gene expression machinery (the core subunits of the prokaryotic-type RNA polymerase, ribosomal proteins, tRNAs, and rRNAs), (ii) subunits of each of the major functional photosynthesis-related complex (*e*.*g*., ribulose-1,5-bisphosphate carboxylase/oxygenase (RuBisCO), photosystem I and II (PSI and PSII), cytochrome *b*_6_*f* complex, NADH dehydrogenase, and the ATP synthase), and (iii) a few proteins involved in other essential processes such as protein import, fatty acid synthesis or protein homeostasis (*e*.*g*., YCF1 and 2, AccD, ClpP1) (Sugiura, 1992; Majeran *et al*., 2012; Yu *et al*., 2014). Despite the limited coding capacity of the plastome, chloroplasts contain 2,500-3,500 different proteins (Zybailov *et al*., 2008), thus the vast majority of chloroplast proteins is encoded by the nuclear genome and must be post-translationally imported. The expression of the cpDNA is, however, essential to chloroplast biogenesis and functions since genetic impairments of plastid gene expression result in albinism (Pfalz & Pfannschmidt 2013).

Transcription of the plastome involves a single-subunit <u>n</u>uclear-<u>e</u>ncoded T3/T7 phage-type RNA <u>p</u>olymerase (NEP) and the multi-subunit <u>p</u>lastid-<u>e</u>ncoded prokaryotic-type RNA <u>p</u>olymerase (PEP). Briefly, the NEP enzyme transcribes the so-called ‘house-keeping’ genes (including *rpo* genes encoding the core subunits of the PEP) while the PEP transcribes preferentially genes encoding proteins of the photosynthetic complexes as well as tRNA genes (Hajdukiewicz *et al*., 1997; Williams-Carrier *et al*., 2012). However recent studies have uncovered that many plastid genes possess promoters for NEP and PEP and, thus, can be transcribed by both RNA polymerases (Weihe & Börner, 1999) (Figure S1). Furthermore, the division of labor between the two RNA polymerases changes with developmental stage and a clear-cut separation between NEP and PEP transcribed genes remains difficult (Zelyaskova *et al*., 2012). The catalytic core enzyme of PEP comprises four subunits called α, β, β’ and β’’ encoded by the genes *rpoA, rpoB, rpoC1* and *rpoC2*, respectively (Börner *et al*., 2015; Pfannschmidt *et al*., 2015). Biochemical studies performed in mustard revealed that in etioplasts (the plastid type found in dark grown seedlings), these subunits assemble to form the prokaryotic-like enzyme PEP-B (Pfannschmidt *et al*., 2000; Yagi & Shiina, 2014). In angiosperm, seedlings illumination initiates a light signaling cascade that triggers photomorphogenesis and chloroplast biogenesis. This involves a structural re-organization of the PEP-B enzyme by association of additional subunits resulting in a much larger multi-subunit PEP-A complex. Biochemical purifications performed in several plants revealed that the complex comprises at least 16 different proteins with an overall molecular mass of more than 900 kDa (Pfannschmidt *et al*., 2000; Suzuki *et al*., 2004; Steiner *et al*., 2011). MS analyzes done with mustard PEP-A complexes identified 12 PEP-associated proteins (PAPs) that are stably bound to the complex. These PAPs are all encoded by the nuclear genome and must be imported from the cytosol. Genetic inactivation of any of these 12 PAPs results in a severe block or disturbance of chloroplast biogenesis, indicating that this re-organization of the PEP complex represents a crucial developmental step in chloroplast biogenesis (Garcia *et al*., 2008; Arsova *et al*., 2010; Chen *et al*., 2010; Gao *et al*., 2011; Steiner *et al*., 2011; Yagi *et al*., 2012; Yu *et al*., 2012; Pfalz & Pfannschmidt 2013; Yu *et al*., 2014).

In contrast to RNAPs I, II and III, for which several 3D structures were solved, nothing is known about the PEP 3D structure. It is generally assumed that the PEP core enzyme resembles that of the bacterial RNA polymerase based on the homologous regions found in bacterial and plastidial *rpo*-genes (Pfannschmidt *et al*., 2015). However, no experimentally obtained structural 3D data of this complex exist. So far, only structural predictions of PAPs have been performed based on their primary amino acid sequences and structure database of proteins with similar functional domains, and recently the 3D structure of PAP9 was solved (Favier *et al*., 2021). It is, however, completely unknown whether these predictions reflect reality and where and how these additional subunits associate with the PEP core enzyme.

Here, we report the characterization of the PEP complex purified from *S. alba* cotyledons. MS-based proteomic analysis identified all known PEP subunits and additional members. A chemical crosslinking coupled to MS approach highlighted some interacting neighbors in the PEP complex. Using negative stain electron microscopy, we calculated the first 3D envelope of the PEP-A complex showing together with the MS analyses that the PAPs are firmly associated with the catalytic core by probably binding to non-conserved regions as revealed by sequence alignments between the catalytic core of PEP from angiosperms and bRNAPs.

## Results

### The PEP complex and its associated proteins

We used a MS-based label-free quantitative proteomic analysis to characterize the *S. alba* PEP-enriched fraction isolated from mustard cotyledon chloroplasts following an established purification scheme with slight modifications. More than 400 different proteins were reproducibly identified and quantified in three independent preparations of PEP (Table S1). Their relative abundances within the PEP fraction were approximated using their extracted iBAQ values (Schwanhäusser *et al*., 2011), showing that these proteins were distributed over four orders of magnitude. Among the 24 most abundant proteins, representing ∼60% of the total amounts of proteins within the fraction, we identified the four core subunits (α, β, β’ and β’’) and the twelve PAPs (Figure 1 and Figure S2). The α subunit was found to be ∼ twice more abundant than the β subunit, consistent with a stoichiometry of two α subunits per one β subunit in the catalytic core complex, as known for eubacterial RNAPs. Beside of these 16 known PEP subunits we identified fructokinase-like protein 2 (FLN2) and PTAC18 (Pfalz *et al*., 2006), two other proteins already known to be part of the plastid nucleoid. Two other proteins present in this shortlist were a protein homolog of *A. thaliana* At4g36700 corresponding to a late embryogenesis abundant protein of the RmlC-like cupin superfamily and the chloroplastic ribosomal protein Rps7; both of which may be found due to high abundance in the young seedling (cupin) or close proximity (anchoring) to the translational machinery within the chloroplast (Rps7). The remaining proteins identified among the 24 high abundant candidates all belong to the family of histones suggesting that nucleosomes co-purify in the PEP fraction likely due to nuclei associated with the chloroplast envelopes. All other proteins detected in the PEP-enriched fractions represent low abundant co-purified proteins probably identified because of the high sensitivity of the MS. They, however, did not interfere with the main goal of our study, the structural analysis of the PEP complex (see further below).

**Figure 1:**
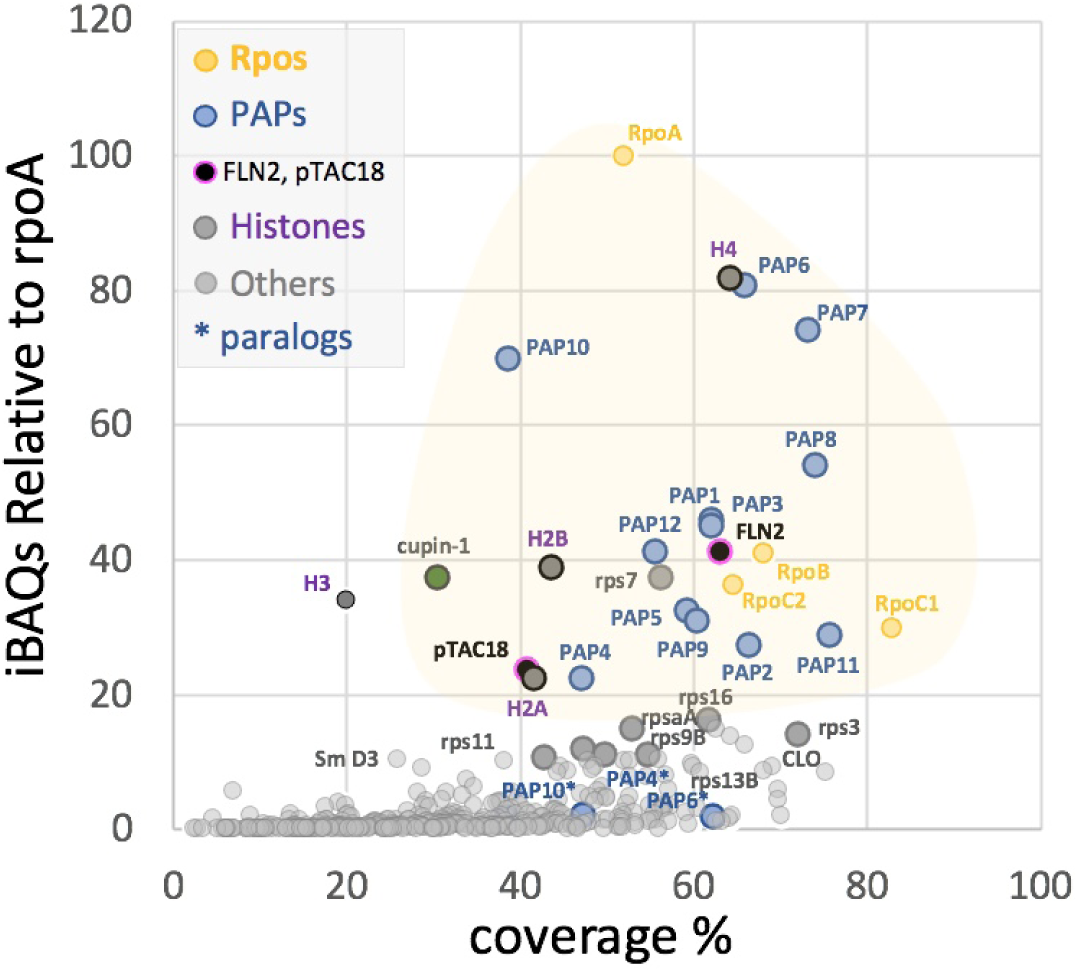
mass spectrometry data presented as iBAQ values relative to that of rpoA (iBAQr) as function of the corresponding protein coverage percentage. RpoA, B, C1 and C2 (subunits α, β, β’ and β’’) are in yellow, PAPs in blue, suspected permanent residents in dark, histones in magenta, and suspected purification contaminants in different shades of grey. In the pink area fall all the expected components of the PEP-A complex and correspond to the major protein mass contribution to the purified sample.

In order to obtain initial structural information about the relative position of the PEP subunits within the complex, we used a combination of biochemical cross-linking coupled to MS. To this end, we treated the PEP-enriched fraction from two independent purifications (replicates 2 and 3 of the preparations used for the proteomic discovery part) with Disuccinimidyl Dibutyric Urea (DSBU) before tryptic digestion and MS analyses. This strategy allowed to reliably identify 39 inter-protein dipeptides from which 12 contained PEP core subunits or PAPs, suggesting a spatial proximity between these subunits within the PEP complex (Table S2). Apparently, the core subunits were partly accessible to the DSBU treatment since two dipeptides linking the β and β’ subunits were identified, suggesting that the associated PAPs do not cover the core completely but leave some gaps that allow the crosslinker molecules to access the core. Structure analyses of the RNAPs from *E. coli* (PDB entries: 3LU0 (Opalka *et al*., 2010) and 6GH5 (Glyde *et al*., 2018)) and *T. thermophilus* (PDB entry: 6ASG (Lin *et al*., 2018)) do not allow to model the dipeptides observed suggesting that these regions in PEP have probably different conformations of those observed in these 3D structures since sequences are conserved (Figures S3-S6). PAP5 and PAP6 were found to both interact with the a subunits, each of them likely interacting with a different monomer since their interaction was within the same peptide of the a subunit (Table S2, Figure 2). Another region of PAP5 was found in close vicinity of the KNYQNER peptide of the β’ subunit (Table S2) that belongs to an insertion of conserved residues found only in angiosperms after the β’a12 domain (Figure S5). This result suggests that surface localized residues that are not conserved between the catalytic cores of bRNAPs and PEP but conserved in plants have evolved towards the interactions with the PAPs (see below). We also found a PAP5/PAP6 dipeptide suggesting that the α, β, β’ subunits, PAP5 and PAP6 may form a structural cluster within the fully assembled PEP complex. A second cluster appears to be formed by PAP1, PAP2 and PAP11/MurE-like for which dipeptides were also found.

**Figure 2:**
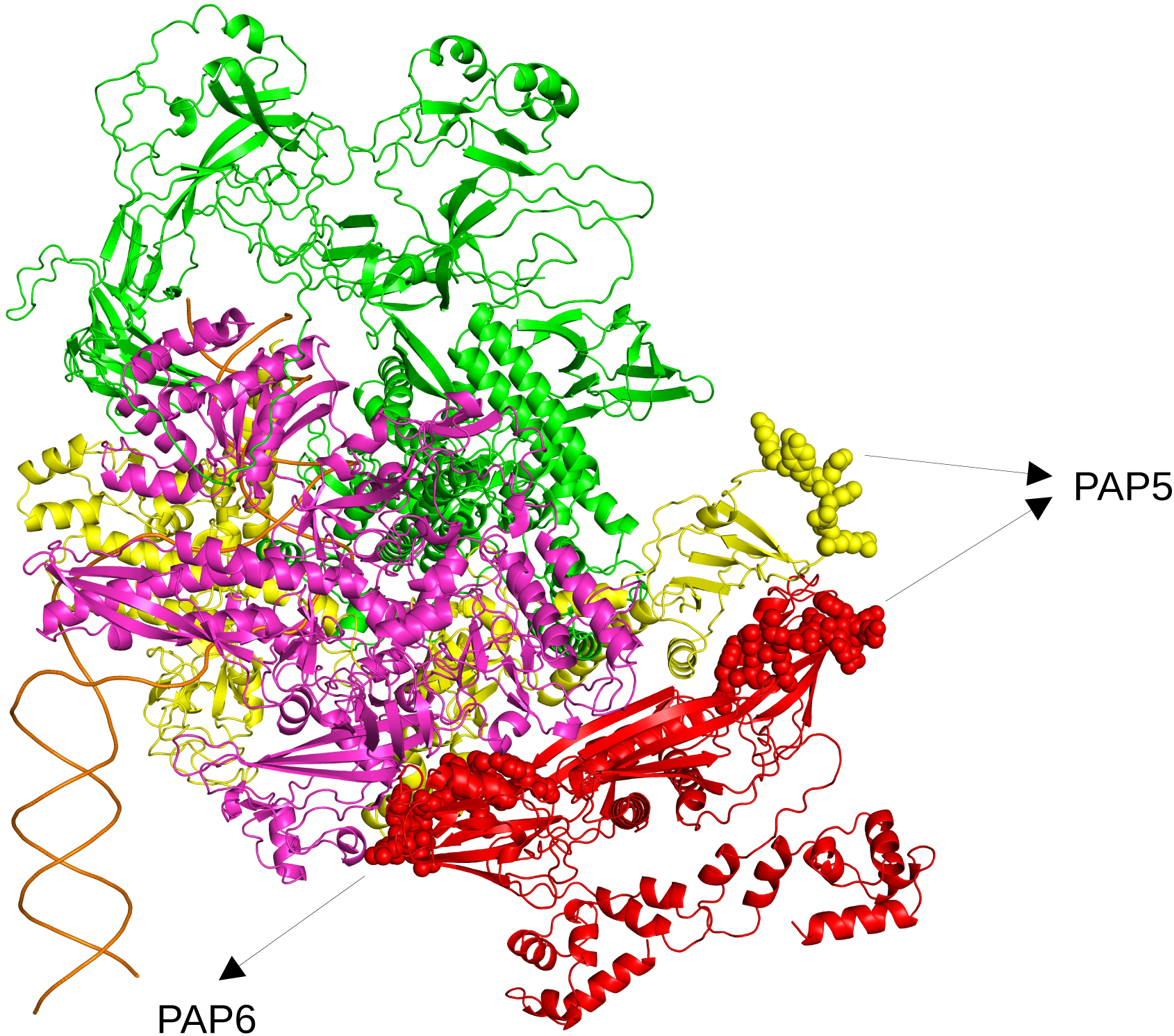
model of the PEP core from *A. thaliana* built from the α, β, β’ and β’’ subunits modeled using AlphaFold (Jumper *et al*., 2021) and superimposed onto the *E. coli* RNAP catalytic core and colored as followed: α subunit in red, β subunit in purple, β’ subunit in yellow and the β’’ in green. The van der Waals spheres display the peptides of the α and β subunits that are nearby to PAP5 and PAP6.

### Patches of mutations in the PEP at the surface of the catalytic core

To highlight differences in the PEP core complex compared to eubacterial RNAPs which could be evolutionary linked to the PAP interactions, we performed a detailed sequence alignment analysis of the α, β, β’ and β’’ core subunits from various species chosen in the tree of the green lineage (Figures S3-S6) as proposed by Finet *et al*. (Finet *et al*., 2010). These sequences were found to be well conserved within the green lineage (Figures 3a and 3b and figures S3-S6). The weakest sequence identity is observed when comparing the sequences from *Physcomitrium* to those of other species; the sequence of the a subunit being the most divergent. Sequence conservation appears to be high in the domains of the β, β’ and β’’ subunits that bear the catalytic activity while the α subunits are responsible for the assembly of the core (Sutherland & Murakami, 2018). Sequence comparisons with RNAPs from bacteria and cyanobacteria reveal also that regions essential to the transcription activity are conserved and the bacterial β’ subunit can be aligned with the β’ and β’’ subunits of the PEP. Whereas the catalytic activity is carried by the β and β’ subunits in bRNAPs, it is supported in the PEP by the β, β’ and β’’ subunits. Unlike in *E. coli*, the β subunit of the PEP does not have the additional βi4, βi9 and βi11 domains (Opalka *et al*., 2010). However, the β’’ subunit of the PEP contains a long plant specific insertion of several hundred residues between the region β’b8 and β’b9 that does not exist neither in β’ subunit from *E. coli* RNAP nor in the β’ subunit from *T. thermophilus* RNAP (Figure 3b and figure S6). The β’’ subunit of RNAP from angiosperms lacks also a part of the β’b10 region observed in the RNAP from *Nostoc*. Most of the strictly conserved residues described for the catalytic core of RNAPs (Lane & Darst, 2010), however, are conserved in PEP. When plotting the sequences into a *E. coli* 3D model most of the mutated residues in the PEP sequences are located at the surface of the catalytic core of the bRNAP supporting the assumption that they may be required for the interaction with PAPs (Figure 4).

**Figures 3:**
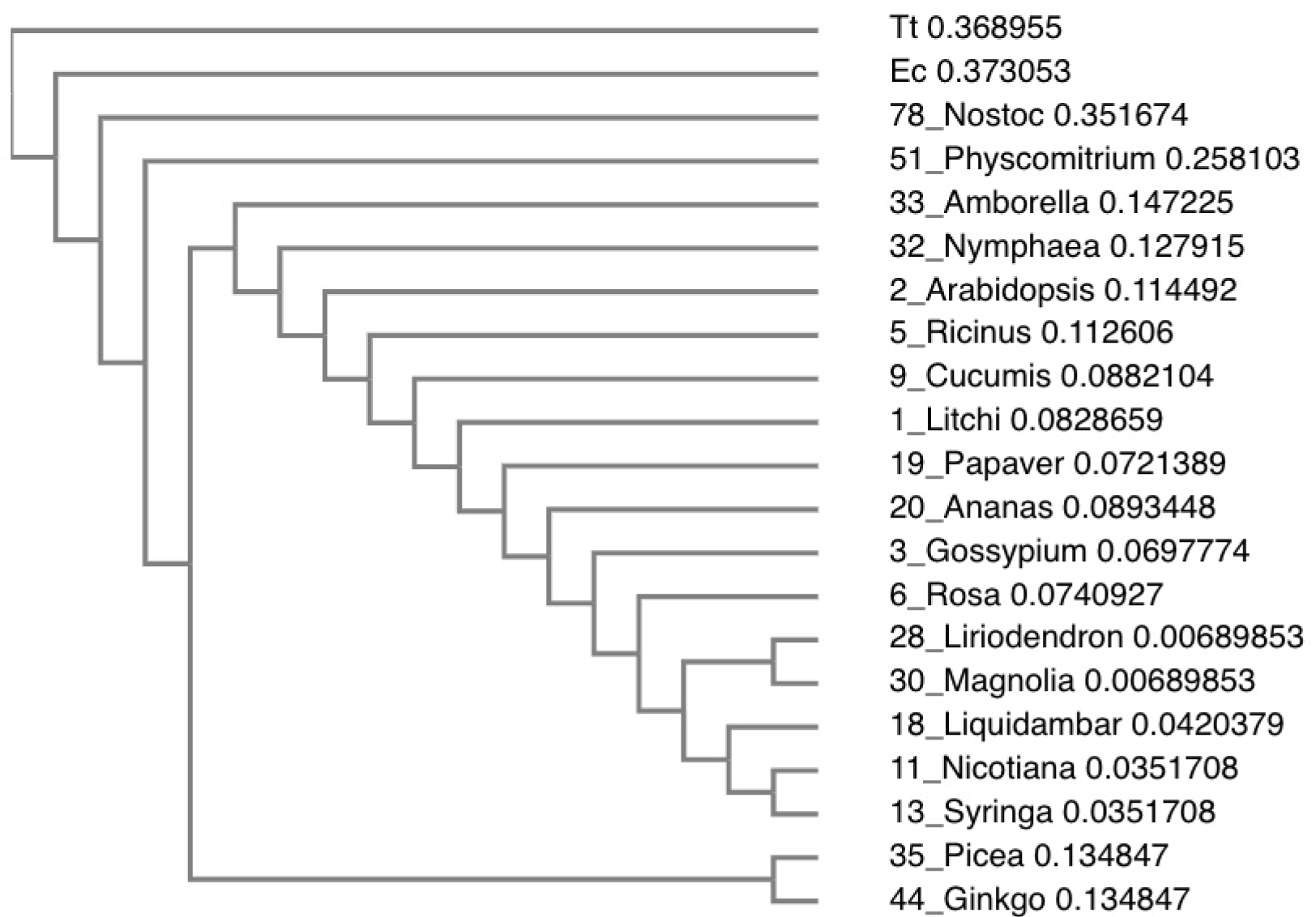
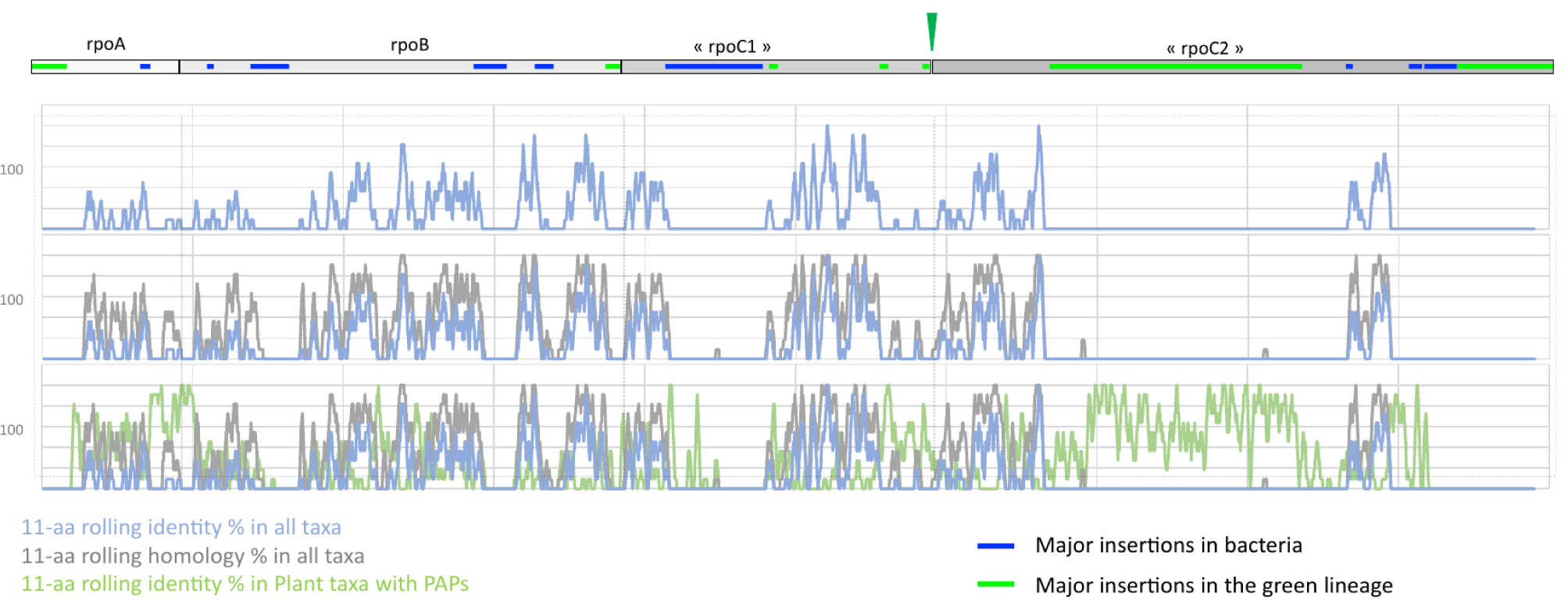
a) phylogram obtained with Clustal Omega multi alignment algorithm. Branch length presented as cladogram. Major taxa included from the collection presented in data source (excel sheet sorted). A major incongruence from the angiosperm phylogeny tree (version IV: http://www.mobot.org) is noted for Magnoliales and likely due to the study of chloroplast genes with cytoplasmic inheritance. b) Global alignment represented as 11-aa rolling identity (blue) or homology (grey) percentages calculated for all taxa. In green, the 11-aa rolling identity percentage calculated in a subset of taxa corresponding to plants with detected PAPs (green). Green triangle is the evolutionary split of the *rpoC* gene in *rpoC1* and *rpoC2* genes in cyanobacteria.

**Figure 4:**
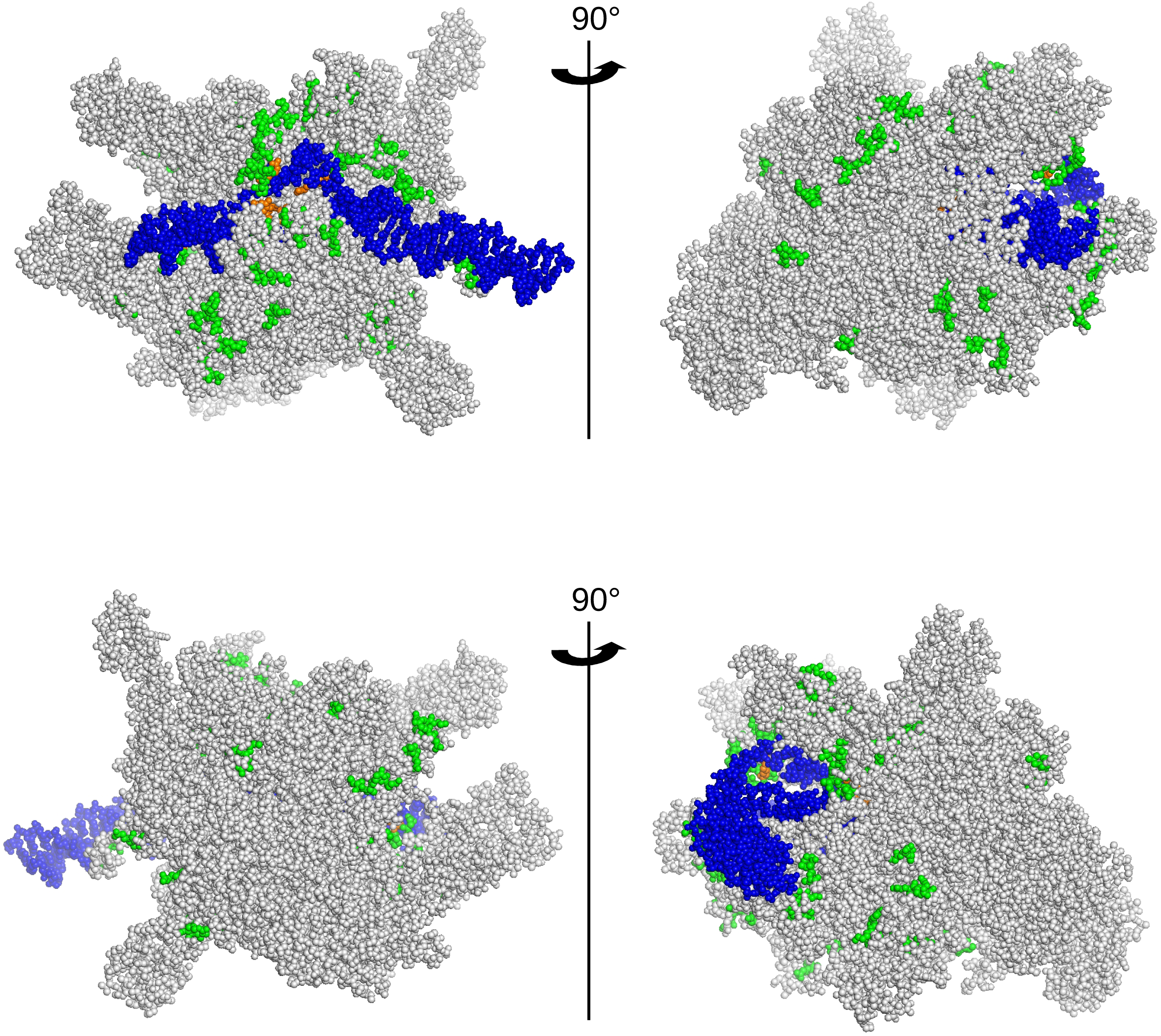
view of the *E. coli* RNAP (PDB entry: 6GH5) without the ω subunit and the σ54 factor. The double stranded DNA is colored in blue. The core subunits are drawn in sphere. The residues colored in green and orange are those mutated as given in the sequence alignments (Figures S3-S6).

### A chloroplastic catalytic core surrounded with nuclear encoded proteins

We then investigated the 3D structure of the fully assembled PEP complex by using negative stain electron microscopy. Overview images of the stained complexes displayed well separated molecules of various shapes, but very limited aggregation (Figure 5a) and no disturbance by other complexes (such as nucleosomes). The homogeneity of the sample was probed by *ab initio* 2D classification of the individual complex images that revealed several well-defined 2D classes (Figure 5b). The overall shapes of the classes are multiple but they are all consistent in sizes with dimensions varying between 150 and 280 Å. Some 2D classes of PEP displayed a more compact center, sometimes with a clear stain filled pocket surrounded by several protrusions of various sizes (Figure 5b). From the particles isolated by 2D classification, a 3D map at 27.5 Å resolution could be determined (Figure 5c) which recapitulates the features seen in the 2D classes such as the central cavity (depression) and the peripheral protrusions.

**Figure 5:**
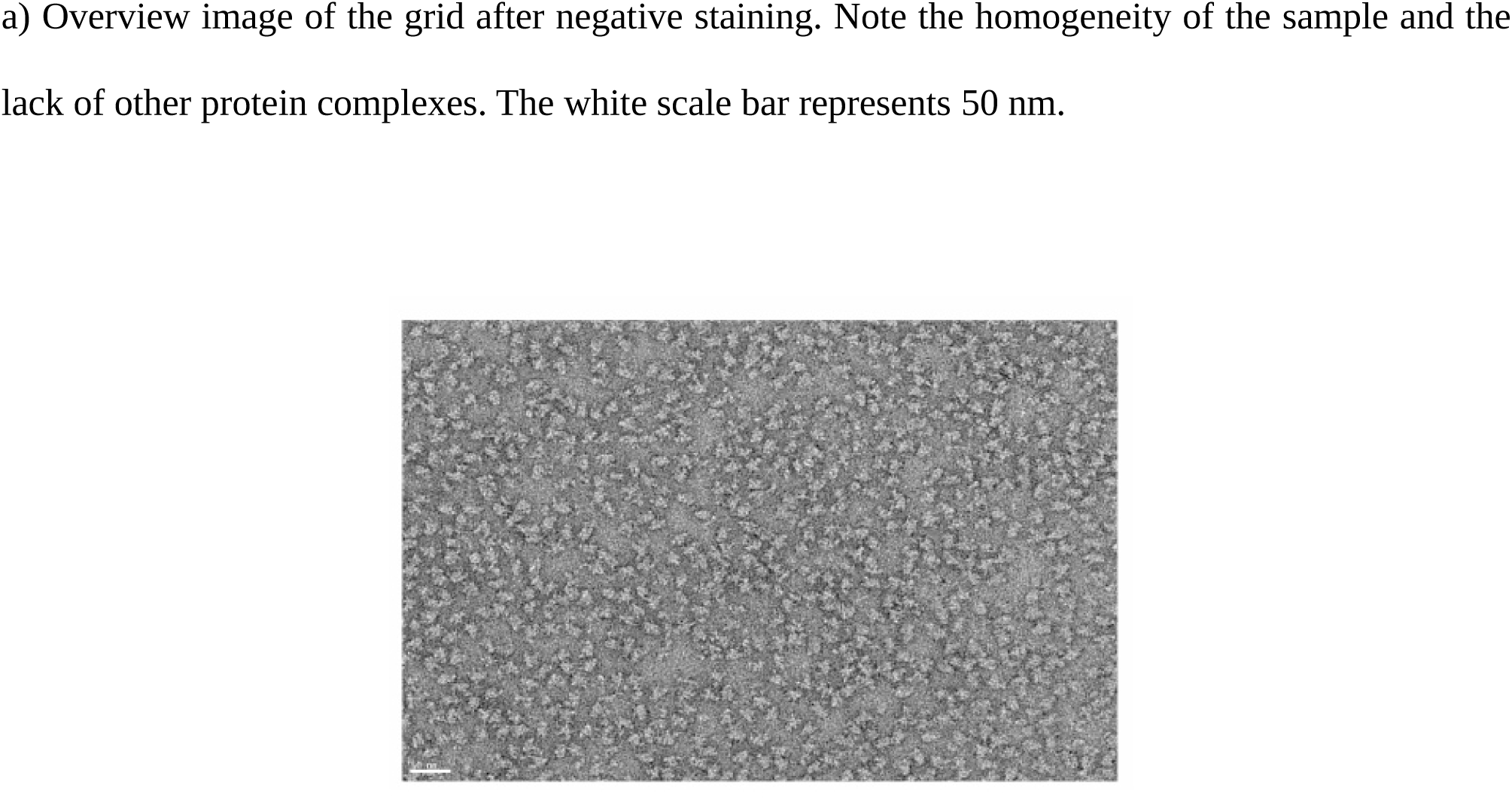
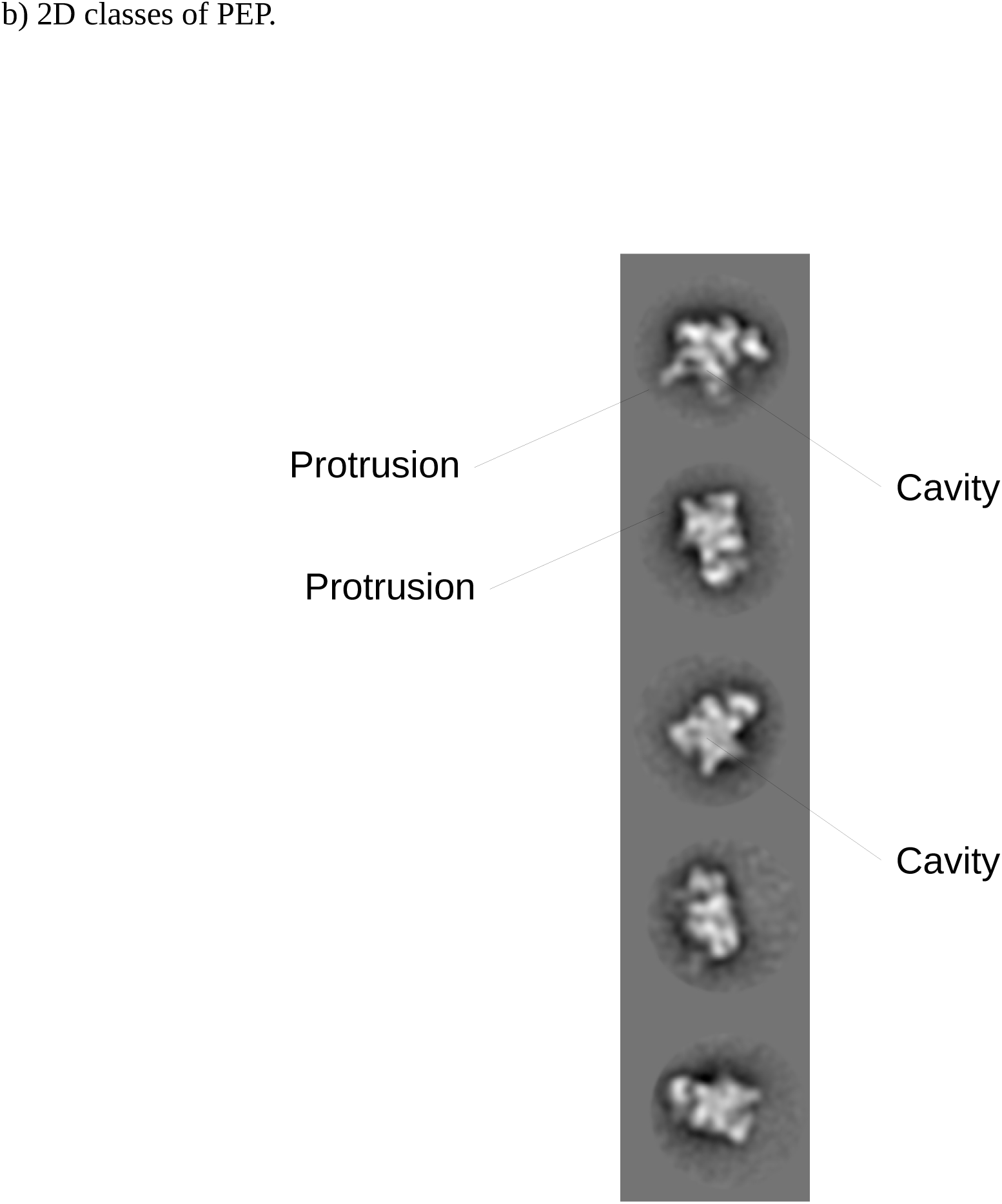
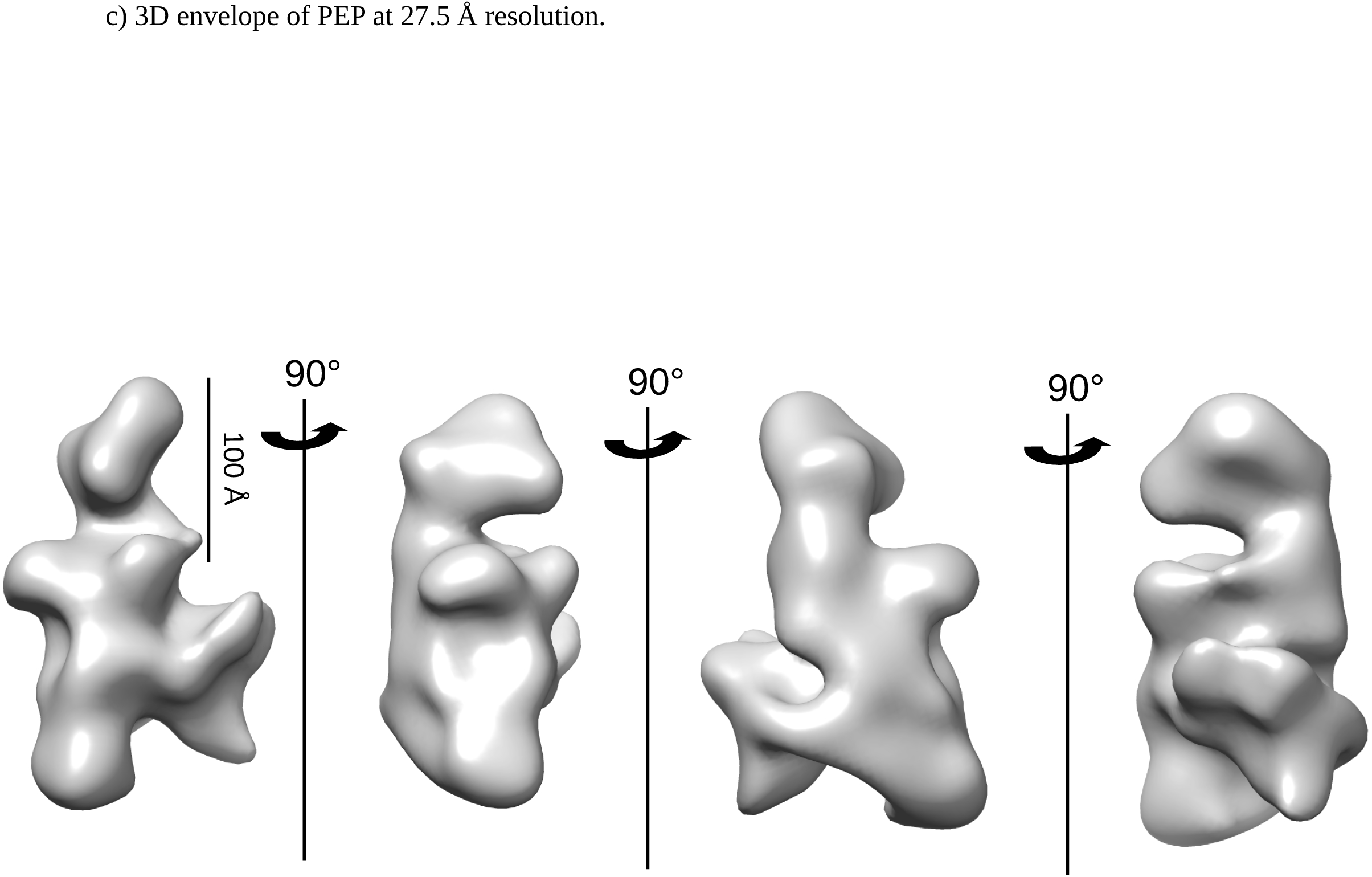
negative staining electron microscopy and 3D envelope of PEP.

## Discussion

The purification protocol used in this study allowed us to retrieve a stable PEP complex with limited amount of contaminant proteins. The core subunits and previously described PEP associated proteins are the most abundant proteins.

The three MS-based proteomic characterizations of *S. alba* PEP fraction revealed the presence of FLN1 (PAP6) and FLN2, two fructokinase-like proteins whose gene deletion lead either to an albino phenotype or that exhibits a delayed greening, respectively (Gilkerson *et al*., 2012). FLN2 is the paralogous protein of FLN1 and both do not exhibit sugar-phosphorylating activity although they contain a fructokinase domain (Arsova *et al*., 2010). They can form homodimers or heterodimers *in vitro* (Riggs & Callis, 2017). Characterization of proximal proteins in *S. alba* PEP fraction using XL-MS showed that FLN1 or FLN2 interact with the α subunit of the catalytic core. Based on the sequence, it is not possible to distinguish which FLN paralog binds to the α subunit due to the high sequence identity between FLN1 and FLN2 that display the same peptide sequence identified. The part of the α subunit observed in this interaction (GY(157)SLK(160)MSNNFEDR) is the same that also interacts with PAP5; involving Y157 and K160 in the dipeptide bond with PAP5 and FLN1/FLN2 respectively. Considering that the complex has a homogenous structure with correctly positioned partners, steric hindrance would not allow for two proteins with predicted different folds (PAP5 and FLN1 or FLN2) to interact with the same region of the α subunit. The MS-based proteomic characterization of *S. alba* PEP fraction suggested also that the a subunit is ∼ twice more abundant than the β subunit. Together, these observations are consistent with a stoichiometry of two subunits per one subunit (Sutherland, & Murakami, 2018) in the core of PEP as observed in the eubacterial RNAPs supporting the assumption that the PEP core resembles that of bRNAPs. In the PEP, PAP5 and PAP6 (or FLN2) form a first cluster with the α, β, β’ subunits (Table S2), suggesting that they are associated first during a PEP-B to PEP-A transformation.

MS-based proteomic characterization using XL-MS revealed also the presence of two other closely related proteins: PAP1 and PAP2. PAP1 and PAP2 have pentatricopeptide repeats involved in RNA binding. Among the PAPs with predicted nucleic acid binding domains, PAP1 possess a SAP domain known for DNA or RNA binding, while PAP3 has a S1-like domain predicted to interact with RNA (Pfannschmidt *et al*., 2015). Since dipeptides between PAP1 and PAP2 are found, both proteins are in close proximity in the PEP, PAP1 being also involved in interactions with PAP11/Mure-like (Table S2), the three proteins forming a second cluster.

The presence of closely related proteins, such as PAP6 and FLN2 or PAP4 and PAP9, two superoxide dismutases, raise the question of the PEP subunit composition. Even if the 3D classifications did not reveal any significant variability in the 3D envelope, PEP heterogenous complexes could exist. Furthermore, the PEP complex of our preparation could contain additional subunits like FLN2 or PTAC18 not observed previously in gel-based MS analyses. It remains open whether these subunits represent loosely or tightly associated PEP subunits. Initial discovery of PTAC18 in the TAC already placed this protein conceptually close to the PEP (Pfalz *et al*., 2006). Further biochemical analyses associated to high-resolution cryo-EM map of the PEP and new XL-MS experiments with other crosslinkers will likely resolve the question about the *bona fide* PEP subunit composition and the potential existence of stage-specific differences.

Indeed, the PEP envelope was calculated at very low resolution preventing fitting of the map with homologous structures of the catalytic core, or PAPs such as PAP9 (Favier *et al*., 2021), or PAP models with high confidence. However, the proposed fitting of the catalytic core of the *E. coli* RNAP (PDB entry: 3LU0 (Opalka *et al*., 2010)) reveals the remaining space for subsequent positioning of the PAPs (Figure S7). It is noteworthy that further 3D classifications did not reveal any significant variability in the 3D envelope of the PEP suggesting that the protrusions which we attribute to the PAPs are firmly associated with the catalytic core (Figures 5b-c). Despite the recognition of some structural features such as the cleft and stalk, the overall shape of the *S. alba* active PEP envelope is different from that of RNAPs II and III (Figures S8).

Sequence comparison (Figures S3-S6) shows that the four insertion regions characterized in the *E. coli* RNAP (Opalka *et al*., 2010) do not exist neither in PEP nor in the RNAP from *Nostoc*. The high sequence identity between the catalytic core of the bacteria and plastids suggests that the overall shape of the PEP core and the associated catalytic activity are conserved. The bacterial β’ subunit has been likely split into two subunits during evolution after the separation of the eubacteria and cyanobacteria branches, the latest uniquely sharing the β’’ subunit with the chloroplast (Schneider & Hasekorn, 1988). Sequence alignment showed that the β’ and β’’ subunits of the PEP can be respectively aligned with the N-terminal and C-terminal part of the β’ subunit from bRNAPs. In addition, a very long insertion in the β’’ subunit of plastids and cyanobacteria (Phe364-Ser1093 in *A. thaliana*) is not observed in the C-terminal part of the β’ subunit from bRNAPs. This insertion is located in the trigger loop region at the surface of the bRNAPs (Figure S6). With such a length, this region could be an additional domain in the PEP, associated with oxygenic photosynthesis.

Sequence divergence with the *T. thermophilus* and *E. coli* RNAPs are mainly observed between residues located at the surface of the core complex. Since the nuclear-encoded PAPs seems to have appeared with the terrestrialization of the green lineage (1^st^ appearance in fresh water algae and mosses), it is likely that the evolution of novel cell types requested some control of the PEP catalytic core activity, providing the capacity to generate novel plastid types. The PAPs, acting as signaling components expressed after the phytochrome activation in nucleus of angiosperms, may have been required to control PEP activity by the nucleus in order to synchronize the transcription of the photosynthesis associated nuclear genes (*PhANGs*) and photosynthesis associated plastid genes (*PhAPGs*) for the proper building of the photosynthetic apparatus upon first illumination. Because of their dual localization some of the PAPs such as PAP5/HEMERA (Nevarez *et al*., 2017) and PAP8 (Liebers *et al*., 2020) provide a potential regulatory link between nucleus and plastids in the expression of photosynthesis genes. It remains to be solved whether their nuclear or their plastid function evolved first.

## Material and methods

### Chloroplast isolation

6 to 7-day-old *Sinapis alba* cotyledons were collected and homogenized using a blender with short pulses (3 × 3 s); 100 g approximately of fresh material in 200 mL homogenization buffer containing 50 mM HEPES-KOH pH 8.0, 0.3 M sorbitol, 5 mM MgCl_2_, 2 mM EDTA and 0.3 mM DTT. The suspension obtained was then filtered through muslin and centrifuged 3 min at 6084.1*g* at 4°C. The pellet was recovered, resuspended in homogenization buffer and poured in a potter to remove all the chloroplast aggregates. The suspension was then loaded on a linear percoll gradient (35 % percoll, 50 mM HEPES-KOH pH 8.0, 0.3 M sorbitol, 5 mM MgCl_2_, 2 mM EDTA and 0.3 mM DTT) and centrifuged 50 min at 4696*g*, 4°C. The fractions containing the chloroplasts were then pooled, diluted in homogenization buffer and centrifuged 10 min at 4000*g*, 4°C to remove percoll. The pellet containing the chloroplasts was solubilized in the lysis buffer containing 50 mM TrisHCl pH 7.6, 25 % glycerol (w/v), 10 mM NaF, 4 mM EDTA, 1 mM DTT, 1 % Triton (w/v) and poured in potter for homogenization. The suspension was then centrifuged 1 h, at 15000*g*, 4°C and the supernatant frozen in liquid nitrogen and stored at -80°C before using to purify the PEP.

### PEP purification

After thawing, the chloroplast lysate was mixed overnight at 4°C with heparin resin equilibrated with 50 mM HEPES pH 7.6, 10 % (w/v) glycerol, 10 mM MgCl_2_, 80 mM (NH_4_)_2_SO_4_, 1 mM DTT, 0,1 % (w/v) Triton. The resin was extensively washed with 50 mM HEPES pH 7.6, 10 % (w/v) glycerol, 10 mM MgCl_2_, 80 mM (NH_4_)_2_SO_4_, 1 mM DTT, 0.1 % Triton (w/v) before elution over 10 fractions of 1 mL with 50 mM HEPES pH 7.6, 10 % (w/v) glycerol, 10 mM MgCl_2_, 1.2 M (NH_4_)_2_SO_4_, 1 mM DTT, 0.1 % Triton (w/v). The fractions were then subjected to SDS-PAGE and Western-Blot analyses with anti-PAP8 antibodies (Liebers *et al*., 2020). The fractions containing PAP8, and therefore the PEP, were pooled, loaded on a 35%-15% glycerol gradient (50 mM HEPES pH 7.6, 35 to 15 % (w/v) glycerol, 10 mM MgCl_2_, 0.01 % (w/v) Triton), and centrifuged at 97083*g* on a SW55-Ti rotor (Beckmann Coulter) during 16 h at 4°C.

The gradient was then analyzed using SDS-PAGE and Western-Blot. The fractions containing the PEP were pooled before the last step of purification, or frozen in liquid nitrogen and stored at - 80°C. The pool containing the PEP was mixed overnight with Q-sepharose resin (Amersham) pre-equilibrated in 50 mM HEPES pH 7.6, 10 % glycerol (w/v), 10 mM MgCl_2_, 0.01 % (w/v) Triton. The complex was eluted using a 0-1 M NaCl gradient. The fractions containing the PEP were pooled and concentrated at 2,000*g* on a 100-kDa cutoff membrane. The purified PEP was then frozen in liquid nitrogen and kept at -80°C before analyses.

### Sequence alignments

Full-length coding sequences of the α, β, β’ and β’’ subunits were retrieved from Blastp. The protein sequences were aligned using Clustal Omega (https://www.ebi.ac.uk/Tools/msa/clustalo/) and then colored using BOXSHADE server using default parameters. The domains of the α, β, β’ and β’’ subunits of the PEP were assigned based on those described (Lane & Darst, 2010; Sutherland & Murakami, 2018).

### MS-based proteomic analyses

Three PEP preparations from independently grown plant batches were analyzed. For this, purified PEP from chloroplasts was solubilized in Laemmli buffer and stacked in the top of a 4-12 % NuPAGE gel (Invitrogen). After staining with R-250 Coomassie Blue (Biorad), proteins were digested in-gel using trypsin (modified, sequencing purity, Promega), as previously described (Casabona *et al*., 2013). The resulting peptides were analyzed by online nanoliquid chromatography coupled to MS/MS (Ultimate 3000 RSLCnano and Q-Exactive Plus, Thermo Fisher Scientific) using a 140-min gradient. For this purpose, the peptides were sampled on a precolumn (300 μm × 5 mm PepMap C18, Thermo Scientific) and separated in a 75 μm × 250 mm C18 column (Reprosil-Pur 120 C18-AQ, 1.9 μm, Dr. Maisch). The MS and MS/MS data were acquired by Xcalibur (Thermo Fisher Scientific). Peptides and proteins were identified by Mascot (version 2.7, Matrix Science) through concomitant searches against the NCBI database (*Sinapis alba* strain: S2 GC0560-79 (white mustard) taxonomy, BioProject PRJNA214277, July 2020 download), the Uniprot database (*Sinapis alba* taxonomy, February 2021 download), a homemade database containing the sequences of classical contaminant proteins found in proteomic analyses (human keratins, trypsin, *etc*.), and the corresponding reversed databases. Trypsin/P was chosen as the enzyme and two missed cleavages were allowed. Precursor and fragment mass error tolerances were set at respectively at 10 and 20 ppm. Peptide modifications allowed during the search were: Carbamidomethyl (C, fixed), Acetyl (Protein N-term, variable) and Oxidation (M, variable). The Proline software (Bouyssié *et al*., 2020) was used for the compilation, grouping, and filtering of the results (conservation of rank 1 peptides, peptide length ≥ 6 amino acids, peptide score ≥ 25, allowing to reach a false discovery rate of peptide-spectrum-match identifications < 1% as calculated on peptide-spectrum-match scores by employing the reverse database strategy, and minimum of one specific peptide per identified protein group). Proline was then used to perform a MS1 label-free quantification of the identified protein groups based on razor and specific peptides. Intensity-based absolute quantification (iBAQ, Schwanhäusser *et al*., 2011) values were calculated from MS1 intensities of razor and specific peptides. The iBAQ values of each protein were normalized by the sum of iBAQ values of all quantified proteins in each sample, before summing the values of the three replicates to generate the final iBAQ value. The gene names for the identified proteins were annotated after Blastp search against *A. thaliana* proteome.

### Crosslinking coupled to MS analyses

A few micrograms of two PEP preparations used for mass spectrometry-based proteomic analyses (replicates 2 and 3) were cross-linked during 1h at room temperature using 100 µM of DSBU in HEPES buffer pH 7.8. To quench the cross-linking reaction, one μL of 1 M ammonium bicarbonate was added and the sample incubated for 15 min at room temperature. To reduce disulfide bonds, 100 mM DTT solution was added to obtain a final concentration of 3.5 mM and the mixture was incubated at 56°C for 30 min in a ThermoMixer. For alkylation of cysteines, 50 mM IAA solution was added to a final concentration of 8 mM and the mixture was incubated at room temperature in the dark for 20 min. Freshly prepared trypsin solution to an enzyme/protein ratio of ∼1:50 was added and the digestion was performed overnight at 37°C. To quench the enzymatic digestion, a final TFA concentration of 1% (v/v) was added. Micro spin columns (Harvard Apparatus) were then used to desalt the samples using 5 % ACN, 0.1 % TFA as washing solution and 75 % ACN, 0.1 % TFA as elution buffer.

The resulting peptides were analyzed by online nanoliquid chromatography coupled to MS/MS (Ultimate 3000 RSLCnano and Orbitrap Exploris 480 for replicate 2, and Q-Exactive HF for replicate 3, Thermo Fisher Scientific). Peptides were sampled on a precolumn (300 μm × 5 mm PepMap C18, Thermo Scientific) and separated using a Pharmafluidics μPAC™ column of 200 cm length (with pillar array backbone at interpillar distance of 2.5 μm) using a 240 min method. Data were acquired in data-dependent MS/MS mode with stepped higher-energy collision induced dissociation (HCD) and normalized collision energies (20 %, 25 %, 35 % for Orbitrap Exploris 480, and 22 %, 27 %, 30 % for Q-Exactive HF).

Data analysis was conducted using MeroX 2.0 (Iacobucci *et al*., 2018). The following settings were applied: proteolytic cleavage: C-ter at Lys and Arg with 3 missed cleavages allowed, peptide length 4 to 30 amino acids, fixed modification: alkylation of Cys by IAA, variable modification: oxidation of Met, cross-linker: DSBU with specificity towards Lys, Ser, Thr, Tyr, N-ter for site 1 and 2, analysis mode: RISEUP mode, maximum missing ions: 2, precursor mass accuracy: 10 ppm, product ion mass accuracy: 30 ppm, signal-to-noise ratio: 2, precursor mass correction activated, pre-score cutoff at 10 % intensity, FDR cut-off: 1 %, and minimum score cut-off: 30. Cross-links identified in the two replicates were then combined using Merox.

### Negative staining electron microscopy

10 µL of PEP was added to a glow discharge grid coated with a carbon supporting film for 3 minutes and the grid was stained with 50 µL of Sodium Silico Tungstate (SST) (1 % (w/v) in distilled water (pH 7-7.5)) for 2 minutes. The excess solution was soaked off by a filter paper and the grid was air-dried. The images were taken at 30,000 magnification (2.2 Å/pixel) under low dose conditions (<10 e-/Å2) with defocus values between -1.2 and -2.5 μm on a Tecnai 12 (Thermo Fischer Scientific) LaB6 electron microscope operating at 120 kV using a Gatan Orius 1000 CCD camera.

### Determination of the PEP envelope

The image processing was entirely done in RELION (Scheres, 2012). The CTF parameters of each micrograph was determined with CTFFIND4 (Rohou & Grigorieff, 2015) and the particles were auto-picked in RELION with the Laplacian of Gaussian option. 2D classification was then performed in 50 classes using a 350 Å mask diameter that resulted in the selection of 17,567 particles. The latter were then used to create an *ab initio* model (C1 symmetry and 300 Å mask diameter) which was then used to calculate a 3D map (C1 symmetry and 320 Å mask diameter) at 27.5 Å resolution (at FSC = 0.143).Acknowledgments

IBS acknowledges integration into the Interdisciplinary Research Institute of Grenoble (IRIG, CEA). This work used the platforms of the Grenoble Instruct-ERIC center (ISBG; UAR 3518 CNRS-CEA-UGA-EMBL) within the Grenoble Partnership for Structural Biology (PSB), supported by FRISBI (ANR-10-INBS-0005-02) and GRAL, financed within the University Grenoble Alpes graduate school (Ecoles Universitaires de Recherche) CBH-EUR-GS (ANR-17-EURE-0003).

### MS and EM data

The MS data have been deposited to the ProteomeXchange Consortium via the PRIDE (Perez-Riverol *et al*., 2019) partner repository with the dataset identifier PXD032738.

The data of crosslinking coupled to MS analyses have been deposited to the ProteomeXchange Consortium *via* the PRIDE (Perez-Riverol *et al*., 2019) partner repository with the dataset identifier PXD032739. Each spectrum corresponding to inter-protein link with the best scores were manually checked. The EM data have been deposited into the EMDB with the accession code EMD-14571.

## Supporting information

Data sources

## Author contributions

DC and RB designed the research. RR, FXG, YC, SKJ, SSM DF, GE, RB, and DC performed the research. YC and SKJ contributed MS data. DF and GE contributed EM data. DC and RB wrote the manuscript with contributions from RR, FXG, YC, SKJ, DF, GE and TP. All authors approved the manuscript.

## Funding

This work was supported by the Agence National de la Recherche (ANR-17-CE11-0031). The proteomic experiments were partly supported by ProFI (ANR-10-INBS-08-01 grant).

## Conflict of interest

The authors declare that the research was conducted in the absence of any commercial or financial relationships that could be construed as a potential conflict of interest.

## Notes

### Competing Interest Statement

The authors have declared no competing interest.

## References

Armstrong, G.A. (1998). Greening in the dark: light-independent chlorophyll biosynthesis from anoxygenic photosynthetic bacteria to gymnosperms. J. Photochem. Photobiol. B, 43, 87–100. doi: 10.1016/j.tplants.2010.07.002

Arsova, B., Hoja, U., Wimmelbacher, M., Greiner, E., Ustün, S., Melzer, M. et al. (2010). Plastidial thioredoxin z interacts with two fructokinase-like proteins in a thiol-dependent manner: evidence for an essential role in chloroplast development in Arabidopsis and Nicotiana benthamiana. Plant Cell, 22, 1498–1515. doi: 10.1105/tpc.109.071001

Bobik, K. & Burch-Smith, T.M. (2015). Chloroplast signaling within, between and beyond cells. Front. Plant Sci., 6, 781. doi: 10.3389/fpls.2015.00781

Bouyssié, D., Hesse, A.M., Mouton-Barbosa, E., Rompais, M., Macron, C., Carapito, C. et al. (2020). Proline: an efficient and user-friendly software suite for large-scale proteomics. Bioinformatics, 36, 3148–3155. doi: 10.1093/bioinformatics/btaa118

Casabona, M.G., Vandenbrouck, Y., Attree, I. & Couté, Y. (2013). Proteomic characterization of Pseudomonas aeruginosa PAO1 inner membrane. Proteomics, 13, 2419–2423. doi: 10.1002/pmic.201200565

Chen, M., Galvão, R.M., Li, M., Burger, B., Bugea, J., Bolado, J. et al. (2010). Arabidopsis HEMERA/pTAC12 initiates photomorphogenesis by phytochromes. Cell, 141, 1230-1240. 10.1016/j.cell.2010.05.007

Cramer, P. (2002). Multisubunit RNA polymerases. Curr. Opin. Struct. Biol., 12, 89–97. doi: 10.1016/s0959-440x(02)00294-4

Favier, A., Gans, P., Boeri Erba, E., Signor, L., Muthukumar, S. S., Pfannschmidt, T. et al. (2021). The plastid-encoded RNA polymerase-associated protein PAP9 is a superoxide dismutase with unusual structural features. Front. Plant. Sci., 12, 668897. doi: 10.3389/fpls.2021.668897

Finet, C., Timme, R.E., Delwiche, C.F., Marlétaz, F. (2010). Multigene phylogeny of the green lineage reveals the origin and diversification of land plants. Curr. Biol., 20, 2217–22. doi: 10.1016/j.cub.2010.11.035

Garcia, M., Myouga, F., Takechi, K., Sato, H., Nabeshima, K., Nagata, N. et al. (2008). An Arabidopsis homolog of the bacterial peptidoglycan synthesis enzyme MurE has an essential role in chloroplast development. Plant J., 53, 924–934. doi: 10.1111/j.1365-313X.2007.03379.x

Gao, Z.P., Yu, Q.B., Zhao, T.T., Ma, Q., Chen, G.X. & Yang, Z.N. (2011). A functional component of the transcriptionally active chromosome complex, Arabidopsis pTAC14, interacts with pTAC12/HEMERA and regulates plastid gene expression. Plant Physiol., 157, 1733–1745. doi: 10.1104/pp.111.184762

Gilkerson, J., Perez-Ruiz, J.M., Chory, J. & Callis, J. (2012). The plastid-localized pfkB-type carbohydrate kinases FRUCTOKINASE-LIKE 1 and 2 are essential for growth and development of Arabidopsis thaliana. BMC Plant Biol., 12, 102. doi: 10.1186/1471-2229-12-102

Glyde, R., Ye, F., Jovanovic, M., Kotta-Loizou, I., Buck, M., & Zhang, X. (2018). Structures of bacterial RNA polymerase complexes reveal the mechanism of DNA loading and transcription initiation. Mol. Cell, 70, 1111–1120.e3. doi: 10.1016/j.molcel.2018.05.021

Hajdukiewicz, P.T., Allison, L.A. & Maliga, P. (1997). The two RNA polymerases encoded by the nuclear and the plastid compartments transcribe distinct groups of genes in tobacco plastids. EMBO J., 16, 4041–4048. doi: 10.1093/emboj/16.13.4041

Hanske, J., Sadian, Y. & Müller C.W. (2018). The cryo-EM resolution revolution and transcription complexes. Curr. Opin. Struct. Biol., 52, 8–15. doi: 10.1016/j.sbi.2018.07.002

Hills, A.C., Khan, S. & López-Juez E. (2015). Chloroplast biogenesis-associated nuclear genes: control by plastid signals evolved prior to their regulation as part of photomorphogenesis. Front. Plant Sci., 6, 1078. doi: 10.3389/fpls.2015.01078

Hirata, A., Klein, B.J. & Murakami, K.S. (2008). The X-ray crystal structure of RNA polymerase from Archaea. Nature, 451, 851–854. doi: 10.1038/nature06530

Iacobucci, C., Götze, M., Ihling, C., Piotrowski, C., Arlt, C., Schäfer, M. et al. (2018). A cross-linking/mass spectrometry workflow based on MS-cleavable cross-linkers and the MeroX software for studying protein structures and protein-protein interactions. Nat. Protoc., 13, 2864–2889. doi: 10.1038/s41596-018-0068-8

Jumper, J., Evans, R., Pritzel, A., Green, T., Figurnov, M., Ronneberger, O.j et al. (2021). Highly accurate protein structure prediction with AlphaFold. Nature, 596, 583–589. doi: 10.1038/s41586-021-03819-2

Kassube, S.A., Fang, J., Grob, P., Yakovchuk, P., Goodrich, J.A., & Nogales, E. (2013). Structural insights into transcriptional repression by noncoding RNAs that bind to human Pol II. J. Mol. Biol., 425, 3639–3648. doi: 10.1016/j.jmb.2012.08.024

Lee, J. & Borukhov, S. (2016). Bacterial RNA polymerase-DNA interaction-the driving force of gene expression and the target for drug action. Front. Mol. Biosci., 3, 73. doi: 10.3389/fmolb.2016.00073

Liebers, M., Chevalier, F., Blanvillain, R. & Pfannschmidt, T. (2018). PAP genes are tissue- and cell-specific markers of chloroplast development. Planta, 248, 629–646. doi: 10.1007/s00425-018-2924-8

Liebers, M., Gillet, F.X., Israel, A., Pounot, K., Chambon, L., Chieb, M. et al. (2020). Nucleo-plastidic PAP8/pTAC6 couples chloroplast formation with photomorphogenesis. EMBO J., 39:e104941. doi: 10.15252/embj.2020104941

Lin, W., Das, K., Degen, D., Mazumder, A., Duchi, D., Wang, D., Ebright, Y. W. et al. (2018). Structural basis of transcription inhibition by fidaxomicin (Lipiarmycin A3). Mol. Cell, 70, 60-71.e15. doi: 10.1016/j.molcel.2018.02.026

Majeran, W., Friso, G., Asakura, Y., Qu, X., Huang, M., Ponnala, L. et al. (2012). Nucleoid-enriched proteomes in developing plastids and chloroplasts from maize leaves: a new conceptual framework for nucleoid functions. Plant Physiol., 158, 156–189. doi: 10.1104/pp.111.188474

Martin, W., Rujan, T., Richly, E., Hansen, A., Cornelsen, S., Lins, T. et al. (2002). Evolutionary analysis of Arabidopsis, cyanobacterial, and chloroplast genomes reveals plastid phylogeny and thousands of cyanobacterial genes in the nucleus. Proc. Natl. Acad. Sci. USA, 99, 12246–12251. doi: 10.1073/pnas.182432999

Myouga, F., Hosoda, C., Umezawa, T., Iizumi, H., Kuromori, T., Motohashi, R. et al. (2008). A heterocomplex of iron superoxide dismutases defends chloroplast nucleoids against oxidative stress and is essential for chloroplast development in Arabidopsis. Plant Cell, 20, 3148–3162. doi: 10.1105/tpc.108.061341

Murakami, K.S. (2015). Structural biology of bacterial RNA polymerase. Biomolecules, 5, 848–864. doi: 10.3390/biom5020848

Nevarez, P.A., Qiu, Y., Inoue, H., Yoo, C. Y., Benfey, P. N., Schnell, D.J. et al. (2017). Mechanism of dual targeting of the phytochrome signaling component HEMERA/pTAC12 to plastids and the nucleus. Plant Physiol., 173, 1953–1966. doi: 10.1104/pp.16.00116

Opalka N., Brown J., Lane W.J., Twist K.A., Landick R., Asturias F.J. et al. (2010). Complete structural model of Escherichia coli RNA polymerase from a hybrid approach. PloS Biol., 8, e1000483. doi: 10.1371/journal.pbio.1000483

Perez-Riverol, Y., Csordas, A., Bai, J., Bernal-Llinares, M., Hewapathirana, S., Kundu, D.J. et al. (2019). The PRIDE database and related tools and resources in 2019: improving support for quantification data. Nucleic Acids Res., 47, D442–D450. doi: 10.1093/nar/gky1106

Pettersen, E.F., Goddard, T.D., Huang, C.C., Couch, G.S., Greenblatt, D.M., Meng, E.C. et al. (2004). UCSF Chimera--a visualization system for exploratory research and analysis. J. Comput. Chem., 25, 1605–1612. doi: 10.1002/jcc.20084

Pfalz, J., Liere, K., Kandlbinder, A., Dietz, K.J. & Oelmüller, R. (2006). pTAC2, -6, and -12 are components of the transcriptionally active plastid chromosome that are required for plastid gene expression. Plant Cell, 18, 176–197. doi: 10.1105/tpc.105.036392

Pfalz, J., & Pfannschmidt, T. (2013). Essential nucleoid proteins in early chloroplast development. Trends Plant Sci., 18, 186–194. doi: 10.1016/j.tplants.2012.11.003

Pfannschmidt, T. & Link, G. (1994). Separation of two classes of plastid DNA-dependent RNA polymerases that are differentially expressed in mustard (Sinapis alba L.) seedlings. Plant Mol. Biol., 25, 69–81. doi: 10.1007/BF00024199

Pfannschmidt, T., Ogrzewalla, K., Baginsky, S., Sickmann, A., Meyer, H.E. & Link, G. (2000). The multisubunit chloroplast RNA polymerase A from mustard (Sinapis alba L.). Integration of a prokaryotic core into a larger complex with organelle-specific functions. Eur. J. Biochem., 267, 253–261. doi: 10.1046/j.1432-1327.2000.00991.x

Pfannschmidt, T., Blanvillain, R., Merendino, L., Courtois, F., Chevalier, F., Liebers, M. et al. (2015). Plastid RNA polymerases: orchestration of enzymes with different evolutionary origins controls chloroplast biogenesis during the plant life cycle. J. Exp. Bot., 66, 6957–6973. doi: 10.1093/jxb/erv415

Riggs, J.W. & Callis, J. (2017). Arabidopsis fructokinase-like protein associations are regulated by ATP. Biochem. J., 474, 1789–1801. doi: 10.1042/BCJ20161077

Rohou, A. & Grigorieff, N. (2015). CTFFIND4: fast and accurate defocus estimation from electron micrographs. J. Struc. Biol., 192, 216–221. doi: 10.1016/j.jsb.2015.08.008

Scheres, S.H.W. (2012). RELION: implementation of a bayesian approach to cryo-EM structure determination. J. Struct. Biol., 180, 519–530. doi: 10.1016/j.jsb.2015.08.008

Schneider, G.J. & Hasekorn, R. (1988). RNA polymerase subunit homology among cyanobacteria, other eubacteria and archaebacteria. J. Bacteriol., 170, 4136–4140. doi: 10.1128/jb.170.9.4136-4140.1988

Schwanhäusser, B., Busse, D., Li, N., Dittmar, G., Schuchhardt, J., Wolf J. et al. (2011). Global quantification of mammalian gene expression control. Nature, 473, 337–342. doi: 10.1038/nature10098

Steiner, S., Schröter, Y., Pfalz, J. & Pfannschmidt, T. (2011). Identification of essential subunits in the plastid-encoded RNA polymerase complex reveals building blocks for proper plastid development. Plant Physiol., 157, 1043–1055. doi: 10.1104/pp.111.184515

Sutherland, C. & Murakami, K.S. (2018). An introduction to the structure and function of the catalytic core enzyme of Escherichia coli RNA polymerase. EcoSal Plus, 8, 10.1128 doi: 10.1128/ecosalplus.ESP-0004-2018

Sugiura, M. (1992). The chloroplast genome. Plant Mol. Biol., 19, 149–168. doi: 10.1007/BF00015612

Suzuki, J.Y., Ytterberg, A.J., Beardslee, T.A., Allison, L.A., Wijk, K.J. & Maliga, P. (2004). Affinity purification of the tobacco plastid RNA polymerase and in vitro reconstitution of the holoenzyme. Plant J., 40, 164–172. doi: 10.1111/j.1365-313X.2004.02195.x

Vannini, A., Ringel, R., Kusser, A. G., Berninghausen, O., Kassavetis, G. A., & Cramer, P. (2010). (2010). Molecular basis of RNA polymerase III transcription repression by Maf1. Cell, 143, 59–70. doi: 10.1016/j.cell.2010.09.002

Weihe, A. & Börner, T. (1999). Transcription and the architecture of promoters in chloroplasts. Trends Plant Sci., 4, 169–170. doi: 10.1016/s1360-1385(99)01407-7

Williams-Carrier, R., Zoschke, R., Belcher, S., Pfalz, J. & Barkan, A. (2014). A major role for the plastid-encoded RNA polymerase complex in the expression of plastid transfer RNAs. Plant Physiol., 164, 239–248. doi: 10.1104/pp.113.228726

Yagi, Y., Ishizaki, Y., Nakahira, Y., Tozawa, Y. & Shiina, T. (2012). Eukaryotic-type plastid nucleoid protein pTAC3 is essential for transcription by the bacterial-type plastid RNA polymerase. Proc. Natl. Acad. Sci. U S A, 109, 7541–7546. doi: 10.1073/pnas.1119403109

Yagi, Y. & Shiina, T. (2014). Recent advances in the study of chloroplast gene expression and its evolution. Front. Plant Sci., 5, 61. doi: 10.3389/fpls.2014.00061

Yu, Q.B., Lu, Y., Ma, Q., Zhao, T.T., Huang, C., Zhao, H.F. et al. (2013). TAC7, an essential component of the plastid transcriptionally active chromosome complex, interacts with FLN1, TAC10, TAC12 and TAC14 to regulate chloroplast gene expression in Arabidopsis thaliana. Physiol. Plant., 148, 408–421. doi: 10.1111/j.1399-3054.2012.01718.x

Yu, Q.B., Huang, C. & Yang, Z.N. (2014). Nuclear-encoded factors associated with the chloroplast transcription machinery of higher plants. Front. Plant Sci., 5, 316. doi: 10.3389/fpls.2014.00316

Yua, Q.B., Ma, Q., Kong, M.M., Zhao, T.T., Zhang, X.L., Zhou, Q. et al. (2014). AtECB1/MRL7, a thioredoxin-like fold protein with disulfide reductase activity, regulates chloroplast gene expression and chloroplast biogenesis in Arabidopsis thaliana. Mol. Plant, 7, 206–217. doi: 10.1093/mp/sst092

Zhelyazkova, P., Sharma, C.M., Förstner, K.U., Liere, K., Vogel, J. & Börner, T. (2012). The primary transcriptome of barley chloroplasts: numerous noncoding RNAs and the dominating role of the plastid-encoded RNA polymerase. Plant Cell, 24, 123–136. doi: 10.1105/tpc.111.089441

Zybailov, B., Rutschow, H., Friso, G., Rudella, A., Emanuelsson, O., Sun, Q. et al. (2008). Sorting signals, N-terminal modifications and abundance of the chloroplast proteome. Plos One, 3, e1994. doi: 10.1371/journal.pone.0001994

