## Supplemental Figures for "Three-dimensional envelope and subunit interactions of the plastid-encoded RNA polymerase from *Sinapis alba*"

***Supplemental tables***

**Table S1:** MS-based proteomic characterization of *S. alba* PEP fraction.

**Table S2:** characterization of proximal proteins in *S. alba* PEP fraction using crosslinking-MS.

***Supplemental figures***

**Figure S1:** dependency of plastome transcription units on NEP and PEP activities. The plastome of *Arabidopsis thaliana* is depicted as circular map including large and small single copy regions (LSC and SSC, respectively), as well as the inverted repeats A and B (IRA and IRB). Circles indicate preferential transcription by NEP or PEP.

**Figure S2:** abundance-based ranking of proteins quantified by MS in PEP enriched samples. Distribution of abundances represented as Log2 of normalized and summed iBAQ values of individual proteins detected in three independent PEP samples (Table S1). Identified subunits of individual complexes are color-coded (blue: PAPs; orange: α, β, β´ and β´´ subunits; green: histones). The annotated zoom-in shows the 24 most abundant proteins in the ranking.

**Figure S3-6:** sequence alignment of the α, β, β´ and β´´ subunits from PEP of angiosperms with those of the RNAPs from *E. coli*, *T. thermophilus* and Nostoc. S3) Sequence alignment of the α subunits, S2) sequence alignment of the β subunits S3) Sequence alignment of the N-terminal part from β´ subunit with β ´ subunit from PEP, S4) Sequence alignment of the C-terminal part from β´ subunit with the β´´ subunit from PEP. The residues conserved more than 50 % are in red, those mutated in similar residues are in blue. The strictly conserved residues described by Lane & Darst (Lane & Darst, 2010) are highlighted in gray. The blue triangles show mutations observed among the strictly conserved residues described (Lane & Darst, 2010). The non-conservative mutations, at least three in a row in the β or β´ domain in *E. coli* and *T. thermophilus*, are high-lighted in green and displayed on the *E. coli* structure (PDB entry: 6GH5). Those colored in orange are nearby to the DNA, those in green are located at the surface of the subunits. The domains described for all-RNA polymerase (a) and the bRNAPs (b) are also given and highlighted in yellow and cyan respectively. The name of the RNAP domains are also given and highlighted in purple and green (Lane & Darst, 2010; Sutherland & Murakami, 2018).

**Figure S7:** view of the catalytic core from the *E. coli* RNAP (PDB entry: 3LU0 (Opalka *et al.*, 2010)) manually fitted into the envelope of PEP using Chimera (Pettersen *et al.*, 2004).

**Figure S8a and S8b:** overall shape of the a) human RNA polymerase II (EMDB entry: EMD-2194; Kassube et al., 2013) and b) yeast RNA polymerase III (EMDB entry: EMD-1753; Vanini et al., 2010) solved at 25 and 21 Å respectively.

***Data source:*** rpos collection from the green lineage

**Fig. S1.** Dependency of plastome transcription units on NEP and PEP activities. The plastome of *Arabidopsis thaliana* is depicted as circular map including large and small single copy regions (LSC and SSC, respectively), as well as the inverted repeats A and B (IRA and IRB). Circles indicate preferential transcription by NEP or PEP.

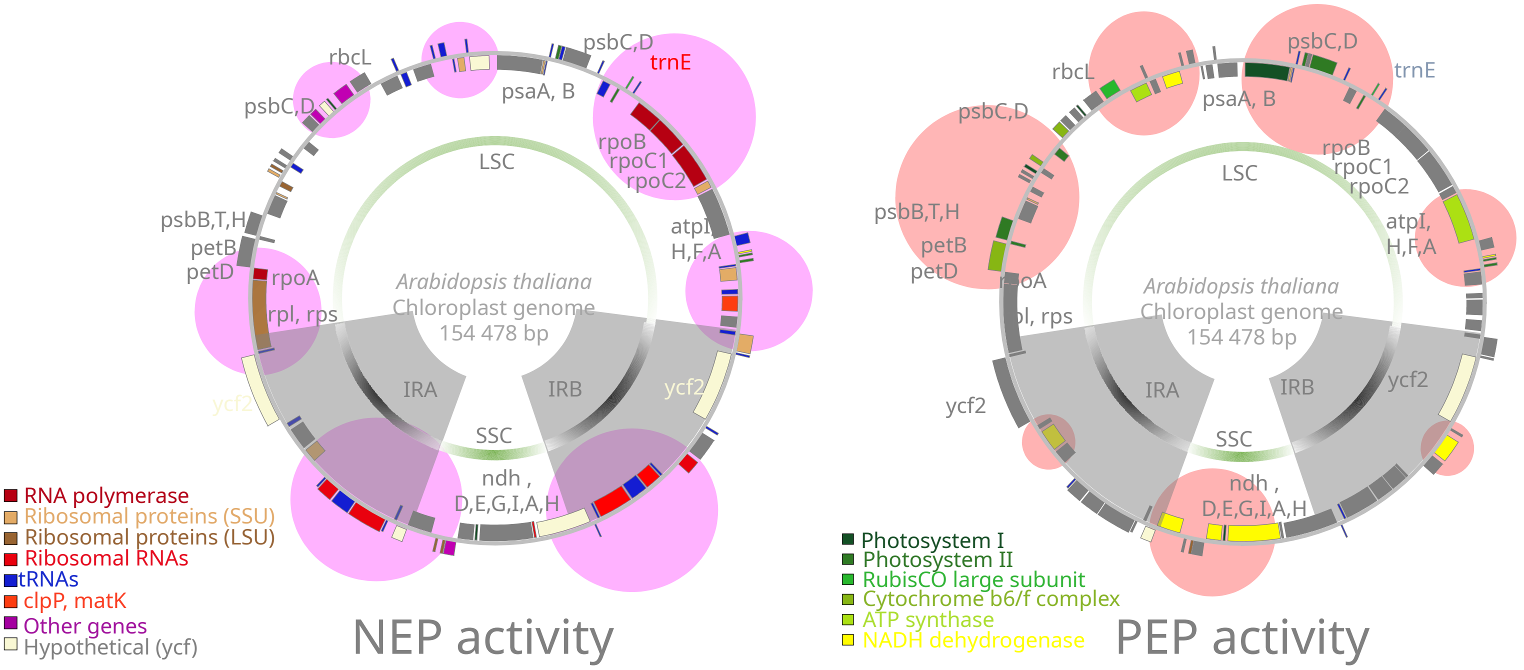

**Figure S2 :** abundance-based ranking of proteins quantified by MS in PEP enriched samples. Distribution of abundances represented as Log2 of normalized and summed iBAQ values of individual proteins detected in three independent PEP samples (Table S1). Identified subunits of individual complexes are color-coded (blue: PAPs; orange: α, β, β´ and β´´ subunits; magenta: histones). The annotated zoom-in shows the 24 most abundant proteins in the ranking.

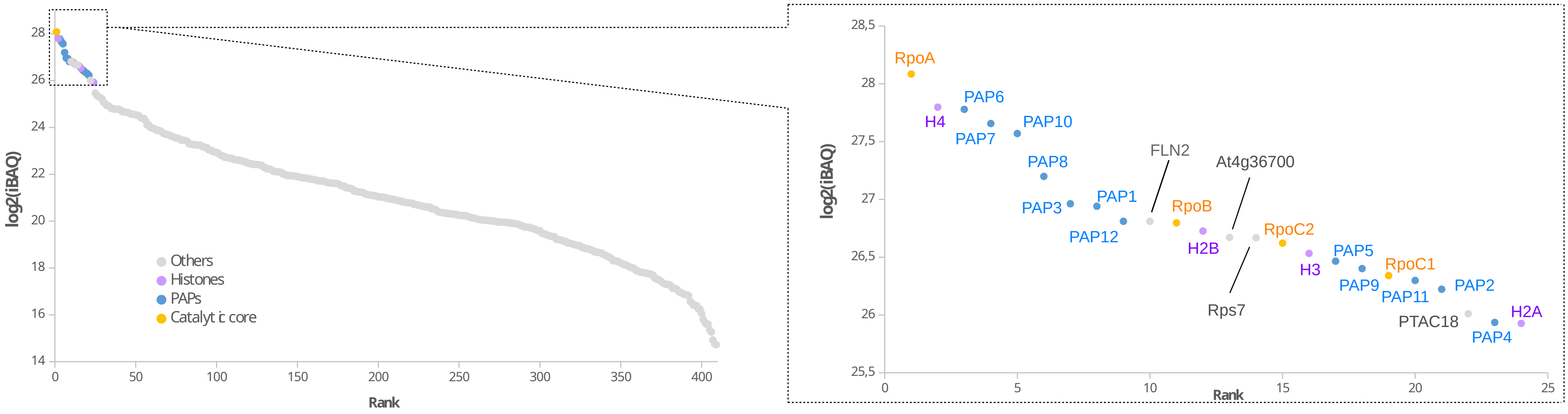

**Fig. S3.** Sequence alignment of the $\alpha$ from PEP of angiosperms with those of the RNAPs from *E. coli*, *T. thermophilus* and Nostoc. The residues conserved more than 50 % are in red, those mutated in similar residues are in blue. The strictly conserved residues described by Lane & Darst (Lane & Darst, 2010) are highlighted in gray. The blue triangles show mutations observed among the strictly conserved residues described (Lane & Darst, 2010). The non-conservative mutations, at least three in a row in the $\beta$ or $\beta$' domain in *E. coli* and *T. thermophilus*, are high-lighted in green and displayed on the *E. coli* structure (PDB entry: 6GH5). Those colored in orange are nearby to the DNA, those in green are located at the surface of the subunits. The domains described for all-RNA polymerase (a) and the bRNAPs (b) are also given and highlighted in yellow and cyan respectively. The name of the RNAP domains are also given and highlighted in purple and green (Lane & Darst, 2010; Sutherland & Murakami, 2018).

```
T. thermophilus     1 --------------------------------------------------------------------------------
E. coli             1 --------------------------------------------------------------------------------
0_Nostoc            1 --------------------------------------------------------------------------------
1_Litchi            1 --------------------------------------------------------------------------------
2_Arabidopsis       1 --------------------------------------------------------------------------------
3_Gossypium         1 --------------------------------------------------------------------------------
5_Ricinus           1 --------------------------------------------------------------------------------
6_Rosa              1 --------------------------------------------------------------------------------
9_Cucumis           1 --------------------------------------------------------------------------------
11_Nicotiana        1 --------------------------------------------------------------------------------
13_Syringa          1 --------------------------------------------------------------------------------
18_Liquidambar      1 --------------------------------------------------------------------------------
19_Papaver          1 --------------------------------------------------------------------------------
20_Ananas           1 --------------------------------------------------------------------------------
28_Liriodendron     1 --------------------------------------------------------------------------------
30_Magnolia         1 --------------------------------------------------------------------------------
32_Nymphaea         1 --------------------------------------------------------------------------------
33_Amborella        1 --------------------------------------------------------------------------------
35_Picea            1 --------------------------------------------------------------------------------
44_Ginkgo           1 --------------------------------------------------------------------------------
51_Physcomitrium    1 MAAVMSAQAMADAVRVVRASISELGKSAKKSDAFSARDLKSRSGFTESVDRLGLVNLQVQARKKRVRGLRLVPTRVVSAI
```

α-NTD

```
T. thermophilus     1 ------------------------------MLDSK-----------LKAPVFTVRTQGREYGEFVLEPLERGFGVTLGNP
E. coli             1 ---------------------------MQGSVTEF-----------LKPRLVDIEQVSSTHAKVTLEPLERGFGHTLGNA
0_Nostoc            1 -----------------------------------------MAQFQIECVESNTEESRNHYSKFVLEPLERGQGTTVGNA
1_Litchi            1 ------------------------------MVREKVK--VFTRTLQWKCVESRTESKRLYYGRFILSPLMKGQADTIGIA
2_Arabidopsis       1 ------------------------------MVREKVK--VSTRTLQWKCVESKRDSKRLYYGRFILSPLMKGQADTIGIA
3_Gossypium         1 ------------------------------MVREKVKVSTSTRTRQWKCVESRTDSKRLYYGRFILSPLMKGQADTIGIA
5_Ricinus           1 ------------------------------MIREKVT--ISTRTLQWKCVESRTDNKRLFYGRFILSPLMKGQADTIGIA
6_Rosa              1 ------------------------------MVREKVT--VSTRTLQWKCVESRADSKRLYYGRFILAPLMKGQADTIGIA
9_Cucumis           1 ----------------------MILSKNITMVREKIR--VSTRTLKWKCVESRADSKRLYYGRFILSPLMKGQGDTIGIA
11_Nicotiana        1 ------------------------------MVREKVT--VSTRTLQWKCVESRTDSKRLYYGRFILSPLMKGQADTIGIA
13_Syringa          1 ------------------------------MVREKVT--VSTRTLQWKCVESREDSKRLYYGRFILSPLMKGQADTIGIA
18_Liquidambar      1 ------------------------------MVREKVT--VSTRTLQWKCVESRADSKRLYYGRFILSPLMKGQADTIGIA
19_Papaver          1 ------------------------------MLREEVA--VSTRTLQWKCVESRADSKRLYYGRFILSPLLKGQADTIGIA
20_Ananas           1 ------------------------------MVREEVA--GYTRTLQWKCVESRVDSKRLYYGRFILSPLMKGQADTIGIA
28_Liriodendron     1 ------------------------------MVREEVA--VSTRTLQWKCVESRTDSKRLYYGRFILAPLMKGQADTIGIA
30_Magnolia         1 ------------------------------MVREEVA--VSTRTLQWKCVESRTDSKRLYYGRFILAPLMKGQADTIGIA
32_Nymphaea         1 ------------------------------MVREEVP--VSTRTLQWKCVESRADSKRLYYGRFVLSPLMKGQADTIGIA
33_Amborella        1 ------------------------------MVRKEVP--VSNRTLQWKCVESKADSKRLYYGRFVLSPLMKGQADTIGTA
35_Picea            1 ------------------------------MIRDEIS--VSIQTLRWKCIESRAYSERLHYGRFALSPLRKGRADTIGIA
44_Ginkgo           1 ------------------------------MIRDEIS--VSIQTLRWKCIESRVNGKRLHYGRFALSPLQKGQANTIGIA
51_Physcomitrium   81 EGSNSTTADAPVDEDVLAWTKAYRAENSTAITRDETLKSNAQSALQWKCVETQVEGERLHYGRFAVSPFRSGQANTVGVS
```

α-NTD

```
T. thermophilus    40 LRRILLSSIPGTAVTSVYIEDVLHEFSTIPGVKEDVVEIILNLKELVVRFLNPSLQTVTLLLKAEGPKEVKARDFLPVAD
E. coli            43 LRRILLSSMPGCAVTEVEIDGVLHEYSTKEGVQEDILEILLNLKGLAVRVQGKDEV--ILTLNKSGIGPVTAADITHDGD
0_Nostoc           40 LRRVLLSNLEGTAVTAVRIAGVSHEFATVPGVREDVLEILMRMKEVILKNYSSQPQ--IGRLLVNGPATITAAHFDLPSE
1_Litchi           49 MRRALLGEIEGTCITRAKFEKIPHECSTILGIQESVHEILMNLKEIVLRSNLYGTR--DALLCVKGPGYVTAQDILLPPS
2_Arabidopsis      49 MRRALLGEIEGTCITRAKSENIPHDYSNIAGIQESVHEILMNLNEIVLRSNLYGTR--NALICVQGPGYITARDIILPPA
3_Gossypium        51 MRRALLGELEGTCITRAKSEKIPHEYSTIVGIQESVHEILMNLKEIVLRGNLYGTR--NAFICAKGPGYVTAQDIILPPS
5_Ricinus          49 TRRALLGEIEGTCITRAKSEKIPHEFSTIAGIQESIHEILMNLKEIVLRSNLYGTC--DASICVKGPGYITAQDIILPPF
6_Rosa             49 MRRALLGEIEGTCITRARSEKIPHEYSTIVGIQESVHEILMNLKEIVLRSNLYGTR--NASICVKGPGYVTAQDIILPPS
9_Cucumis          57 MRKALLGEIEGTCITRAKSEKIPHEYSTIVGIQESVHEILMNLKEIVLRSNLYGTR--DASICVKGPGCVTAQDIILPPS
11_Nicotiana       49 MRRALLGEIEGTCITRVKSEKVPHEYSTITGIQESVHEILMNLKEIILRSNLYGTS--DASICVKGPGSVTAQDIILPPY
13_Syringa         49 MRRTLLGEIEGTCITRVKSENVPHEYSTIAGIQESVHEILMNLKEIVLRSNLYGTW--DASICVRGPGYVTAQDIILPPY
18_Liquidambar     49 MRRALLGEIEGTCITRAKSEKIPHEYSTIVGIQESVHEILMNLKEVVLRSNLYGTR--DASICVRGPGYVTAQDIISPPS
19_Papaver         49 MRRALLGEIEGTCITRAKSDKIPHEYSTIVGIEESVHEILMNLKEIVLRSNLYGTR--DASICVRGPGYVTAQDIISPPS
20_Ananas          49 MRRALLGEIEGTCITRAKSEKVPHEYSTIVGIEESVHEILLNLKEIVLRSNLYGVR--DALICVRGPRYVTAQDIISPPS
28_Liriodendron    49 MRRALLGEIEGTCITRVKSEKVPNEYYTIVGIEESVHEILMNLKEIVLRSHLYGTR--DASICVRGPRYVTAQDIIPPPS
30_Magnolia        49 MRRALLGEIEGTCITRAKSEKVPHEYSTIVGIEESVHAILMNLKEIVLGSHLYGTR--DASICVRGPRYVTAQDIIPPPS
32_Nymphaea        49 MRRALLGELEGTCITRAKSDKVPHEYSTIVGIEESVHEILMNLKKIVLRSDLYGTL--NASICVRGPRHVTAQDIISPPS
33_Amborella       49 MRRSLLGEIEGTCITCAKSERVPHEYSTIVGIEESVHEILMNLKEIIVRSDLYGTR--DAFICVRGPKYVTAQDIISPPS
35_Picea           49 MRRVLLGEVEGTCITHVKLENIKHEYSAIIGIEESVHDILMNLKEIVLRSDSYGIR--GASICIVGPRNVTAQDIILPPS
44_Ginkgo          49 MRRALLGEVEGTCITHAKFENMTHEYSAIMGIEESVHDISINLRGIVLQSDPYGIR--EASIYSVGPRDVTAQDIILPPS
51_Physcomitrium  161 MQKALLGEVEGAAVSCATFKNVKSEYAAMKGVEETPMDILVNLKELVIRSDSDEPQ--KAIISAIGPGPVTAGDIVLPPS
```

```
                                                            α-NTD
T. thermophilus   120 VEIMNPDLHIATL-EEGGRLNMEVRVDRGVGYVPAEKHGI----KDRINAIPVDAVFSPVRRVAFQVEDTRLGQRTDLDK
E. coli           121 VEIVKPQHVICHLTDENASISMRIKVQRGRGYVPASTRIHSEEDERPIGRLLVDACYSPVERIAYNVEAARVEQRTDLDK
0_Nostoc          118 VEVIDPTQYVATI-AEGGKLEMEFRIERGKGYRTVERGRE---EATSLDFLQIDSVFMPVRKVNYSVEEARGDGSITKDR
1_Litchi          127 VEIIDNTQHIASL-AEPIDFCIGLQIERNRGYNIKAPNNF---QD---RSYSIDAVFMPVRNANYSIHSYGSGN-EKQEI
2_Arabidopsis     127 VEIIDNTQHIATL-TEPIDLCIELKIERNRGYSLKMSNNF---ED---RSYPIDAVFMPVENANHSIHSYGNGN-EKQEI
3_Gossypium       129 VEIVDNTQHVASL-TEPIDLCIGLQIERNRGYGIKTPKNF---HD---GSYPIDAVFMPVRNANHSIHCYGNDN-EKQEI
5_Ricinus         127 VEIIDNTQHIASL-TEPIDLCIGLQIERNRGYRIKPTNNF---HE---GSYPIDAVFMPVRNANHSVHSYGNGN-EKQEI
6_Rosa            127 VEIVDNTQHIANL-TEPINLCIQLQIERNRGYRIKTPNNF---QD---GSYPIDAVFMPVRNANHSIHSYVNGN-EKQEI
9_Cucumis         135 VEIVDNTQHIANL-TEPINFCIELKIERNRGYHIQTPNNF---QD---ASYPMDAIFMPVRNVNHSIHSYVNGN-EKQEI
11_Nicotiana      127 VEIVDNTQHIASL-TEPIDFCIGLQIERNRGYLIKTPHNF---QD---GSYPIDAVFMPVRNANHSIHSYGNGN-EKQEI
13_Syringa        127 VEIVDNTQHIASL-TEPIDLCIGLQIERNRGYLIKMPDNF---QD---GSYPIDAVFMPVRNANHSIHSYGNGN-EKQEI
18_Liquidambar    127 VEIVDNTQHIASL-TEPIDLCIGLKMERNRGYRIKTPNNF---QD---RSYPIDAVFMPVRNANHSIHSYGNEN-EKQEI
19_Papaver        127 VEIVDTTQHIASL-TEPMDLCIGLQIERNRGYRMKTPNNA---QD---GSFTIDAVFMPVRNANHSIHSYGNGN-EKQEI
20_Ananas         127 VEIVDTTQHIANL-TEPIDLCIGLQIKRDRGYRMEPINDS---QD---VSYPIDAVSMPVRNANHSIHSYGNGN-EKQEI
28_Liriodendron   127 VEIVDTTQHIASL-TEPIDLCIELQIERDRGYRMKTPNNY---QD---GSYPIDAVSMPVRNANHSIHSYGNRN-EKQEI
30_Magnolia       127 VEIVDTTQHIASL-TEPIDLCIELQIERDRGYRMKTPNNY---QD---GSYPIDAVSMPVRNANHSIHSYGNGN-EKQEI
32_Nymphaea       127 VEIVDTTQHIAVL-TEPVDLCIELQIERGRGYCTRTPNNY---QD---GSYPIDAVSMPVRNANHSIHSYGNGN-EKQEI
33_Amborella      127 VKIVDTTQHIASI-TEPINFCIGLEIERNRGYRTRTTN-----QD---ASYPIDAVSMPVRNANYSIQCYGNEN-EKQEI
35_Picea          127 VKIIDTTQHIARL-TKSITSDIRLQIEKNRGYIIHSPNNY---QD---GIFPIDAVFMPVRDANYSIHSYGSGN-EIREV
44_Ginkgo         127 VRIIDTTQHIASL-KESITSDIRSQIGKDRGYRIQSPNNS---RD---GIFSTDAVSMPVRNANYSIHSYGNGN-EMQEI
51_Physcomitrium  239 LEVTDPTQHIAYL-TKEVSLDIELDVEKGCGYRMGDHTKS---GD---GRFYIDSVFMPVRNANYSVHSYESEPDVTQEI

                                        α-NTD                                          α-CTD
T. thermophilus   195 LTLRIWTDGSVTPLEALNQAVEILREHLTYFSNPQAAAVAAPEEAKE-----------------PEAPPEQEEELDLPL
E. coli           201 LVIEMETNGTIDPEEAIRRAATILAEQLEAFVDLRDVRQ--PE---V------------------KEEKPEFDPILLRPV
0_Nostoc          194 LLLEVWTNGSISPQEALSSAAGILVDLFNPLKDISLEPTD-------T-----------------GSEISEDPTAQIPI
1_Litchi          199 LFLEIWTNGGLTPKEALHEASRNLIDLFIPFLQAAE--ENLHLENNQYKVSLPYLTFRDRVAKLKKKK--KEIAGKSIFI
2_Arabidopsis     199 LFLEIWTNGSLTPKEALHQASRNLINLFIPFLHVEE--ETFYLENNQHQVTLPFFPFHNRLVNLRKKKKTKELAFQYIFI
3_Gossypium       201 LFLEIWTNGSLTPKEALHEASRNLIDLFIPFLHTEE--ENLHLENNQHDVTLPFFPFHDRLVKLTKKK--KEIALKYIFI
5_Ricinus         199 LFLEIWTNGSLTPKEALHEASRNLIDLFIPFLHAEE--ENFHLEKNQHKVTLPLFTFYDRLTKLRKNQ--NEITLKYVFI
6_Rosa            199 LFIEIWTNGSLTPKEALHEASRNLIDLFIPFLHAEE--ENFHFENNQHKVTLPLFTFHDKFANPRKSK--TEITLKYIFI
9_Cucumis         207 LFLEIWTNGSLTPKEALHEASRNLIDLFIPFLHAEEEKENFHFKNNKQKVTLPLFTFHEKLAKLRKKK--KERALKYIFI
11_Nicotiana      199 LFLEIWTNGSLTPKEALHEASRNLIDLFIPFLHMEE--DNLYLQDNQHTVPLSPFTFHDKLAKLIKNK--KKIALKSIFI
13_Syringa        199 LFLEIWTNGSLTPKEALHEASRNLIDLFIPFLHKEE--ENLPLEDNQHTVPLSPLTFHEKLDKLRKNK--KKIALKSIFI
18_Liquidambar    199 LFLEIWTNGSLTPKEALHEASRNLIDLFIPFLHAEE--ENLHLEDNQHKTILPLFTFHDRLAKLKKNK--KKKALKSIFI
19_Papaver        199 LFLEIWTNGSLTPKEALHEASRNLIDLFIPFLHAEERDTN--LEDNQNIVTIPFFTFQNKLDKL--SKSKKDIALKCIFI
20_Ananas         199 LFLEIWTNGSLTPKEALYEASRNLIDLFIPFLHAEEEDIN--FEENKNRFTLPFLTFQDRLTNL--KKNKKGIPLNCIFI
28_Liriodendron   199 LFLEIWTNGSLTPKEALHEASRNLIDLFIPFLHAEEQDIV--LEDNPNRFTVPLFTFHDRLANI--RKNKKGIALKCIFI
30_Magnolia       199 LFLEIWTNGSLTPKEALHEASRNLIDLFIPFLHAEEQDIN--LEDNPNRFTVPFFTFHDRLANI--RKNKKGIALKCIFI
32_Nymphaea       199 LFLEIWTNGSLTPKEALYEASRNLIDLFIPFLHAEEQDINRNMEDNLKRASVPFFTFDDGLDNM--K---REIILKRIFI
33_Amborella      197 LFLEIWTNGSLTPKEALYEASRNLIDLFIPFLHAEERDLH--LDDDQNIFTAPFFPFHEGLAYIGIGESKKRIALKRIFI
35_Picea          199 LFLEIWTNGGLTPREALSEASRILIELFIPFLHGEEQNI--DGMNNRKGSNMPPFPLSHVLTDTGET-KEKKIAFQHIFI
44_Ginkgo         199 LFSEIWTNGSLAPGEALYEASRNLIDLFIPFMRAEEQNI--DGIDNWNGYNMSLFPFSHISTDTEK--MEEEIAFKHIFI
51_Physcomitrium  312 LFLEIWTNGSITPEEALHEAARCLIDLFLPFLHPKKKEVTNSATKMHKSFTMSQFNSS-------AEMSAKEVDLRHVYV

                                                       α-CTD
T. thermophilus   257
EELGLSTRVLHSLKEEGIESVRALLALNLKDLKNIPGIGERSLEEIKEALEKKGFTLKE----------------
E. coli           258 DDLELTVRSANCLKAEAIHYIGDLVQRTEVELLKTPNLGKKSLTEIKDVLASRGLSLGMRLENW---PPASIADE
0_Nostoc          249 EELQLSVRAYNCLKRAQVNSVADLLDYTQEDLLEIKNFGQKSAEEVVEALQRRL-GITLPQERGSKHP-------
1_Litchi          275 DQPELSPRIYNCLKKSNIHTLFDLLNKSQDDLMKIEHFRIEDVKQILGILEKK----------------------
2_Arabidopsis     277 DQLELPPRIYNCLKKSNIHTLLDLLNNSQEDLIKIEHFHVEDVKKILDILEKK----------------------
3_Gossypium       277 DQSELPPRIYNCLKKSNIHTLLDLLNNSREDLMKIEHFRIEDVKQILGILEKK----------------------
5_Ricinus         275 DQSELTPKIYNCLKRSNIHTLSDLLNKSQEDLMKIEHFRIDDVKHILGILEIEK-NFAI----------------
6_Rosa            275 DQSELPPRVYNCLKRSNIHTLLDLLNNSQEDLIKMKHLRIEDVKHILNILEKKK-HFE-----------------
9_Cucumis         285 DQSELPPRIYNCLKRCNIHTLFDLLNNSPDELMKIKHFRIEDVKHILDILEMEK-NFA-----------------
11_Nicotiana      275 DQSELPSRIYNCLKMSNIYTLLDLLNNSQEDLMKIEHFRSEDVKRILGILEKYF-VIDLAKNKF-----------
13_Syringa        275 DQSEFSPRVYNCLKRSNIYTLLDLLNNSQEDLMKMEHLRLEDIKQILVILEKHF-AIDLPKKNF-----------
18_Liquidambar    275 DQSELPPRIYTCLKRSNIHTLLDLLNNNQEDLMKIEHFRIEDVKKILGILEKHF-AIDLPKNKF-----------
19_Papaver        275 DQSELSPRIYNCLKRSNIYTLLDLLNKSQEDLMKIDHFRIEDVKQILDILQKRF-AIDLPKNKFSFLNPLE----
20_Ananas         275 DQLELPSRTYNCLKRSNIHTLLDLLSNSQEDLMRIEYFRMEDVKQILDTLQKHF-AIDLPKNKFSF---------
28_Liriodendron   275 DQSELPPRTYNCLKRSNIHTLLDLLSNSQEDLMRIEHFRIEDVKQILDILQKHF-TIDLPKNKF-----------
30_Magnolia       275 DQSELPPRTYNCLKRSNIHTLLDLLSNSQEDLMRIEHFRIEDVKQILDILQKHF-TIDLPKNKF-----------
32_Nymphaea       274 DQLELPPRTYNCLKRSNIHTLLDLLSKSQEDLMRIEHFRVEDVKQIFDILQKGF-TIDLLKNFK-NSNQFESR--
33_Amborella      275 DQLELSPRTYNCLKRSNIHTLLDLLSKSQEDLRKIQHFRVEDVKRVLDILKKRF-AMNLFT--------------
35_Picea          276 DQLEFPPRVYNCLRRANIHTLSDLLNYSREDLMRIERFGKESVEQVFEILRKRF-AIDPPRN-------------
44_Ginkgo         275 DQSELPPRVYNCLKRVNIHTLSDLLNYSRDDPTRIENFGNQSVEQILEVLQKRF-AIDLPKKKF-----------
51_Physcomitrium  385 DQLRIPSKAYNSLKRANINTVSDLLDYTQDDLLSIPNFGRKSVDDILEALQAQF-SIDLPENNPLCN--------
```

**Fig. S4.** Sequence alignment β subunits from PEP of angiosperms with those of the RNAPs from *E. coli*, *T. thermophilus* and Nostoc. The residues conserved more than 50 % are in red, those mutated in similar residues are in blue. The strictly conserved residues described by Lane & Darst (Lane & Darst, 2010) are highlighted in gray. The blue triangles show mutations observed among the strictly conserved residues described (Lane & Darst, 2010). The non-conservative mutations, at least three in a row in the β or β' domain in *E. coli* and *T. thermophilus*, are highlighted in green and displayed on the *E. coli* structure (PDB entry: 6GH5). Those colored in orange are nearby to the DNA, those in green are located at the surface of the subunits. The domains described for all-RNA polymerase (a) and the bRNAPs (b) are also given and highlighted in yellow and cyan respectively. The name of the RNAP domains are also given and highlighted in purple and green (Lane & Darst, 2010; Sutherland & Murakami, 2018).

```
                                                              β1 domain: Q22-N130 + V336-S392
                                                    βa1:P16-L30           βa2:G43-R49
                                                    βb1:P16-L30           βb2:G43-G60
T. thermophilus     1 ---------------MEIKRFGRIREVIPLPPLTEIQVESYRRALQADVPPEKRENVGIQAAFRETFPIEEEDKGKGGLV
E. coli             1 ------MVYSYTEKKRIRKDFGKRPQVLDVPYLLSIQLDSFQKFIEQDPEG----QYGLEAAFRSVFPIQSYS---GNSE
0_Nostoc            1 -----------------MTKETYMEPAFLLPDLIEIQRSSFRWFLEEGLIEEL-----------NSFSPITDYTGKLELH
1_Litchi            1 ------------------MRGDVNVRMSTIPGFNQIQFEGFCRFIDQGLTEEL-----------YKFPKIEDTDQEIEFQ
2_Arabidopsis       1 ------------------MLGDEKEGTSAIPGFNQIQFEGFYRFIDQGLIEEL-----------AKFPKIEDIDHEIEFQ
3_Gossypium         1 ------------------MLGDENGEMSTIPGLNQIQFEGFCGFMDRGLTEEL-----------YKFPKIEDTEQEIEFQ
5_Ricinus           1 ------------------MLGDGNEGMSTIPGLNQIQFEGFCRFIDQGLTEEL-----------YKFPKIEDTDQEIEFQ
6_Rosa              1 ------------------MLGGGNEAISTIPGFNQIQFEGFCRFIDQGLTEEL-----------YKFPKIEDTDQEIEFQ
9_Cucumis           1 MMNKQIMGFFFYKWEINKMLGGGNERMSTIPGFNQIQFEGFCRFIDHGLTEEL-----------SKFPKIEDTDQEIEFQ
11_Nicotiana        1 ------------------MLGDGNEGISTIPGFNQIQFEGFCRFIDQGLTEEL-----------YKFPKIEDTDQEIEFQ
13_Syringa          1 ------------------MLGDGNEGMSTIPGFNQIQFEGFCRFIDQGLTEEL-----------YKFPKIEDTDQELEFQ
18_Liquidambar      1 ------------------MLRDGNEGMSTIPGLNQIQFEGFCRFIDQGLTEEL-----------YKFPKIEDTDQEIEFQ
19_Papaver          1 ------------------MLRDGNEGMSTIPGFSQIQFEGFCRFIDQGLMEEL-----------YKFPKIEDIDQEIEFQ
20_Ananas           1 ------------------MLRNGNEGMSTIPGFSQIQFEGFCRFINQGLTEEF-----------HKFPKIEDTDQEIEFK
28_Liriodendron     1 ---------MFSINGKLKMLRDGNEGMSTIPGFSQIQFEGFCRFIDQGLTEEL-----------HKFPKIEDTDQEIEFQ
30_Magnolia         1 ---------MFSINGKLKMLRDGNEGMSTIPGFSQIQFEGFCRFIDQGLTEEL-----------HKFPKIEDTDQEIEFQ
32_Nymphaea         1 -----------------MLRDGGDEEMFTIPGFSQIQFEGFCRFIDQGLMEEL-----------HQFPKIEDTDQEIEFQ
33_Amborella        1 ------------------MLRDGNEGMSTIPGFSQIQFEGFCRFVDQGLAEEL-----------HKFPKIEDTDQEIEFQ
35_Picea            1 ------------------MRLDENEGAFTIPEFGKIQFEGFCRFIDQGLMEEL-----------HNFPKIEDTDKEIESR
44_Ginkgo           1 ------------MDGKLPMLLDENKGTSTIPGFGQIQFEGFCRFIDQGLIEEL-----------SNFPEIEYTDQEIESR
51_Physcomitrium    1 -MKK-IITL---SAPPPSQFSFLSEFQFSLPELRQIQFKSYYYFIYKNLISEL-----------NIFPEIFDLNQEFQFE

                                       β1 domain: Q22-N130 + V336-S392
                                  βa3:F78-I101                      βa4:E112-F148
                              βb3:L64-I101                          βb4:D111-F148
T. thermophilus    66 LDFLEYRLGEPPFPQDECREKDLTYQAPLYARLQLIHKD-------TGLIKEDEVFLGHIPLMTEDGSFIINGADRVIVS
E. coli            68 LQYVSYRLGEPVFDVQECQIRGVTYSAPLRVKLRLVIYEREAPEGTVKDIKEQEVYMGEIPLMTDNGTFVINGTERVIVS
0_Nostoc           53 FLGQNYKLKEPKYSVEEAKRRDSTYAVQMYVPTRLINKE-------TGEIKEQEVFIGDLPLMTDRGTFIINGAERVIVN
1_Litchi           52 LFVETYQLVEPLIKERDAVYESLTYSSELYVSAGLIWKS-------RGDMQEQTIFIGNIPLMNSLGTSIVNGIYRIVIN
2_Arabidopsis      52 LFVETYQLVEPLIKERDAVYESLTYSSELYVSAGLIWKT-------SRNMQEQRIFIGNIPLMNSLGTSIVNGIYRIVIN
3_Gossypium        52 LFVETYQLVEPLIKERDAVYESLTYSSELYVSAGLIWKT-------SKDMQEQTIFIGNIPLMNSLGTSIVNGIYRIVIN
5_Ricinus          52 LFVETYQLVEPLIKEGDAVYESLTYSSELYISAGLIWKT-------SRDMQEQTIFIGNIPLMNSLGTFIINGIYRIVIN
6_Rosa             52 LFVETYQLVEPLIKERDAVYESLTYSSELYVSAGLIWKN-------SRDMQEQTIFIGNIPLMNSLGTSIVNGIYRIVIN
9_Cucumis          70 LFVETYKLVEPLIKERDAVYESLTYSSELYVSAGLIWKT-------RRDMQEQTIFIGNIPLMNSLGTSIVNGLYRIVIS
11_Nicotiana       52 LFVETYQLVEPLIKERDAVYESLTYSSELYVSAGLIWKN-------SRDMQEQTIFIGNIPLMNSLGTSIVNGIYRIVIN
13_Syringa         52 LFVERYQLVEPLIKERDAVYESLTYSSELYVSGGLIWKT-------SRDMQEQTIFIGNIPLMNSLGTSIVNGIYRIVIN
18_Liquidambar     52 LFVETYQLVEPLIKERDAVYESLTYSSELYVSAGLIWKS-------SGDMQEQTIFIGNIPLMNSLGTSIVNGIYRIVIN
19_Papaver         52 LFVETYQLVEPLIKERDAVYESLTYSSELYVPAGLIWKT-------GRDIQEQTIFIGNIPLMNSLGTFIVNGIYRIVIN
20_Ananas          52 LFAERYQLVEPLIKERDAVYESLTYSSELYVPAGLIWKT-------GRDMQEQTVFIGNIPLMNSLGTSIVNGIYRIVIN
28_Liriodendron    61 LFVETYQLVEPLIKERDAVYESLTYSSELYVPAGLIWKT-------GRDMQEQTVFIGNIPLMNSLGTSIVNGIYRIVIN
30_Magnolia        61 LFVETYQLVEPLIKERDAVYESLTYSSELYVPAGLIWKT-------GRDMQEQTVFIGNIPLMNSLGTSIVNGIYRIVIN
32_Nymphaea        53 LFEESYQLVEPLIKERDAVYESITYSSELYVPAGLIWRT-------GRNMQEQTVLLGNIPLMNSLGTSIVNGIYRIVIN
33_Amborella       52 LLVETYQLAEPLIKERDAVYESLTHSSELYVPAGLIWKV-------GRDMQEQTVFIGNIPLMNSLGTFIVNGIYRIVIN
35_Picea           52 LFGNEYELAEPFIKERDAVYQSLTYYSELYVPARSIRRN-------SSKIQKQTVFLGNIPLMNSHGTFVVNGIYRIVVN
44_Ginkgo          58 LSGKKYKSAEPLIEERNAVYQSLTYSSELYVPARLIQKN-------RRKIQKQTVFLGNIPLMNSRGTFVVNGISRIVVD
51_Physcomitrium   65 LLNKEYKLIKPEKTT--IKFHYNTYSSDLYVTCRLLRRK------KKIEIQKQTIFIGSIPLIDYQSTFRINSVTRVIIN

                                             β2 domain: R142-D324
                                                                     βa5:F191-G201
                                  βb5:Y158-L165          βb6:R168-Y202
T. thermophilus   139 QIHRSPGVYFTPDPARP--GRY-IASIIPLPKRGPWIDLEVEPNGVVSMKVN-KRKFPLVLLLRVLGYDQETLARELGAY
E. coli           148 QLHRSPGVFFDSDKGKTHSSGKVLYNARIIPYRGSWLDFEFDPKDNLFVRIDRRRKLPATIILRALNYTTEQILDLF--F
0_Nostoc          126 QIVRSPGVYYKSEIDKN--GRR-TYSASLIPNRGAWLKFETDRNDLVWVRIDKTRKLSAQVLLKALGLSDNEIFDAL-RH
1_Litchi          125 QILQSPGIYYQSEFDHN--GIL-IYAGTIISDWGGRLELEIDRKARIWARVSRKQKISILVLSSAMGSNLREILENI-CY
2_Arabidopsis     125 QILQSPGIYYQSELDHN--GIS-VYTGTIISDWGGRLELEIDKKARIWARVSRKQKISILVLSSAMGLNLREILENV-CY
3_Gossypium       125 QILQSPGIYYRSELDHN--GIS-VYTGTIISDWGGRLELEIDRKARIWARVSRKQKISILVLSSAMGSNLREILENV-CY
5_Ricinus         125 QILQSPGIYYRSELDHN--GIS-VYTGTIISDWGGRVELEIDRKARIWARVSRKQKISILVLLSAMGLNLREILENV-RY
6_Rosa            125 QILQSPGIYYRSELDHN--GIS-VYTGIIISDWGGRLELEIDRKARIWARVSRKQKISILVLSSAMGSNLREILENA-RY
9_Cucumis         143 QILQSPGIYYRSELDHN--GIS-VYTGTIISDWGGRLELEIDRKARIWARVSRKQKISILVLSSAMGSNLREILENV-CY
11_Nicotiana      125 QILQSPGIYYRSELDHN--GIS-VYTGTIISDWGGRSELEIDRKARIWARVSRKQKISILVLSSAMGLNLREILENV-CY
13_Syringa        125 QILQSPGIYYRSELDHN--GIS-VYTGTIISDWGGRSELEIDRKARIWARVSRKQKISILVLSSAMGSNLREILDNV-CY
18_Liquidambar    125 QILQSPGIYYRSELDHN--GIS-IYTGTIISDWGGRLELEIDRKARIWARVSRKQKISILVLSSAMGSNLREILENV-CY
19_Papaver        125 QILQSPGIYYRSELDHN--GIS-VYTATIISDWGGRSELEIDRKARIWARVSRKQKISILVLSSAMGSNLREILDNV-CY
20_Ananas         125 QILQSPGIYYRSELDHN--GIS-VYTSTIISDWGGRSEFEIDRKARIWARVSRKQKISILVLSSAMGSNLREILDNV-CY
28_Liriodendron   134 QILQSPGIYYRSELDHN--GIS-VYTGTIISDWGGRSELEIDRKARIWARVSRKQKISILVLSSAMGSNLREILDNV-CY
30_Magnolia       134 QILQSPGIYYRSELDHN--GIS-VYTGTIISDWGGRSELEIDRKARIWARVSRKQKISILVLSSAMGSNLREILDNV-CY
32_Nymphaea       126 QILQSPGIYYSTGLDHN--GIS-VYTGTIISDWGGRSELEIDRKERIWARVSRKQKISILVLSSAMGSSLREILDNV-CY
33_Amborella      125 QILQSPGIYYSSELDHN--GIS-VYTGTIISDRGGRSELEIDRKARIWARVSRKQKISILVLPSAMGSNLREILDNV-CY
35_Picea          125 QILISPGIYYRSELDHN--RINYIYTGTLISDWGRRSKLEIDVGERIWARVSRKRKISIPVLLSAMGLNLEEILDNT-RY
44_Ginkgo         131 QILRSPGIYYSSEPGHN--GIA-IYTGTIISDWGGRPKLEIDGKTRIWARVSRKQKVSIPVLLSAMGSNFEEILDNV-CY
51_Physcomitrium  137 QILRSPGIYYNSELDHN--GIS-IYTGTIISDWGGRLKLEIDSKTRIWARISKKRKVSILVLLLAMGLTIKQILDSV-CS
```

```
T. thermophilus  215 GELVQGLM------------------------------------------------------------------------
E. coli          226 EKVIFEIRDNKLQMELVPERLRGETASFDIEANGKVYVEKGRRITARHIRQLEKDDVKLIEVPVEYIAGKVVAKDYIDES
0_Nostoc         202 PEYFQKTIE-----------------------------------------------------------------------
1_Litchi         201 PEIFLSFLTD----------------------------KE----------------------------------------
2_Arabidopsis    201 PEIFLSFLTD----------------------------KE----------------------------------------
3_Gossypium      201 PEIFLSFLTD----------------------------KE----------------------------------------
5_Ricinus        201 PEIFLSFLND----------------------------KE----------------------------------------
6_Rosa           201 PEIFLSFLND----------------------------KE----------------------------------------
9_Cucumis        219 PEIFLSFLND----------------------------KE----------------------------------------
11_Nicotiana     201 PEIFLSFLSD----------------------------KE----------------------------------------
13_Syringa       201 PEIFLSFLND----------------------------KE----------------------------------------
18_Liquidambar   201 PEIFLSFLND----------------------------KE----------------------------------------
19_Papaver       201 PEIFLSFPND----------------------------KE----------------------------------------
20_Ananas        201 PEIFLSFPND----------------------------KE----------------------------------------
28_Liriodendron  210 PEIFLSFPND----------------------------KE----------------------------------------
30_Magnolia      210 PEIFLSFPND----------------------------KE----------------------------------------
32_Nymphaea      202 PEIFLSFPNE----------------------------KE----------------------------------------
33_Amborella     201 PEILLYFPNE----------------------------KE----------------------------------------
35_Picea         202 PEKIFFLLKK----------------------------KKGRW-------------------------------------
44_Ginkgo        207 PEIFLSFL-------------------------------NGRQ-------------------------------------
51_Physcomitrium 213 SKIFLDFLKE----------------------------KK----------------------------------------
```

β2 domain: R142-D324

βb7:A234-K280

```
T. thermophilus  223 -------------------------DE--------------------SVFAMRPEEALIRLFTLLRPGDPPKR--DKAVA
E. coli          306 TGELICAANMELSLDLLAKLSQSGHKRIETLFTNDLDHGPYISETLRVDPTNDRLSALVEIYRMMRPGEPPTR--EAAES
0_Nostoc         211 -----------------------------------------------KEGQFSEEEALMELYRKLRPGEPPTVLG--GQQ
1_Litchi         213 -----------------------------------------------KKKIGSKENAILEFYQQFACVGGDPIFSESLCK
2_Arabidopsis    213 -----------------------------------------------KKKIGSKENAILEFYQQFSCVGGDPIFSESLCK
3_Gossypium      213 -----------------------------------------------KKKIGSKENAILEFYQQFSCVGGDPVFSESLCK
5_Ricinus        213 -----------------------------------------------KKKIGSKENAILEFYQQFTCVGGDPVFSESLCK
6_Rosa           213 -----------------------------------------------KKKIGSKENAILEFYQQFACVGGDPVVSESLCK
9_Cucumis        231 -----------------------------------------------KKKIGSKENAILEFYQQFSCVGGDPVFSESLCK
11_Nicotiana     213 -----------------------------------------------RKKIGSKENAILEFYQQFACVGGDPVFSESLCK
13_Syringa       213 -----------------------------------------------RKKIGSKENAILEFYQQFACVGGDPVFSESLCK
18_Liquidambar   213 -----------------------------------------------KKKIGSKENAILEFYQQFACVGGDPVFSESLCK
19_Papaver       213 -----------------------------------------------KKKIGSKENAILEFYQQFACVGGDPVFSESLCK
20_Ananas        213 -----------------------------------------------KKKIGSKENAILEFYQQFACVGGDPVFSESLCK
28_Liriodendron  222 -----------------------------------------------KKKIGSRENAILEFYQQFACVGGDPVFSESLCK
30_Magnolia      222 -----------------------------------------------KKKIGSRENAILEFYQQFACVGGDPVFSESLCK
32_Nymphaea      214 -----------------------------------------------KKKISSKENAILEFYQKFACVGGDPVFSESLCK
33_Amborella     213 -----------------------------------------------KKKIGSKENAILEFYQQFSCVGGDPVFSESLCK
35_Picea         217 -------------------------ER--------------------EEYIWSKEKAILEFYKKLYCVSGDLVFSESLCK
44_Ginkgo        219 -------------------------KR--------------------KKYLRSEENAILEFHKKLYCVGGDLVFSESLCK
51_Physcomitrium 225 -------------------------KK--------------------KEHLQSTEDAMVELYKQLYYIGGDLLFSESIRK
```

β2 domain: R142-D324

βa6:D323-V355

βb7:A234-K280 βb8:V302-G316 βb9:D323-M359

```
T. thermophilus  256 YVYGLIADPRRYDLGEAGRYKAEEKLGIRLSGRTLARFEDGEFKDEVFLPTLRYLFALTAGVPGHEVDDIDHLGNRRIRT
E. coli          384 LFENLFFSEDRYDLSAVGRMKFNRSLLREEIEGS------GILSKDDIIDVMKKLIDIRNGK--GEVDDIDHLGNRRIRS
0_Nostoc         242 LLDSRFFDPKRYDLGRVGRYKLNKKLRLSVPDTMRVLTSS------DILAAVDYLINLEYDI--GNIDDIDHLGNRRVRS
1_Litchi         246 ELQKKF-FHQRCELGRIGRRNMNRRLNLNIPQNNTFLLPR------DVLAAADHLIELKFGM--GTLDDMNHLKNKRIRS
2_Arabidopsis    246 ELQKKF-FHQRCELGRIGRRNINWRLNLNIPQNNIFLLPR------DVLAAADHLIGMKFGM--GTLDDMNHLKNKRIRS
3_Gossypium      246 ELQKKF-FQQRCELGRIGRRNMNQRLNLNIPQNNTFLLPR------DILAAADRLIGMKFGM--GPLDDMNHLKNKRIRS
5_Ricinus        246 ELQKKF-FQQRCELGRIGRLNMNRRLNLDIPHNNTFLLPR------DILAAADHLIGMKFGM--GTLDDMNHLKNKRIRS
6_Rosa           246 ELQKKF-FQQRCELGRIGRRNMNRRLNLDIPQNNTFLLPR------DILAAADHLIGMKFGM--GTLDDMNHLKNKRIRS
9_Cucumis        264 ELQKKF-FQQRCELGRIGRRNLNQRLNLDIPENNTFLLPR------DILAAADHLIGLKFGM--GTLDDMNHLKNKRIRS
11_Nicotiana     246 ELQKKF-FQQRCELGRIGRRNMNRRLNLDIPQNNTFLLPR------DILAAADHLIGLKFGM--GALDDMNHLKNKRIRS
13_Syringa       246 ELQKKF-FQQRCELGRIGRRNMNRRLNLDIPQNNTFLLPR------DILAAADHLIELKFGM--GTLDDMNHLKNKRIRS
18_Liquidambar   246 ELQKKF-FQQRCELGRIGRRNMNRRLDLNIPQNNTFLLPR------DILAAADHLIGMKFGM--GTLDDMNHLKNKRIRS
19_Papaver       246 ELQKKF-FQQRCELGRIGRRNMNRRLNLDIPQNNTFLLPR------DVLAAADHLIGMKFGM--GTLDDMNHLKNKRIRS
20_Ananas        246 ELQKKF-FQQRCELGRIGRRNMNRRLNLDIPQNNTFLLPR------DVLAAADHLIGMKFGM--GTFDDMNHLKNKRIRS
28_Liriodendron  255 ELQKKF-FQQRCELGRIGRRNMNRRLNLDIPQNNTFLLPR------DVLAAADHLIGMKFGM--GTLDDMNHLKNKRIRS
30_Magnolia      255 ELQKKF-FQQRCELGRIGRRNMNRRLNLDIPQNNTFLLPR------DVLSAADHLIRMKFGM--GTLDDMNHLKNKRIRS
32_Nymphaea      247 ELQKKF-FQQRCELGRIGRRNMNQRLNLDIPQNNTFLLPR------DVLAAADHLIGMKFGM--GTLDDMNHLKNKRIRS
33_Amborella     246 ELQKRF-FQQRCELGRIGRQNMNQRLNIDIPQNNTFLLPR------DVLAATDHLIGMKFGM--GTLDDMNHLKNKRIRS
35_Picea         252 ELQEKF-FRQRCELGKIGRRNPNQKLNLDIPENEIFSLPQ------DVLAAVDYLIGVKFGM--GTLDDIDHLRNRRIRS
44_Ginkgo        254 ELQKKS-LQQRCELGRIGRRNPNQKLNLDIPENEIFSLPQ------DVLAAADYSIRVKFGM--GTLDDMDHLKNKRIRS
51_Physcomitrium 260 ELQKKF-FQQRCELGKIGRLNVNKKLSLDIPENEFFLLPQ------DILAAIDYLIKIKFGI--GTLDDIDHLKNRRIRS
```

```
                                 β1 domain: Q22-N130 + V336-S392
                       βa6:D323-V355                         βa7:S375-E421
                       βb9:D323-M359                    βb10:L367-V474
T. thermophilus   336 VGELMTDQFRVGLARLARGVRERMLMGSE--DSLTPAKLVNSRPLEAAIREFFSRSQLSQFKDETNPLSSLRHKRRISAL
E. coli           456 VGEMAENQFRVGLVRVERAVKERLSLGDL--DTLMPQDMINAKPISAAVKEFFGSSQLSQFMDQNNPLSEITHKRRISAL
0_Nostoc          314 VGELLQNQVRVGLNRLERIIRERMTVS--DAEVLTPASLVNPKPLVAAIKEFFGSSQLSQFMDQTNPLAELTHKRRLSAL
1_Litchi          317 VADLLQDQFGLALVRLENVVRGAIGGAIRHKLMPTPQNLVTSTPLTTTYDSFFGLHPLSQVLDRTNPLTQIVHGRKLSYL
2_Arabidopsis     317 VADLLQDQLGLALARLENVVKGTISGAIRHKLIPTPQNLVTSTPLTTTYESFFGLHPLSQVLDRTNPLTQIVHGRKLSYL
3_Gossypium       317 VADLLQDQFGLALVRLENVVRGTICGAIRHKLIPTPQNLVTSTPLTTTYESFFGLHPLSQVLDRTNPLTQIVHGRKLSYL
5_Ricinus         317 VADLLQDQFGLALIRLENVVRGTICGAIRHKLIPTPQTLVTSTPLTTTYESFFGLHPLSQVLDRTNPLTQIVHGRKSSYL
6_Rosa            317 VADLLQDQFGLALVRLENMVRGTICGAIKHKLIPTPQNLVTSTPLTTTYESFFGLHPLSQVLDQTNPLTQIVHGRKSSYL
9_Cucumis         335 VADLLQDQFGLALVRLENMVRGTICGAIRHKLIPTPQNLVTSTPLTTTFESFFGLHPLSQVLDRTNPLTQIVHGRKLSYL
11_Nicotiana      317 VADLLQDQFGLALVRLENVVRGTICGAIRHKLIPTPQNLVTSTPLTTTYESFFGLHPLSQVLDRTNPLTQIVHGRKLSYL
13_Syringa        317 VADLLQDQFGLALVRLENVVRGTICGAIRHKLIPTPQNLVTSTPLTTTYESFFGLHPLSQVLDRTNPLTQIVHGRKLSYL
18_Liquidambar    317 VADLLQDQFGLALVRLENAVRGTICGAIRHKLIPTPQNLVTSTPLTTTYEYFFGLHPLSQVLDRTNPLTQIVHGRKSSYL
19_Papaver        317 VADLLQDQFGLALVRLENVVRGTICGAIRHKFIPTPHNLVTSTPLTTTYESFFGLHPLSHVLDRTNPLTQIVHGRKLSYL
20_Ananas         317 VADLLQDQFGLALVRLENAVRGTICGAIRHKLIPTPQNLVTSTSLTTTYESFFGLHPLSQVLDRTNPLTQIVHGRKLSYL
28_Liriodendron   326 VADLLQDQFGLALVRLENAVRGTICGAIRHKLIPTPQNLVTSTPLTTTYESFFGLHPLSQVLDRTNPLTQIVHGRKSSYL
30_Magnolia       326 VADLLQDQFGLALVRLENAVRGTICGAIRHKLIPTPQNLVTSTPLTTTYESFFGLHPLSQVLDRTNPLTQIVHGRKSSYL
32_Nymphaea       318 VADLLQDQFGLALVRLENVVRGTICGAIRHKLIPTPRNLVTSTPLTTTYESFFGLHPLSQVLDRTNPLTQIVHGRKSSYL
33_Amborella      317 VADLLQDQFGLALVRLENVVRGTICGAIRHKFIPTPQNLVTSTPLTTTYESFFGLHPLSQVLDRTNPLTQIVHGRKSSYL
35_Picea          323 VADLLQNQFRLALGRLEDAVRRTIHRATKRR--STPQNLVTSTLLKNTFQDFFGSHPLSQFLDQTNPLTEIAHGRKLSHL
44_Ginkgo         325 VADLLQNQFGLALGRLVNSVRRTIRRATKCKCLPTPKNLVTSTPLTTTFQDFFGLHPLSQFLDQTNPLTEAVHRRKLSYL
51_Physcomitrium  331 VADLLQDQLKLALIRLENSVRQVMRRTTKRKRLLSPKNLITQTPLIATFKEFFGSHPLSQFLDQTNSLAEIVHKRRLSSL

                 Fork-loop 2: S411-R428
                                         βa8:D426-Y471
                                        βb10:L367-V474                      βb11:V479-L503
T. thermophilus   414 GPGGLTRERAGFDVRDVHRTHYGRICPVETPEGANIGLITSLAAYARVDELGFIRTPYRRVVGGVVT--DEVVYMTATEE
E. coli           534 GPGGLTRERAGFEVRDVHPTHYGRVCPIETPEGPNIGLINSLSVYAQTNEYGFLETPYRKVTDGVVT--DEIHYLSAIEE
0_Nostoc          392 GPGGLTRERAGFAVRDIHPSHYGRICPIETPEGPNAGLIGSLATHARVNQYGFLETPFRPVENARVRFDLPPVYMTADE-
1_Litchi          397 GPGGLTGRTASFRIRDIHPSHYGRICPIDTSEGINVGLIGSLAIHARIGYWGSLESPFYEIFEKSKK--MRMLYLSPSID
2_Arabidopsis     397 GPGGLTGRTANFRIRDIHPSHYGRICPIDTSEGINVGLIGSLSIHARIGDWGSLESPFYELFEKSKKARIRMLFLSPSQD
3_Gossypium       397 GPGGLTGRTANFRIRDIHPSHYGRICPIDTSEGINVGLIGSLAIHARIGHWGSLESPFYKIFERSKK--AQMLYLSPSRD
5_Ricinus         397 GPGGLTGRTASFRIRDIHPSHYGRICPIDTSEGINVGLIGSLAIHAKIGHWGSLESPFYVISEESKK--VRMFYLSPNRE
6_Rosa            397 GPGGLTGRTASFRIRDIHPSHYGRICPIDTSEGINVGLIGSLAIHAKIGYWGSLESPFYEISERSKK--VRMLYLSPSKD
9_Cucumis         415 GPGGLTGRTASFRIRDIHPSHYGRICPIDTSEGINVGLIGSLAIHARIGHWGSLETPFYEISERSKK--VRMLYLSPSRD
11_Nicotiana      397 GPGGLTGRTASFRIRDIHPSHYGRICPIDTSEGINVGLIGSLAIHARIGHWGSLESPFYEISERSTG--VRMLYLSPGRD
13_Syringa        397 GPGGLTGRTASFRIRDIHPSHYGRICPIDTSEGINVGLIGSLAIHARIGHWGSLESPFYEISERSTG--VRMLYLSPGRD
18_Liquidambar    397 GPGGLTGRTASFRIRDIHPSHYGRICPIDTSEGINVGLIGSLAIHARIGRWGSLESPFYEISERSKK--VRMLYLSPSRD
19_Papaver        397 GPGGLTGRTASFRIRDIHPSHYGRICPIDTSEGINVGLIGSLAIHARIGHWGSLESPFFEIDERF--KGVRVVYLSPSRD
20_Ananas         397 GPGGLTGRTASFRIRDIHPSHYGRICPIDTSEGINVGLIGSLAIHVRIGHWGSIESPFYEISEKQKEKEPQMVYLSPNRD
28_Liriodendron   406 GPGGLTGRTASFRIRDIHPSHYGRICPIDTSEGINVGLIGSLAIHARIGHWGSIESPFYEISERS--KEVQMVYLSPSRD
30_Magnolia       406 GPGGLTGRTASFRIRDIHPSHYGRICPIDTSEGINVGLIGSLAIHARIGHWGSIESPFYEISERS--KEVQMVYLSPSRD
32_Nymphaea       398 GPGGLTGRTASFRIRDIHPSHYGRICPIDTSEGINVGLIGSLAIHARVGDWGSIETPFYEISERS--KEEQMVYLSPSRD
33_Amborella      397 GPGGLTGRTASFRIRDIHPSHYGRICPIDTSEGINVGLIGSLAIHARIGDWGSIRSPFYEISERS--KEEQMVYLSPRRD
35_Picea          401 GPGGLTGRTASFRTRDIHPSYYGRICPIDTSEGMNAGLVASLSIHAKIGQCGSLQSPFYKISERSRE--EHMVYLLPGED
44_Ginkgo         405 GPGGLTRRTASFRIRDIHPSHYGRICPIETSEGINAGLVASLAIHAKIGHCGSLRSPFHKISEGSKE--EHMVYPSPGE-
51_Physcomitrium  411 GPGGVTRRTAGFQVRDIHFSHYTRICPIETSEGMNAGLIASLAIHANVNNWGFLESPFYKISKNVKE--EKIINLSAGED
                                ▲

                   βa9:Y485-A499                              βa10:V529-D590
                    βb11:V479-L503                   βb12:I508-V613
T. thermophilus   492 --DRYTIAQANTPLEGNRIAAERV-VARRKGEPVIVSPEEVEFMDVSPKQVFSVNTNLIPFLEHDDANRALMGSNMQTQA
E. coli           612 --GNYVIAQANSNLDEEGHFVEDLVTCRSKGESSLFSRDQVDYMDVSTQQVVSVGASLIPFLEHDDANRALMGANMQRQA
0_Nostoc          471 -EDDLRVAPGDIPVDENGHIIGPQVPVRYRQEFSTTTPEQVDYVAVSPVQIVSVATSMIPFLEHDDANRALMGSNMQRQA
1_Litchi          475 EYCMV--AAGNSLALSQGIQEEQVVPTRYRQEFLTIAWERVHLRSIFPSQYFSIGASLIPFIEHNDANRALMSSNMQRQA
2_Arabidopsis     477 EYYMI--AAGNSLALNRGIQEEQAVPARYRQEFLTIAWEEVHLRSIFPFQYFSIGASLIPFIEHNDANRALMSSNMQRQA
3_Gossypium       475 EYYMV--AAGNSLALNQGIQEEQVVPARYRQEFLTIAWEQVHLRSIFPFQYFSIGASLIPFIEHNDANRALMSSNMQRQA
5_Ricinus         475 EYHMV--AAGNSLALNRGVQEEQVAPARYRQEFLTIAWEQVHLRSIFPFQYFSIGASLIPFIEHNDANRALMSSNMQRQA
6_Rosa            475 EYYMI--AAGNSLALNRGIQEEQVVPARYRQEFLTVEWEQVHLRSIFPFQYFSIGASLIPFIEHNDANRALMSSNMQRQA
9_Cucumis         493 EYYMV--ATGNSLALNPGIQEEQIVPARYRQEFLTIEWEQVHLRSIFPFQYFSIGASLIPFIEHNDANRALMSSNMQRQA
11_Nicotiana      475 EYYMV--AAGNSLALNQDIQEEQVVPARYRQEFLTIAWEQVHLRSIFPFQYFSIGASLIPFIEHNDANRALMSSNMQRQA
13_Syringa        475 EYYMV--AAGNSLALNQDIQEEQVVPARYRQEFLTIAWEQVHLRSIFPFQFFSIGASLIPFIEHNDANRALMSSNMQRQA
18_Liquidambar    475 EYYMV--AAGNSLALNQGIQEEQVVPARYRQEFLTIAWEQVHLRSIFPFQYFSIGASLIPFIEHNDANRALMSSNMQRQA
19_Papaver        475 EYYMV--SAGNSLALNQGIQEEQVVPARYRQEYLTIAWEQIHLRSIFPFQYFSIGASLIPFIEHNDANRALMSSNMQRQA
20_Ananas         477 EYYMV--AAGNSLALNRGIQEEQVVPARYRQEFLTIAWEQIHLRSILPFQYFSIGASLIPFIEHNDANRALMSSNMQRQA
28_Liriodendron   484 EYYMV--AAGNSLALNWGIQEEQVVPARYRQEFLTIAWEQIHLRSIFPFQYFSIGASLIPFIEHNDANRALMSSNMQRQA
30_Magnolia       484 EYYMV--AAGNSLALNWGVQEEQVVPARYRQEFLTIAWEQIHLRSIFPFQYFSIGASLIPFIEHNDANRALMSSNMQRQA
32_Nymphaea       476 EYYMV--AAGNSLALTRGIQEEEVGPARYRQEFLTIAWEQIHLRNIYPFQYFSIGASLIPFIEHNDANRALMSSNMQRQA
33_Amborella      475 EYYMVMVAAGNSLALNQDIQDEQVVPARYRQEFVTIAWEHIDLRSIYPLQYFSIGASLIPFIEHNDANRALMSSNMQRQA
35_Picea          479 EDEYYRIATGNSLALNQGIQEEQITPARYRQEFIVIAWEQIHFRSIFPFQYFSVGVSLIPFLEHNDANRALMGSNMQRQA
44_Ginkgo         482 -DEYYRIATGNSLALNQGIQEEQVTPARYRQEFLAIAREQIHFRSIFPFQYFSVGVSLIPFLEHNDANRALMGSNMQRQA
51_Physcomitrium  489 --EYYRIATGNCLALDQGTQKIQITPARYRQEFLAIAWEQIHLRSIYPLQYFSVGVSLIPFLEHNDANRALMGSNMQRQA
```

```
                           βa10:V529-D590
                           βb12:I508-V613                                    βb13:Y623-R808
T. thermophilus  569 VPLIRAQAPVVMTGLEERVVRDSLAALYAEEDGEVAKVDGNRIVVRY-ED------GRLVEYPLRRFYRSNQGTALDQRP
E. coli          690 VPTLRADKPLVGTGMERAVAVDSGVTAVAKRGGVVQYVDASRIVIKVNEDEMYPGEAGIDIYNLTKYTRSNQNTCINQMP
0_Nostoc         550 VPLLKPERPLVGTGLEAQGARDSGMVVVSRTDGDVTYVDATEIRVRPKPN------TTEIRYPLSKYQRSNQDTCLNQKP
1_Litchi         553 VPLSRSEKCIVGTGLERHVALDSGVPAIADHEGRVLYTDIDKIVLSG--N------GDTIGIPLVMYQRSNKNTCMHQKT
2_Arabidopsis    555 VPLSRSEKCIVGTGLERQVALDSGVPAIAEHEGKILYTDTEKIVFSG--N------GDTLSIPLIMYQRSNKNTCMHQKP
3_Gossypium      553 VPLSRSEKCIVGTGLERQVALDSGVPAIADHEGKIISTDTDKIILSG--N------GDALGIPLVMYQRSNKNTCMHQTA
5_Ricinus        553 VPLSRSEKCIVGTGLERQVALDSGVPAIAEREGKIIYTDIDKIILSG--N------GDTLRIPLVMYQRSNKNTCMHQKP
6_Rosa           553 VPLSRSEKCIVGTGLERQVALDSGVPAIAEHEGKIIYTDTDKIILSG--N------GDTLNIPLVIYQRSNKNTCMHQKP
9_Cucumis        571 VPLSRSEKCIVGTGLERQVARDSGVAAIAEHGGKIIYTDTDKIIFSG--N------GHTRRIPLVMYQRSNKNTCMQQKS
11_Nicotiana     553 VPLSRSEKCIVGTGLERQAALDSGALAIAEREGRVVYTNTDKILLAG--N------GDILSIPLVIYQRSNKNTCMHQKL
13_Syringa       553 VPLSRPEKCIVGTGLERQAALDSGALAIAEREGKIIYTDTEKILFSG--N------GDTLSIPLVMYQRSNKNTCMHQKP
18_Liquidambar   553 VPLSRSEKCIVGTGLERQAALDSGIPVLAEHEGKIVYTDTDKIILSG--N------GDTLSIPLVMYQRSNKNTCMHQKP
19_Papaver       553 VPLSRSEKCIVGTGLERQVALDSGVSAIAEHEGKVVSTDTDKIVFSG--N------GDTLSIPLVMYQRSNKNTCMHQKS
20_Ananas        555 VPLSRSEKCIVGTGLERQTALDSGVLVIAEHEGKIIYTDTHKIMLSS--N------GNTISIPLVMYQRSNKNTCMHQKP
28_Liriodendron  562 VPLSRSEKCIVGTGLECQAALDSGVSAIAEHEGKIVYTDTDKIVLSG--N------GDTISIPLVMYQRSNKNTCMHQKP
30_Magnolia      562 VPLSRSEKCIVGTGLECQAALDSGVSAIAEHEGKIVSTDTDKIVLSG--N------GNTISIPLVMYQRSNKNTCMHQKP
32_Nymphaea      554 VPLSQSEKCIVGTGLERQAALDSGGSAIAEREGKIIYTDAEKIVLSG--N------GDTISIPLVMYQRSNKNTWMHQKP
33_Amborella     555 VPLSRSEKCIVGTGLERQAALDSGGSAIAQHEGKVIYTDTEKILLSG--N------GDTISIPLLMYQRSNKNTCMHQKP
35_Picea         559 VPLFQPEKCIAGTGLEGQAALDSGSVAIATQEGRIEYIDAVNITSSV--N------GDTVRTELVIYQRSNTNTCTHQKP
44_Ginkgo        561 IPLFQPEKCITGTGLEGQVALDSGSVTIAIQEGRIEYTDAENITFSF--N------GDTIGTELVLYQRSNKNTCMHQKP
51_Physcomitrium 567 VPLIKLEKCIVGTGLESQVALDSGNVMITKQSEKIMYTDGKKISLLN-NT------NETVNTHLIIYQRSNNSTCIHQKP
```

```
                                                                                       β-flap
                                               βa11:F665-K716
                                      βb13:Y623-R808
T. thermophilus  642 RVVVGQRVRKGDLLADGPASENGFLALGQNVLVAIMPFDGYNFEDAIVISEELLKRDFYTSIHIERYEIEARDTKLGPER
E. coli          770 CVSLGEPVERGDVLADGPSTDLGELALGQNMRVAFMPWNGYNFEDSILVSERVVQEDRFTTIHIQELACVSRDTKLGPEE
0_Nostoc         624 LVRIGEKVVAGQVLADGSSTEGGELALGQNIVVAYMPWEGYNYEDAILISERLVQDDIYTSIHIEKYEIEARQTKLGPEE
1_Litchi         625 QVGRGKCIKKGQVLADGAATVGGELALGKNVLVTYMPWEGYNFEDAVLISERLIYRDIYTSFHIQKYEIQTHVTSQGPER
2_Arabidopsis    627 QVRRGKCIKKGQILADGAATVGGELALGKNILVAYMPWEGYNFEDAVLISECLVYGDIYTSFHIRKYEIQTHVTTQGPER
3_Gossypium      625 RVRRGKCIKKGQILADGAATVGGELALGKNVLVAYMPWEGYNFEDAVLISERLVYEDIYTSFHIRKYEIQTHVTSQGPER
5_Ricinus        625 QVPRGKCIKKGQVLADGAATIGGELALGKNVLVAYMPWEGYNFEDAVLISERLVYEDIYTSFHIRKYEIQTHVTSQGPER
6_Rosa           625 QVQRGKCIKKGQILADGAATVGGELALGKNVLVAYMPWEGYNSEDAVLISERLVYGDIYTSFHIRKYEIQTHVTSNGPER
9_Cucumis        643 QVHQGKCIKKGQILADGAATVGGELALGKNVLVAYMPWEGYNFEDAVLISERLVYEDIYTSFHIRKYEIQTHVTSHGPER
11_Nicotiana     625 QVPRGKCIKKGQILADGAATVGGELALGKNVLVAYMPWEGYNSEDAVLISERLVYEDIYTSFHIRKYEIQTHVTSQGPEK
13_Syringa       625 QVQRGKCIKKGQILADGAATVGGELALGKNVLVAYMPWEGYNFEDAVLISERLVYEDIYTSFHIRKYEIQTHVTSQGPER
18_Liquidambar   625 QVQRSKCIKKGQILADGAATVGGELALGKNVLVAYMPWEGYNFEDAVLISERLVYGDIYTSFHIRKYEIQTHVTSQGPER
19_Papaver       625 QVQRGNCIKKGQILADGVATVGGELALGKNVLVAYMPWEGYNFEDAVLISERLVYGDVYTSFHIRKYEIQTHVTSQGPER
20_Ananas        627 RVRRGKCIKKGQILADGAATVGGELALGKNVLVAYMPWEGYNSEDAVLISERLVYEDIYTSFHIRKYEIQTHVTSQGPER
28_Liriodendron  634 QVRRGKCIKKGQILADGAATVGGELALGKNVLVAYMPWEGYNSEDAVLISERLVYGDIYTSFHIRKYEIQTHVTSQGPER
30_Magnolia      634 QVRRGKCIKKGQILADGAATVGGELALGKNVLVAYMPWEGYNSEDAVLISERLVYGDIYTSFHIRKYEIQTHVTSQGPER
32_Nymphaea      626 QVHRGKYLKKGQILADGAATVGGELALGKNVSVAYMPWEGYNSEDAVLISERLVYDDIYTSFHIRKYEIQTHVTSQGPER
33_Amborella     627 QVHRDKYVKKGQVLADGAATVGGELALGKNVLVAHMPWEGYNFEDAVLISERLVYGDIYTSFHIRKYEIQTHVTSHGPEK
35_Picea         631 QVRQGECVKKGQILADGAATVGGELSLGKNVLVAYMPWEGYNFEDAILISERLVYEDIYTSFHIVRYRIEICMTSQGPER
44_Ginkgo        633 RVRQGECVKKGQILADGAATVEGELSPGKNILVAYMPWEGYNFEDAILISERLVYEDIYTSFHIERHGIRTCMTSQGPER
51_Physcomitrium 640 QVISKKFLKKGQVLTDGAAILKGELTLGKNILVAYMPWEGYNFEDAILISERLIYEDIYTSIHIERYEIESRNTNQGPEK
                                                                       Tip    Helix    Tip
```

```
                                     β-flap
                           βa12:A733-K762                                   βa13:D787-V804
                                      βb13:Y623-R808
T. thermophilus  722 ITRDIPHLSEAALRDLDEEGVVRIGAEVKPGDILVGRTSFKG--ESEPTPEERLLRSIFGEKARDVKDTSLRVPPGEGGI
E. coli          850 ITADIPNVGEAALSKLDESGIVYIGAEVTGGDILVGKVTPKG--ETQLTPEEKLLRAIFGEKASDVKDSSLRVPNGVSGT
0_Nostoc         704 ITREIPNVGEDALRQLDEQGIIRIGAWVEAGDILVGKVTPKG--ESDQPPEEKLLRAIFGEKARDVRDNSLRVPNGEKGR
1_Litchi         705 ITNEIPHLEARLLRNLDQNGIVMLGSWVETGDILVGKLTPQTAKESSYAPEDRLLRAILGIQVSTSKETCLKLPIGGRGR
2_Arabidopsis    707 ITKEIPHLEGRLLRNLDKNGIVMLGSWVETGDILVGKLTPQVAKESSYAPEDRLLRAILGIQVSTSKETCLKLPIGGRGR
3_Gossypium      705 ITNEIPHLEAHLLRNLDKNGIVMLGSWVETGDILVGKLTPQVAKESSYAPEDRLLRAILGIQVSTSKETCLKLPIGGRGR
5_Ricinus        705 ITNEIPHLEAHLLRNLDKNGIVMLGSWVETGDILVGKLTPQMAKESSYAPEDRLLRAILGIQVSTSKETCLKLPIGGRGR
6_Rosa           705 ITNEIPHLEAHLLRNLDKNGIVRLGSWVKTGDILVGKLTPQMAKESSYAPEDRLLRAILGIQVSTSKETCLKLPIGGRGQ
9_Cucumis        723 ITNEIPHLEARLLCNLDKNGIVMLGSWVETGDILVGKLTPQMAKESSYAPEDRLLRAILGIQVSTSKETCLKLPIGGRGR
11_Nicotiana     705 VTNEIPHLEAHLLRNLDKNGIVMLGSWVETGDILVGKLTPQVVKESSYAPEDRLLRAILGIQVSTSKETCLKLPIGGRGR
13_Syringa       705 ITNEIPHLEAHLLRNLDKNGIVMLGSWVETGDILVGKLTPQMVKESSYAPEDRLLRAILGIQVSTSKETCLKLPIGGRGR
18_Liquidambar   705 ITNEIPHLEAHLLRNLDKNGIVMLGSWVETGDILVGKLTPQMAKESSYAPEDRLLRAILGIQVSTSKETCLKLPIGGRGR
19_Papaver       705 ITNEIPHLEAHLLRNLDRNGIVMLGSWVETGDILVGKLTPQMTKESSYAPEDRLLRAILGIQVSTAKETCLKLPIGSRGR
20_Ananas        707 ITKEIPHLEAHLLRNLDRNGIVMLGSWVETGDILVGKLTPQTANESSYSPEDRLLRAILGIQVSTAKETSLKLPIGGRGR
28_Liriodendron  714 ITNEIPHLEAHLLRNLDKNGIVMLGSWIETGDILVGKLTPQTAKESSYAPEDRLLRAILGIQVSTAKETCLKLPIGGRGR
30_Magnolia      714 ITNEIPHLEAHLLRNLDKNGIVMLGSWIETGDILVGKLTPQTAKESSYAPEDRLLRAILGIQVSTAKETCLKLPIGGRGR
32_Nymphaea      706 ITNEIPHLEPYLLRNLDRNGIVMLGSWVETGDVLVGKLTPQTAKESSYAPEDRLLRAILGIQVSTAKETCLKLPIGGRGR
33_Amborella     707 ITNEIPHLEAHLLRNLDRNGIVMLGSWVETGDVLVGKLTPQTAKESSYAPEDRLLRVILGIQVSTAKETCLKLPIGGRGR
35_Picea         711 ITREIPHLDAHSLRHLDENGLVMLGSWIETGDVLVGKLTPQTTEESLCAPEGRLLQTIFGIEVSTARENCLRTPIGGRGR
44_Ginkgo        713 ITKEIPHLDAHLLRHLDENGLVMLGSWIETGDVLVGKLTPQKEEESLCAPEGRLLRTIFGIQVSTARESCLRVPIGGRGR
51_Physcomitrium 720 ITKEIPHLENSVLRHLDKNGLVIPGSWVETGDVLVGKLTPQETEESLRAPEGKLLQAIFGIQVTNAKETCLKVPLNGKGR
```

```
                     β-flap
                     βa13
                     βb13
T. thermophilus  800 VVRTVRLRR-----------------------------------------------------------------------
E. coli          928 VIDVQVFTRDGVEKDKRALEIEEMQLKQAKKDLSEELQILEAGLFSRIAVLVAGGVEAEKLDKLPRDRWLELGLTDEEK
0_Nostoc         782 VVDV-RLFT-----------------------------------------------------------------------
1_Litchi         785 VIDV-RWIQ-----------------------------------------------------------------------
2_Arabidopsis    787 VIDV-RWVQ-----------------------------------------------------------------------
3_Gossypium      785 VIDV-RWVQ-----------------------------------------------------------------------
5_Ricinus        785 VIDV-RWVQ-----------------------------------------------------------------------
6_Rosa           785 VIDV-RWIQ-----------------------------------------------------------------------
9_Cucumis        803 VIDV-RWIQ-----------------------------------------------------------------------
11_Nicotiana     785 VIDV-RWIQ-----------------------------------------------------------------------
13_Syringa       785 VIDV-RWIQ-----------------------------------------------------------------------
18_Liquidambar   785 VIDV-RWIQ-----------------------------------------------------------------------
19_Papaver       785 VIDV-RWIQ-----------------------------------------------------------------------
20_Ananas        787 VIDV-RWIQ-----------------------------------------------------------------------
28_Liriodendron  794 VIDV-RWIQ-----------------------------------------------------------------------
30_Magnolia      794 VIDV-RWIQ-----------------------------------------------------------------------
32_Nymphaea      786 VIDV-RWGQ-----------------------------------------------------------------------
33_Amborella     787 VIDV-RWIQ-----------------------------------------------------------------------
35_Picea         791 VIDV-RWIN-----------------------------------------------------------------------
44_Ginkgo        793 VIDV-RWIH-----------------------------------------------------------------------
51_Physcomitrium 800 VIDV-IWIS-----------------------------------------------------------------------
```

```
                                                          β-flap
                                                                      βa14:V823-G894
                                                                  βb14:L815-G894
T.thermophilus   809 ----------------------------GDPGVELKPGVREVVRVYVAQKRKLQVGDKLANRHGNKGVVAKILPVEDMPH
E.coli          1008 QNQLEQLAEQYDELKHEFEKKLEAKRRKITQGDDLAPGVLKIVKVYLAVKRRIQPGDKMAGRHGNKGVISKINPIEDMPY
0_Nostoc         790 ----------------------------REQGDELPPGANMVVRVYVAQKRKIQVGDKMAGRHGNKGIISRILPAEDMPY
1_Litchi         793 ----------------------------KKGGSSY---NPETIRVYISQKREIKVGDKVAGRHGNKGIVSKILPRQDMPY
2_Arabidopsis    795 ----------------------------KKGGSSY---NPEIIRVYISQKREIKVGDKVAGRHGNKGIISKILPRQDMPY
3_Gossypium      793 ----------------------------KKGGSSY---NPETIRVYISQKREIKVGDKVAGRHGNKGIISKILPRQDMPY
5_Ricinus        793 ----------------------------KKGGSSY---NPETIRIYILQKREIKVGDKVAGRHGNKGIISKILPRQDMPY
6_Rosa           793 ----------------------------KKGGSSY---NPETIRVYISQKREIKVGDKVAGRHGNKGIISKILPRQDMPY
9_Cucumis        811 ----------------------------KKGGSSY---NPEMIRVYISQKREIKVGDKVAGRHGNKGIVSKILPREDMPY
11_Nicotiana     793 ----------------------------KRGGSSY---NPETIRVYILQKREIKVGDKVAGRHGNKGIISKILPRQDMPY
13_Syringa       793 ----------------------------KRGGSSY---NPETIRIYISQKREIKVGDKVAGRHGNKGIISKILPRQDMPY
18_Liquidambar   793 ----------------------------KKGGSSY---NPEMIRVYISQKREIKVGDKVAGRHGNKGIISKILPRQDMPY
19_Papaver       793 ----------------------------KKGGSSY---NPEMIRVYISQKREIKVGDKVAGRHGNKGIISKILPRQDMPY
20_Ananas        795 ----------------------------KKDGSSY---NSEMICVYILQKREIKVGDKVAGRHGNKGIISKILPRQDMPY
28_Liriodendron  802 ----------------------------KKGGSSY---NPEMIRVYISQKREIKVGDKVAGRHGNKGIISKILPRQDMPY
30_Magnolia      802 ----------------------------KKGGSSY---NPEMIRVYISQKREIKVGDKVAGRHGNKGIISKILPRQDMPY
32_Nymphaea      794 ----------------------------KKGGSID---NPEMIRVYISQKRKIKVGDKVAGRHGNKGIISKILPRQDMPY
33_Amborella     795 ----------------------------KKGASSY---NPEKIRVYISQKREIKVGDKVAGRHGNKGIISKILPRQDMPY
35_Picea         799 ----------------------------RVDDSGD---NAETVHVYISQKRKIQVGDKVAGRHGNKGIISIVLPRQDMPY
44_Ginkgo        801 ----------------------------EEENSGD---NAETVHVYILQKRKIQVGDKVAGRHGNKGIISKVLPRQDMPY
51_Physcomitrium 808 ----------------------------KKENSSN---YEKIIHVYIAQKRKIQVGDKVAGRHGNKGIISKILPRQDMPY
```

```
                                        βa14:V823-G894
                                βb14:L815-G894          βb15:R900-D907 (T. Th)
T.thermophilus   861 LPDGTPVDVILNPLGVPSRMNLGQILETHLGLAGYFLGQRYISPIFDGAKEPEIKELLAQAFEVYFGKRKGEGFGVDKRE
E.coli          1088 DENGTPVDIVLNPLGVPSRMNIGQILETHLGMAAKGIGDKINAMLKQQQEVAKLREFIQRAYDLGADVRQKVDLST-FSD
0_Nostoc         842 LPDGSPVDIVLNPLGVPSRMNVGQVFECLLGWAGHTLGVRFKITPFDEMYGEESSRRIVHG-KLQE--------------
1_Litchi         842 LQDGRPVDMVFNPLGVPSRMNVGQIFECSLGLAGGLLNRHYRIAPFDERYEQEASRKLVFS-ELYE--------------
2_Arabidopsis    844 LQDGRPVDMVFNPLGVPSRMNVGQIFECSLGLAGSLLDRHYRIAPFDERYEQEASRKLVFS-ELYE--------------
3_Gossypium      842 LQDGRPVDMVFNPLGVPSRMNVGQLFECSLGLAGSLLDRHYRIAPFDERYEQEASRKLVFS-ELYQ--------------
5_Ricinus        842 LQDGRPVDMVFNPLGVPSRMNVGQIFECSLGLAGGLLDRHYRIAPFDERYEQEASRKLVFS-ELYE--------------
6_Rosa           842 LQDGRPVDMVFNPLGVPSRMNVGQIFECSLGLAGGLLDRHYRIAPFDERYEQEASRKLVFS-ELYE--------------
9_Cucumis        860 LQNGRPVDMVFNPLGVPSRMNVGQIFECSLGLAGSLLDRHYRIAPFDERYEQEASRKLVFS-ELYE--------------
11_Nicotiana     842 LQDGRSVDMVFNPLGVPSRMNVGQIFECSLGLAGSLLDRHYRIVPFDERYEQEASRKLVFS-ELYE--------------
13_Syringa       842 LQDGRPVDMVFNPLGVPSRMNVGQIFECSLGLAGGLLDRHYRIAPFDERYEQEASRKLVFS-ELYE--------------
18_Liquidambar   842 VQDGRPVDMVFNPLGVPSRMNVGQIFECSLGLAGGLLDRHYRIAPFDERYEQEASRKLVFS-ELYE--------------
19_Papaver       842 LQDGTPVDMVFNPLGVPSRMNVGQIFECSLGLAGDLLDRHYRIAPFDERYEQEASRKLVFS-ELYE--------------
20_Ananas        844 LQDGTPVDMVFNPLGVPSRMNVGQIFECSLGLAGNLLKRHYRIAPFDERYEQEASRKLVFS-ELYE--------------
28_Liriodendron  851 LQDGTPVDMVFNPLGVPSRMNVGQIFECSLGLAGDLLDRHYRIAPFDERYEQEASRKLVFS-ELYS--------------
30_Magnolia      851 LQDGTPVDMVFNPLGVPSRMNVGQIFECSLGLAGDLLDRHYRIAPFDERYEQEASRKLVFS-ELYS--------------
32_Nymphaea      843 LQDGTPVDMVFNPLGVPSRMNVGQMFECSLGLAGDLLGRHYRITPFDERYEQEASRKLVFS-ELYE--------------
33_Amborella     844 LQDGTPVDMVFNPLGVPSRMNVGQIFECSLGLAGDLLDRHYRITPFDERYEQEASRKLVFP-ELYE--------------
35_Picea         848 LQNGIPVDMVLNPLGVPSRMNVGQIFECLPGLAGNSMNKHYRITPFDERYEREASRKLVFP-ELYK--------------
44_Ginkgo        850 SQNGTPVDMVFNPLGVPSRMNVGQIFECLPGLAGNPMNKHYRITPFGERYEREASRKLVFP-ELYR--------------
51_Physcomitrium 857 LQDGTPIDMVLSPLGVPSRMNVGQIFECLLGLAGYFLGKHYRITPFDEKYEREASRKLVFS-ELYK--------------
```

```
                                                                   βa15:G970-I1071
                          βb15:P1181-D1188 (E. coli)               βb16:G970-Q1100
T.thermophilus    941 VEVLRRAEKLGLV------------TPGKTPEEQLKELFLQGKVVLYDGRTGEPIEGPIVVGQMFIMKLYHMVEDKMHAR
E.coli           1167 EEVMRLAENLRKGMPIATPVFDGAKEAEIKELLKLGDLPTSGQIRLYDGRTGEQFERPVTVGYMYMLKLNHLVDDKMHAR
0_Nostoc          907 ------------------------ARDETSKDWVYNPDDPGKIMLFDGRTGEAFDRPITVGVAYMLKLVHLVDDKIHAR
1_Litchi          907 ------------------------AGKQTANPWIFEPEYPGKSRIFDGRTGDPFEQPVIIGKPYILKLIHQVDDKIHGR
2_Arabidopsis     909 ------------------------ASKQTANPWVFEPEYPGKSRIFDGRTGDPFEQPVIIGKPYILKLIHQVDDKIHGR
3_Gossypium       907 ------------------------ASKQTANPWVFEPEYPGKSRIFDGRTGGPFEQPVIIGKPYILKLIHQVDDKIHGR
5_Ricinus         907 ------------------------ASKQTANPWVFEPEYPGKSRIFDGRTGDPFEQPVIIGKPYILKLIHQVDDKIHGR
6_Rosa            907 ------------------------ASKQTANPWVFEPEYPGKSRIFDGRTGDPFEQPVIIGKPYILKLIHQVDDKIHGR
9_Cucumis         925 ------------------------ASKQTASPWVFEPEYPGKSRIFDGRTGNPFEQPVIIGKPYILKLIHQVDDKIHGR
11_Nicotiana      907 ------------------------ASKQTANPWVFEPEYPGKSRIFDGRTGNPFEQPVIIGKPYILKLIHQVDDKIHGR
13_Syringa        907 ------------------------ASKQTANPWVFEPEYPGKSRIFDGRSGNPFEQPVLIGKPYILKLIHQVDDKIHGR
18_Liquidambar    907 ------------------------ASKQTANPWVFEPEYPGKSRIFDGRTGDPFEQPVIIGKPYILKLIHQVDDKIHGR
19_Papaver        907 ------------------------ASKQTANPWVFEPEYPGKSRIFDGRTGDPFEQPVIIGKSYILKLIHQVDDKIHGR
20_Ananas         909 ------------------------ASKQTRNPWVFEPEYPGKSRIFDGRTGDPFEQPVLMGKSYILKLIHQVDDKIHGR
28_Liriodendron   916 ------------------------ASKQTANPWVFEPECPGKSRIFDGRTGDPFEQPVIIGKSYILKLIHQVDDKIHGR
30_Magnolia       916 ------------------------ASKQTANPWVFEPECPGKSRIFDGRTGDPFEQPVIIGKSYILKLIHQVDDKIHGR
32_Nymphaea       908 ------------------------ASKQTANPWVFEPEYPGKSRIFDGRTGDPFEQPVIIGKSYMLKLIHQVDDKIHGR
33_Amborella      909 ------------------------ASKRTANPWVFEPEYPGKSRIFDGRTGDPFEQPVIIGKSYMLKLIHQVDDKIHGR
35_Picea          913 ------------------------ASEQTANPWVFEPDHPGKHRLIDGRTGDVFEQPVTIGKAYMSKLSHQVDDKIHAR
44_Ginkgo         915 ------------------------ASEQTANPWVFEPDHPGKNRLIDGRTGDLFEQPVTIGKAYIPKLIHQVDDKIHAR
51_Physcomitrium  922 ------------------------ASKKTGNLWLFEPENPGKSRLLNGRTGEIFEQAVTVGKAYMLKLIHQVDDKIHAR

                         Switch-3                                      Clamp
                                     βa15:G970-I1071                             βa16:S1080-L1097
                                             βb16:G970-Q1100
T.thermophilus   1009 STGPYSLITQQPLGGKAQFGGQRFGEMEVWALEAYGAAHTLQEMLTLKSDDIEGRNAAYEAIIKGEDVPEPS-VPESFRV
E.coli           1247 STGSYSLVTQQPLGGKAQFGGQRFGEMEVWALEAYGAAYTLQEMLTVKSDDVNGRTKMYKNIVDGNHQMEPG-MPESFNV
0_Nostoc          962 STGPYSLVTQQPLGGKAQQGGQRFGEMEVWALEAFGAAYTLQELLTVKSDDMQGRNEALNAIVKGKAIPRPG-TPESFKV
1_Litchi          962 SSGHYALVTQQPLRGRSKQGGQRVGEMEVWALEGFGVAHILQEMLTYKSDHIRARQEVLGTTIIGGTIHKPEDAPESFRL
2_Arabidopsis     964 SSGHYALVTQQPLRGRSKQGGQRVGEMEVWALEGFGVAHILQEMLTYKSDHIRARQEVLGTTIIGGTIPKPEDAPESFRL
3_Gossypium       962 SSGHYALVTQQPLRGRSKQGGQRVGEMEVWALEGFGVAHILQEMLTYKSDHIRARQEVLGTTIIGGTIPKPEDAPESFRL
5_Ricinus         962 SSGHYALVTQQPLRGRAKQGGQRVGEMEVWALEGFGVSHILQEMLTYKSDHIRARQEVLGTTIIGGTIPKPEDAPESFRL
6_Rosa            962 SSGHYALVTQQPLRGRAKQGGQRVGEMEVWALEGFGVAHILQEMLTYKSDHIRARQEVLGTTIIGGTIPQPADAPESFRL
9_Cucumis         980 SSGHYALVTQQPLRGRAKQGGQRVGEMEVWALEGFGVAHILQEMLTYKSDHIRARQEVLGTTIIGGTIPKPEDTPESFRL
11_Nicotiana      962 SSGHYALVTQQPLRGRAKQGGQRVGEMEVWALEGFGVAHILQEMLTYKSDHIRARQEVLGTTIIGGTIPNPEDAPESFRL
13_Syringa        962 SSGHYALVTQQPLRGRAKQGGQRVGEMEVWALEGFGVAHILQEMLTYKSDHIRARQEVLGTTIIGGTIPNPEDAPESFRL
18_Liquidambar    962 SSGHYALVTQQPLRGRAKQGGQRVGEMEVWALEGFGVAHILQEMLTYKSDHIRARQEVLGTTIIGGTIPNPKDAPESFRL
19_Papaver        962 SSGHYALVTQQPLRGRAKQGGQRVGEMEVWALEGFGVAHILQEMLTYKSDHIRARQEVLGTTIIGGTIPKPEDAPESFRL
20_Ananas         964 SSGHYALVTQQPLRGRAKQGGQRVGEMEVWALEGFGVAHILQEMLTYKSDHIRARQEVLGATIIGGRVPNPEDAPESFRL
28_Liriodendron   971 SSGHYALVTQQPLRGRAKQGGQRVGEMEVWALEGFGVAHISQEMLTYKSDHIRARQEVLGTTIIGGTIPNPEDAPESFRL
30_Magnolia       971 SSGHYALVTQQPLRGRAKQGGQRVGEMEVWALEGFGVAHISQEMLTYKSDHIRARQEVLGTTIIGGTIPNPEDAPESFRL
32_Nymphaea       963 SSGHYALVTQQPLRGRAKQGGQRVGEMEVWALEGFGVAHILQEMLTYKSDHIRARQEVLGTTIVGGTIPNPEGAPESFRL
33_Amborella      964 SSGHYALVTQQPLRGRAKQGGQRVGEMEVWALEGFGVAHILQEMLTYKSDHIRARQELLGTTIVGGTIPKPEGAPESFRL
35_Picea          968 SSGPYARVTQQPLRGKSKRGGQRIGEMEVWALEGFGVAYILQEMLTLKSDHIRTRNEVLGAIITGGPIPKPDTAPESFRL
44_Ginkgo         970 SSGPYALVTQQPLRGKSKRGGQRVGEMEVWALEGSGVAYISQEMLTLKSDHIIARHEVLGAIITGEPIPKPGTVPESFRL
51_Physcomitrium  977 SSGPYALVTQQPLRGRSRRGGQRVGEMEVWALEGFGVAYILQEMLTIKSDHIHARYEVLGAIITGEPIPKPKTAPESFLL

                                  Clamp

                     βb16:G970-Q1100
T.thermophilus   1088 LVKELQALALDVQTLDE--KDNP------VDIFEGLASKR------------------
E.coli           1326 LLKEIRSLGINIELEDE-----------------------------------------
0_Nostoc         1041 LMRELQSLGLDIAVHKVETQADGSSLDVEVDLMADQSARRTPPRPTYESLSRESLEDDE
1_Litchi         1042 LVRELRSLALELNHFLVSEKNFQINK-KEA----------------------------
2_Arabidopsis    1044 LVRELRSLALELNHFLVSEKNFQINR-KEV----------------------------
3_Gossypium      1042 LVRELRSLALELNHFLVSEKNFQINR-KEA----------------------------
5_Ricinus        1042 LVRELRSLALELNHFLVSEKNFQMNR-KEA----------------------------
6_Rosa           1042 LVRELRSLALELNHFLVSEKNFQINR-KEA----------------------------
9_Cucumis        1060 LVRELRSLALELNHFLVSEKNFQINR-KEA----------------------------
11_Nicotiana     1042 LVRELRSLALELNHFLVSEKNFQINR-KEA----------------------------
13_Syringa       1042 LVRELRSLALELNHFLVSEKNFQINR-KEA----------------------------
18_Liquidambar   1042 LVRELRSLALELNHFLVSEKNFQINR-KEA----------------------------
19_Papaver       1042 LVRELRSLALELNHFLVSEKNFQIQR-KEA----------------------------
20_Ananas        1044 LVRELRSLALELNHFLVSEKNFQINR-KEA----------------------------
28_Liriodendron  1051 LVRELRSLALELNHFLVSEKNFQINR-KEA----------------------------
30_Magnolia      1051 LVRELRSLALELNHFLVSEKNFQINR-KEA----------------------------
32_Nymphaea      1043 LVRELRSLSLELNHFLVSEKNFQINR-KEV----------------------------
33_Amborella     1044 LVRELRSLALELKHFLVSEKNFQINR-KEA----------------------------
35_Picea         1048 LIRELRSLALELNHAIISEKDFQIDR-EEV----------------------------
44_Ginkgo        1050 LVRELRSLAPELNHAIISEKDFQIDK-KEV----------------------------
51_Physcomitrium 1057 LVRELRSLSLELDHAVIFEKNLNIKF-KDV----------------------------
```

**Fig. S5.** Sequence alignment of the β' subunits from PEP of angiosperms with those of the RNAPs from *E. coli*, *T. thermophilus* and Nostoc. The residues conserved more than 50 % are in red, those mutated in similar residues are in blue. The strictly conserved residues described by Lane & Darst (Lane & Darst, 2010) are highlighted in gray. The blue triangles show mutations observed among the strictly conserved residues described (Lane & Darst, 2010). The non-conservative mutations, at least three in a row in the β or β' domain in *E. coli* and *T. thermophilus*, are high-lighted in green and displayed on the *E. coli* structure (PDB entry: 6GH5). Those colored in orange are nearby to the DNA, those in green are located at the surface of the subunits. The domains described for all-RNA polymerase (a) and the bRNAPs (b) are also given and highlighted in yellow and cyan respectively. The name of the RNAP domains are also given and highlighted in purple and green (Lane & Darst, 2010; Sutherland & Murakami, 2018).

```
                                           Clamp                     Zipper          Clamp
                                  β’a1:S14-S22                          β’a2:D42-G51
                                                        β’b1:A13-L135
T. thermophilus    1 ------------MKKEVRKVRIALASPEKIRSWS---------YGEVEKPETINYRTLKPERDGLFDERIFGPIKDYECA
E. coli            1 MKDLLKFLKAQTKTEEFDAIKIALASPDMIRSWS---------FGEVKKPETINYRTFKPERDGLFCARIFGPVKDYECL
0_Nostoc           1 --------MRPAQTNQFDYVKIGLASPERIRQWGERTLPNGQVVGEVTKPETINYRTLKPEMDGLFCERIFGPAKDWECH
1_Litchi           1 -------MI---DRYKHQQLRIGSVSPQQISAWTKKILPNGEIVGEVTKPYTFHYKTNKPEKDGLFCERIFGPIKSGICA
2_Arabidopsis      1 -------MI---DRYKHQQLRIGLVSPQQISAWATKIIPNGEIVGEVTKPYTFHYKTNKPEKDGLFCERIFGPIKSGICA
3_Gossypium        1 -------MI---DRYKHQQLRIGSVSPQQISAWAKKILPNGETVGEVTKPYTFHYKTNKPEKDGLFCERIFGPIKSGICA
5_Ricinus          1 -------MI---DRYKHQQLRIGSVSPQQISAWAKKILPNGEIVGEVTKPYTFHYKTNKPEKGGLFCERIFGPIKSGICA
6_Rosa             1 MNQNFSSMI---DRYKHQQLRIGLVSPQQISAWAQKILPNGEIVGEVTKPYTFHYKTNKPEKDGLFCERIFGPIKSGICA
9_Cucumis          1 MNQKIFSMI---DRYKHQQLRIGLVSPQQISAWANKTLPNGEIVGEVTKPYTFHYKTNKPEKDGLFCERIFGPIKSGICA
11_Nicotiana       1 MNNNFSSMI---DRYKHQQLRIGSVSPQQISAWATKILPNGEIVGEVTKPYTFHYKTNKPEKDGLFCERIFGPIKSGICA
13_Syringa         1 MNQNFSSMI---DRYKHQQLRIGLVSPQQISAWATKILPNGEIVGEVIKPYTFHYKTNKPEKDGLFCERIFGPIKSGICA
18_Liquidambar     1 -------MI---DRYKHQQLRIGSVSPQQISAWANKILPNGEIVGEVTKPYTFHYKTNKPEKGGLFCERIFGPIKSGICA
19_Papaver         1 -------MI---DQYKHQHLRIGSVSPEQISAWAKKILPNGEVVGEVTKPYTFHYKTNKPEKDGLFCERIFGPIKSGICA
20_Ananas          1 -------MI---DQYKHQQLRIGPVSPQQIKAWANKILPNGEIVGEVTKPYTFHYKTNKPEKDGLFCERISGPIKSGICA
28_Liriodendron    1 -------MI---DRYKHQQLRIGSVSPQQISAWANKILPNGEIVGEVTKPYTFHYKTNKPEKDGLFCERIFGPIKSGICA
30_Magnolia        1 -------MI---DRYKHQQLRIGSVSPQQISAWANKILPNGEIVGEVTKPYTFHYKTNKPEKDGLFCERIFGPIKSGICA
32_Nymphaea        1 MNQNFSSMI---DQYKHQQLRIGLVSPKQIRAWANKILPNGEIVGEVTKPYTFHYKTNKPEKDGLFCERIFGPIKSGICA
33_Amborella       1 -------MI---DRYKHQQLRIGLVSPQQITAWANKILPNGEMVGEVTKPYTFHYKSNKPEKDGLFCERIFGPIKSGICA
35_Picea           1 -------MI---DQNKHQQLRIGLASPEQICAWSEKILPNGEIVGQVTKPYTLHYETNKPERDGSFCERIFGPIKSRVCS
44_Ginkgo          1 MNRNLSFTI---ARDKHQQLRIGLASPEKICAWSEKILPNGEIVGQVTKPHTSHYKTNEPEKDGLFCERIFGPIKSGVCA
51_Physcomitrium   1 -------MI---HREKYHHLRIRLASPEQIRSWAERVLPNGEIVGQVTKPYTLHYKTHKPEKDGLFCERIFGPIKSGICA

                                              β'a1–a6; clamp
                 β’a3:C58-G61                         β’a4:R89-V105      β’a5:S110-Y128
                                                 β’b1:A13-L135
T. thermophilus   60 CGKYKRQR---FEGKVCERCGVEVTKSIVRRYRMGHIELATPAAHIWFVKDVPSKIGTLLDLSATELEQVLYFSKYIVLD
E. coli           72 CGKYKRLK---HRGVICEKCGVEVTQTKVRRERMGHIELASPTAHIWFLKSLPSRIGLLLDMPLRDIERVLYFESYVVIE
0_Nostoc          73 CGKYKRVR---HRGIVCERCGVEVTESRVRRHRMGYIKLAAPVAHVWYLKGIPSYISILLDMPLRDVEQIVYFNSYVVLS
1_Litchi          71 CGNYRVIGDEKEDPQFCEQCGVEFVNSRIRRYQMGYIKLGCPVTHVWYLKRLPSYIANLLDKPLKELEGLVYCD----FS
2_Arabidopsis     71 CGNYRVIGDEKEDPKFCEQCGVEFVDSRIRRYQMGYIKLTCPVTHVWYLKRLPSYIANLLDKPLKELEGLVYCD----FS
3_Gossypium       71 CGNYRVIGNQKEGPKFCEQCGVEFVDSRIRRYQMGYIRLACPVTHVWYLKRLPSYIANLLDKPLKELEGLVYCD----FS
5_Ricinus         71 CGNYRVIRNEKEDQKFCEQCGVEFVDSRIRRYQMGYIKLACPVTHVWYLKRLPSYIANLLDKPLKELEGLVYCD----V-
6_Rosa            78 CGNYRVIGDEKKDPRFCEQCGVEFVDSRIRRYQMGYIKLACPVTHVWYLKRLPSYIANLLDKPLKELEGLVYCD----FS
9_Cucumis         78 CGNYRVIGDKKEDSKFCEQCGVEFVDSRIRRYQMGYIKLACPVTHVWYLKRLPSYIANLLDKPLKELEGLVYCD----FS
11_Nicotiana      78 CGNYRVIGDEKEDPKFCEQCGVEFVDSRIRRYQMGYIKLACPVTHVWYLKRLPSYIANLLDKPLKELEGLVYCD----FS
13_Syringa        78 CGNYRVIGDEKEDPKFCEQCGVEFVDSRIRRYQMGYIKLACPVTHVWYLKRLPSYIANLLDKPLKELEGLVYCD----FS
18_Liquidambar    71 CGNYRVIGDEKEDPKFCEQCGVEFVDSRVRRYQMGYIKLACPVTHVWYLKRLPSYIANLLDKPLKELEGLVYCD----FS
19_Papaver        71 CGNYRVIGDEKEDPKFCEQCGVEFVDSRIRRYQMGYIKLACPVTHVWYLKRLPSYIANLLDKPLKELEGLVYCD----VS
20_Ananas         71 CGNYRVIRAEKEDPKFCEQCGVEFVDSRIRRYQMGYIKLACPVTHVWYLKRLPSYIANLLDKPLKQLEGLVYCDVYLDFS
28_Liriodendron   71 CGNYRVIGNEKEDPKFCEQCGVEFVDSRIRRYQMGYIKLACPVTHVWYLKRLPSYIASLLDKPLKELEGLVYCD----FS
30_Magnolia       71 CGNYRVIGNEKEDPKFCEQCGVEFVDSRIRRYQMGYTKLACPVTHVWYLKRLPSYIASLLDKPLKELEGLVYCD----FS
32_Nymphaea       78 CGNYRVIGGEKEEPKFCEQCGVESVDSRIRRYQMGYIKLACPVTHVWYLKRLPSYIANLLDKPLKELEGLVYCD----FS
33_Amborella      71 CGNYRVIGDEKEDPKFCEQCGVESVDSRIRRYQMGYIKLACPVTHVWYLKRLPSYIANLSDRPLKELEGLVYCD----FS
35_Picea          71 CGNSPGIGNEKIDSKFCTQCGVEFVDSRIRRYRMGYIKLACPVAHIWYLKRLPSYIANLLAKTRKELEGPVYCDLF----
44_Ginkgo         78 CGNSRVIRNEKEDSKFCEQCGVEFVDSRSRRYRMGYIKLACPVVHVWYSKRLPSYIANLLAKPLKELEGPVYCDLF----
51_Physcomitrium  71 CGKYQIIE---KYSKFCEQCGVEFVESRVRRYRMGYIKLACPVTHVWYLKRLPSYIANLLAKPLKELESLVYCDLF----

                         Clamp
T. thermophilus  137 PKGAILNGVPVEKRQLLTDEEYRELRYGKQETYPLPPGVDALVKDGEEVVKGQELAPGVVSRLDGVALYRFPRRVRVEYV
E. coli          149 GGMT-----NLERQQILTEEQYLDAL------------------------------------------------------
0_Nostoc         150 AGNAE----TLTYKQLLSEDQWLEIE------------------------------------------------------
1_Litchi         147 FARPIAKKPTFLRLRGLF-------EY-----------------------------------------------------
2_Arabidopsis    147 FARPITKKPTFLRLRGSF-------EY-----------------------------------------------------
3_Gossypium      147 FARPIAKKPTFLRLRGSF-------EY-----------------------------------------------------
5_Ricinus        146 --------------------------------------------------------------------------------
6_Rosa           154 FARPIAKKPTFLRLRGSF-------EY-----------------------------------------------------
9_Cucumis        154 FARPIAKKPTFLRLRGSF-------EY-----------------------------------------------------
11_Nicotiana     154 FARPITKKPTFLRLRGLF-------EY-----------------------------------------------------
13_Syringa       154 FARPITKKPTFLRLRGLF-------EY-----------------------------------------------------
18_Liquidambar   147 FARPIAKKPTFLRLRGSF-------EY-----------------------------------------------------
19_Papaver       147 FARTVAKKPTFLRLRGSF-------EY-----------------------------------------------------
20_Ananas        151 FARPIAKKPTFLRLRGSF-------EY-----------------------------------------------------
28_Liriodendron  147 FARPIAKKPTFLRLRGSF-------ES-----------------------------------------------------
30_Magnolia      147 FARPIAKKPTFLRLRGSF-------ES-----------------------------------------------------
32_Nymphaea      154 FARPIAKKPTFLRLRGSF-------EY-----------------------------------------------------
33_Amborella     147 FARPIAKKPTFLRLRGSF-------EY-----------------------------------------------------
35_Picea         147 IARPIANKPTLLRSRGTF-------NY-----------------------------------------------------
44_Ginkgo        154 IARPIANKPTSLRSRGTF-------KY-----------------------------------------------------
51_Physcomitrium 144 LARPISKKPILLKLRGLF-------KY-----------------------------------------------------
```

```
T. thermophilus   217 KKERAGLRLPLAAWVEKEAYKPGEILAELPEPYLFRAEEEGVVELKELEEGAFLVLRREDEPVATYFLPVGMTPLVVHGE
E. coli           170 --------------------------------------------------------------------------------
0_Nostoc          172 --------------------------------------------------------------------------------
1_Litchi          167 ---------EIQSWKYSI-------------PLFFTT--------------------------------QGFD-------
2_Arabidopsis     167 ---------EIQSWKYSI-------------PLFFTT--------------------------------QGFD-------
3_Gossypium       167 ---------EIQSWKYSI-------------PLFFTT--------------------------------QGFD-------
5_Ricinus         146 --------------KYSI-------------PLFFTA--------------------------------QGFD-------
6_Rosa            174 ---------EIQSWKYSI-------------PLFFTT--------------------------------PGFD-------
9_Cucumis         174 ---------EIQSWKYSI-------------PLFFTT--------------------------------QGFD-------
11_Nicotiana      174 ---------EIQSWKYSI-------------PLFFTT--------------------------------QGFD-------
13_Syringa        174 ---------EIQSWKYSI-------------PLFFTT--------------------------------QGFD-------
18_Liquidambar    167 ---------EIQSWKYSI-------------PLFFTT--------------------------------QGFD-------
19_Papaver        167 ---------EIQSWKYSI-------------PLFFTT--------------------------------QGFD-------
20_Ananas         171 ---------EIQSRNYSI-------------PLFFTT--------------------------------SGFE-------
28_Liriodendron   167 ---------EIQSRKYSI-------------PLFFTT--------------------------------QDFD-------
30_Magnolia       167 ---------EIQSRKYSI-------------PLFFTT--------------------------------QGFD-------
32_Nymphaea       174 ---------EIQSRKYSI-------------PLFFTT--------------------------------QGFD-------
33_Amborella      167 ---------EIQSRKYSI-------------PLFFTT--------------------------------QCFN-------
35_Picea          167 ---------EIQSWRDII-------------PHYLSAR-------------------------SHYLFARGSG-------
44_Ginkgo         174 ---------DIQSWGDIL-------------PHYLSA--------------------------------QGFG-------
51_Physcomitrium  164 ---------EDQSWREIF-------------PRYFSS--------------------------------RGFE-------

T. thermophilus   297 IVEKGQPLAEAKGLLRMPRQVRAAQVEAEEEGETVYLTLFLEWTEPKDYRVQPHMNVVVPEGARVEAGDKIVAAIDPEEE
E. coli           170 ------------------------------------------------------------------------------EE
0_Nostoc          172 ------------------------------------------------------------------------------DQ
1_Litchi          186 --------------------------------------------------------------------------------
2_Arabidopsis     186 --------------------------------------------------------------------------------
3_Gossypium       186 --------------------------------------------------------------------------------
5_Ricinus         160 --------------------------------------------------------------------------------
6_Rosa            193 --------------------------------------------------------------------------------
9_Cucumis         193 --------------------------------------------------------------------------------
11_Nicotiana      193 --------------------------------------------------------------------------------
13_Syringa        193 --------------------------------------------------------------------------------
18_Liquidambar    186 --------------------------------------------------------------------------------
19_Papaver        186 --------------------------------------------------------------------------------
20_Ananas         190 --------------------------------------------------------------------------------
28_Liriodendron   186 --------------------------------------------------------------------------------
30_Magnolia       186 --------------------------------------------------------------------------------
32_Nymphaea       193 --------------------------------------------------------------------------------
33_Amborella      186 --------------------------------------------------------------------------------
35_Picea          193 --------------------------------------------------------------------------------
44_Ginkgo         193 --------------------------------------------------------------------------------
51_Physcomitrium  183 --------------------------------------------------------------------------------

T. thermophilus   377 VIAEAEGVVHLHEPASILVVKARVYPFEDDVEVSTGDRVAPGDVLADGGKVKSDVYGRVEVDLVRNVVRVVESYDIDARM
E. coli           172 -----------------------F------------------------------------------------GDEFDAKM
0_Nostoc          174 -----------------------IYS-E-----------------------------------------DSVLQGVEVGI
1_Litchi          186 -------------------------TFR----------------------------------------------NREIST
2_Arabidopsis     186 -------------------------IFR----------------------------------------------NREIST
3_Gossypium       186 -------------------------TFR----------------------------------------------SREIST
5_Ricinus         160 -------------------------TFR----------------------------------------------NREIST
6_Rosa            193 -------------------------TFR----------------------------------------------NREIST
9_Cucumis         193 -------------------------TFR----------------------------------------------NREIST
11_Nicotiana      193 -------------------------TFR----------------------------------------------NREIST
13_Syringa        193 -------------------------TFR----------------------------------------------NREIST
18_Liquidambar    186 -------------------------TFR----------------------------------------------NREIST
19_Papaver        186 -------------------------TFR----------------------------------------------NREIST
20_Ananas         190 -------------------------TFR----------------------------------------------NREIST
28_Liriodendron   186 -------------------------TFR----------------------------------------------NREIST
30_Magnolia       186 -------------------------TFR----------------------------------------------NREIST
32_Nymphaea       193 -------------------------TFR----------------------------------------------NREIST
33_Amborella      186 -------------------------LFR----------------------------------------------NREIST
35_Picea          193 -------------------------TFQ----------------------------------------------EREIAT
44_Ginkgo         193 -------------------------AFQ----------------------------------------------NREIAT
51_Physcomitrium  183 -------------------------AFQ----------------------------------------------NKEIAT
```

```
Clamp                                    Clamp
β'a6:G457-L470                           β'a7:R500-F502
β'b2:A454-M481                      β'b2:A454-M481      β'b3:K494-S596
T. thermophilus  457 GAEAIQQLLKELDLE----------------------ALEKELLEEMKHP-SRARRAKARKRLEVVRAFLDSGNRPEWMI
E. coli          181 GAEAIQALLKSMDLE----------------------QECEQLREELNETNSETKRKKLTKRIKLLEAFVQSGNKPEWMI
0_Nostoc         189 GAEALLRLLADINLE----------------------QEAESLREEIGNAKG-QKRAKLIKRLRVIDNFIATGSKPEWMV
1_Litchi         195 GAVAIREQLADLDLQIIIDYSLVDWKELG--------EEGP-TGNEWEDRKIGRRRDFLVRRIELAKHFLRTNIEPEWMV
2_Arabidopsis    195 GAGAIREQLADLDLRIIIENSLVEWKQLG--------EEGP-TGNEWEDRKIVRRKDFLVRRMELAKHFIRTNIEPEWMV
3_Gossypium      195 GAGAIREQLADLDLRILIDYSVVEWKELG--------EEGL-TGNEWEDRKIGRRKDFLVRRMELAKHFIRTNIEPEWMV
5_Ricinus        169 GAGAIREQLADLDLRIIIDYSSVEWKELG--------EEGP-TGNEWEDRKVGRRKDFLVRRVELAKHFIRTNIEPEWMV
6_Rosa           202 GAGAIREQLADLDLRIIIDYSLLEWKELG--------EEGS-TGNEWEDRKVGRRKDFLVRRMELAKHFIRTNIEPEWMV
9_Cucumis        202 GAGAIREQLADLDLRLIIDYSLVEWKELG--------EEGP-ACNEWEDRKVGRRKDFLVRRMELAKHFIRTNIEPEWMV
11_Nicotiana     202 GAGAIREQLADLDLRIIIENSLVEWEELG--------EEGH-TGNEWEDRKVGRRKDFLVRRVELAKHFIRTNIEPEWMV
13_Syringa       202 GAGAIREQLADLDLRIILDNSLVEWKELG--------EEGP-TGNEWEDRKVGRRKDFLVRRMELAKHFIRTNIEPEWMV
18_Liquidambar   195 GAGAIREQLADLDLRIIIDYSLVEWKELG--------EEGS-AGNEWEDRKIGRRKDFLVRRMELAKHFIRTNIEPEWMV
19_Papaver       195 GASAIREQLADLDLRLIIDCSLVEWKELG--------EEGP-TGNEWEDRKIGRRKDFLVRRMELAKHFIRTNVEAEWMV
20_Ananas        199 GAGAIREQLADSDLRIIIDNSLAEWKELG--------DEGS-TGNEWEDRKIRRRKDFLVRRMELAKHLIRTNVEPEWMV
28_Liriodendron  195 GAGAIKEQLADPDLRIITDHSLVEWKELG--------EEGSADGNEWEDRKIGRRKDFLVRRMELAKHFIRTNVEPERMV
30_Magnolia      195 GAGAIREQLADPDLRIITDHSLVEWKELG--------EEGSADGNEWEDRKIGRRKDFLVRRIELAKHFIRTNVEPERMV
32_Nymphaea      202 GATAIREQLADLDLRIIIDRSLVEWKELG--------EEGS-TGNDWEDRKIGRRKDFLVRRMELAKHFLRTNVEPEWMV
33_Amborella     195 GAGAIREQLADPDLRIITDRSLVEWKELG--------EERS-AENEWEDKKIVRRKDFLVRRMELAKHLLRTNVEPERMV
35_Picea         202 GGDAIREQLTGLDLQIIIDRSHMEWKNLVELKWNRLEEDQESTVDGWEDETIRRKDFLVGRMKLAKHFLRTNIEPKWMV
44_Ginkgo        202 GGDAIREQLAGPDLRILMANSYMEWKILE---------EQKSTGNEWEDEKIQRRKDFSVRRMELAKHFIQTNIEPEWMV
51_Physcomitrium 192 GGDAIKKQLSNLDLQGVLDYAYIEWKELV---------EQKSTGNEWEDRKIQRRKDLLVRRIKLAKQFLQTNIKPEWMV
                                 Lid        β'coiled-coil region (ccr)        Rudder
```

```
Clamp
β'a8:P509-V530        β'a9:T537-L554         β'a10:K571-D583
β'b3: K494-S596
T. thermophilus  514 LEAVPVLPPDLRPMVQVDGGRFAT-SDLNDLYRRLINRNNRLKKLLAQG--APEIIIRNEKRMLQEAVDALLDNGRRGAP
E. coli          239 LTVLPVLPPDLRPLVPLDGGRFAT-SDLNDLYRRVINRNNRLKRLLDLA--APDIIVRNEKRMLQEAVDALLDNGRRGRA
0_Nostoc         246 MAVIPVIPPDLRPMVQLDGGRFAT-SDLNDLYRRVINRNNRLARLQEIL--APEIIVRNEKRMLQEAVDALIDNGRRGRT
1_Litchi         266 LCLLPVLPPELRPIIQIEGGKLMS-SDINELYRRVIYRNNTLTDLLTTSRSTPGELVMCQEKLVQEAVDTLLDNGIRGQP
2_Arabidopsis    266 LCLLPVLPPELRPIIQIEGGKLMS-SDINELYRRVIYRNNTLTDLLTTSRSTPGELVMCQEKLVQEAVDTLLDNGIRGQP
3_Gossypium      266 LCLLPVLPPELRPIIQIDGGKLMS-SDINELYRRVIYRNNTLTDLLTTSRSTPGELVMCQEKLVQEAVDTLLDNGIRGQP
5_Ricinus        240 LCLLPVLPPELRPIIQIDGGKLMS-SDINELYRRVIYRNNTLIDLLTTSRSTPGELVMCQEKLVQEAVDTLLDNGIRGQP
6_Rosa           273 LCLLPVLPPELRPIIQIDGGKLMS-SDINELYRRVIYRNNTLIDLLTTSRSTPGELVMCQEKLVQEAVDTLLDNGIRGQP
9_Cucumis        273 LCLLPVLPPELRPIIQIDGGKLMS-SDINELYRRVIYRNNTLIDLLTTSRSTPGELVMCQEKLVQEAVDTLLDNGIRGQP
11_Nicotiana     273 LCLLPVLPPELRPIIQIDGGKLMS-SDINELYRRVIYRNNTLTDLLTTSRSTPGELVMCQEKLVQEAVDTLLDNGIRGQP
13_Syringa       273 LCLLPVLPPELRPIIQIDGGKLMS-SDINELYRRVIYRNNTLTDLLTTSRSTPGELVMCQEKLVQEAVDTLLDNGIRGQP
18_Liquidambar   266 LCLLPVLPPELRPIIQIDGGKLMS-SDINELYRRVIYRNNTLTDLLTTSRSTPGELVMCQEKLVQEAVDTLLDNGIRGQP
19_Papaver       266 LCLLPVLPPELRPIIQIDGGKLMS-SDINELYRRVIYRNNTLTDLLTTSRSTPGELVMCQEKLVQEAVDTLLDNGIRGQP
20_Ananas        270 LCLLPVLPPELRPIIQIDGGKLMS-SDINELYRRVIYRNNTLTDLLATSRSTPGELVMCQEKLVQEAVDTLLDNGIRGQP
28_Liriodendron  267 LCLLPVLPPELRPIIQIDGGKPMS-SDINELYRRVIYRNNTLTDPLTTSRSTPGESVMCQEKLVQEAVDTLLDNGIRGQP
30_Magnolia      267 LCLLPVLPPELRPIIQIDGGKPMS-SDINELYRRVIYRNNTLTDPLTTSRSTPGESVMCQEKLVQEAVDTLLDNGIRGQP
32_Nymphaea      273 LSLLPVLPPELRPIIQIDGGKSMS-SDINELYRRVIYRNNTLTDLLTTSRSTPGELVMCQEKLVQEAVDTLLDNGIRGQP
33_Amborella     266 LCLLPVLPPELRPIIQIDGGKPMS-SDINELYRRVIYRNNTLIDPLTTSRSTPGESVMCQEKLVQEAVDTLLDNGIRGQP
35_Picea         282 LCLLPVLPPEPRPIVQLGEGGLITSSDLNELYRRVINRNNTLTNLLARSGSE--SFVICQKKLIQEAVDALLDNGICGQP
44_Ginkgo        273 LCLLPVLPPEPRPIVQLSEGELIT-SDLNELYRKVIHRNNTLTNLSARSGSAPGGLVICQKKLVQEAVDALLDNGIRGQP
51_Physcomitrium 263 LSLLPVLPPELRPMIELGEGELIT-SDLNELYRRVIYRNNTLIDFLARSGSTPGGLVVCQIRLVQEAVDGLIDNGIRGQP
                     Rudder      ccr   Switch2
```

```
Clamp
β'a11:S602-R674
β'b4:R598-D682
T. thermophilus  591 VTNPGSDRPLRSLTDILSGKQGRFRQNLLGKRVDYSGRSVIVVGPQLKLHQCGLPKRMALELFKPFLLKKMEEKGIAPNV
E. coli          316 ITG-SNKRPLKSLADMIKGKQGRFRQNLLGKRVDYSGRSVITVGPYLRLHQCGLPKKMALELFKPFIYGKLELRGLATTI
0_Nostoc         323 VVG-ANNRPLKSLSDIIEGKQGRFRQNLLGKRVDYSGRSVIVVGPKLKIHQCGLPREMAIELFQPFVINRLIRSGMVNNI
1_Litchi         345 MRD-GHNKVYKSFSDVIEGKEGRFRETLLGKRVDYSGRSVIVVGPSLSLHRCGLPREIAIELFQTFVICSLIRQHLASNI
2_Arabidopsis    345 MRD-GHNKVYKSFSDVIEGKEGRFRETLLGKRVDYSGRSVIVVGPSLSLHRCGLPREIAIELFQTFVIRGLIRQHLASNI
3_Gossypium      345 MRD-GHNKVYKSFSDVIEGKEGRFRETLLGKRVDYSGRSVIVVGPSLSLHRCGLPREIAIELFQTFVIRGLIRQHLAPNI
5_Ricinus        319 MRD-GHNKVYKSFSDVIEGKEGRFRETMLGKRVDYSGRSVIVVGPSLSLHRCGLPREIAIELFQIFVIRGLIRQHLASNI
6_Rosa           352 MRD-GHNKVYKSFSDVIEGKEGRFRETLLGKRVDYSGRSVIVVGPSLSLHRCGLPREIAIELFQTFVIRGLIRQHFASNI
9_Cucumis        352 MRD-GHNKVYKSFSDVIEGKEGRFRETLLGKRVDYSGRSVIVVGPSLSLHRCGLPREIAIELFQTFLIRGLIRQHFASNI
11_Nicotiana     352 MRD-GHNKVYKSFSDVIEGKEGRFRETLLGKRVDYSGRSVIVVGPSLSLHRCGLPREIAIELFQTFVIRGLIRQHLASNI
13_Syringa       352 MRD-GHNKVYKSFSDVIEGKEGRFRETLLGKRVDYSGRSVIVVGPSLSLHRCGLPREIAIELFQTFVIRGLIRQHLASNI
18_Liquidambar   345 MRD-GHNKVYKSFSDVIEGKEGRFRETLLGKRVDYSGRSVIVVGPSLSLHRCGLPREIAIELFQTFVIRGLIRQHLASNI
19_Papaver       345 MRD-GHNKVYKSFSDVIEGKEGRFRETLLGKRVDYSGRSVIVVGPSLSLHRCGLPREIAIELFQTFVIRGLIRQQVASNI
20_Ananas        349 MRD-GHNKVYKSFSDVIEGKEGRFRETLLGKRVDYSGRSVIVVGPSLSLHQCGLPGEIAIELFQTFVIRGLIRQHGVSNI
28_Liriodendron  346 MRD-GHNKVYKSFSDVIEGKEGRFRETLLGKRVDYSGRSVIVVGPSLSLHRCGLPREIAIELFQTFVIRGLIRQHVASNI
30_Magnolia      346 MRD-GHNKVYKSFSDVIEGKEGRFRETLLGKRVDYSGRSVIVVGPSLSLHRCGLPREIAIELFQTFVIRGLIRQHVASNI
32_Nymphaea      352 MRD-GHNKVYKSFSDVIEGKEGRFRETLLGKRVDYSGRSVIVVGPSLSLHQCGLPREIAIELFQTFVIRGLIRQHLASNI
33_Amborella     345 MRD-GHNKVYKSFSDVIEGKEGRFRETLLGKRVDYSGRSVIVVGPSLSLHRCGLPREIAIELFQTFVIRGLIRQHVASNI
35_Picea         360 MRD-SHDRPYKSFSDVIEGKEGRSRENLLGKRVDYSGRSVIVVGPFLSLYQCGLPSEIAIELFQAFVIRSLIGRHIAPNL
44_Ginkgo        352 MKD-SRDRPYKSFSDVIEGKEGRSRENLLGKRVDYSGRSVIVVGPSLPLHQCGLPREIAIELFQAFVIRGPIGRHLAPNL
51_Physcomitrium 342 MKD-SHNRPYKSFSDVIEGKEGRFRENLLGKRVDYSGRSVIVVGPFLSLHQCGLPREMAIELFQAFVIRGLIGRHLAPNL
                                                ▲
```

```
                                                                  β'a12:I695-T793
                                                                  β'b5:D686-T793
T. thermophilus   671 KAARRMLERQRDIKDEVWDALEEVIHGKVVLLNRAPTLHRLGIQAFQPVLVEGQSIQLHPLVCEAFNADFDGDQMAVHVP
E. coli           395 KAAKKMVEREE---AVVWDILDEVIREHPVLLNRAPTLHRLGIQAFEPVLIEGKAIQLHPLVCAAYNADFDGDQMAVHVP
0_Nostoc          402 KAAKKLISRND---PSVWDVLEEVIEGHPVMLNRAPTLHRLGIQSFEPILVEGRAIQLHPLVCPAFNADFDGDQMAVHVP
1_Litchi          424 GVAKSQIRDKG---PIVWEILQEVMQGHPVLLNRAPTLHRLGIQAFQPILVEGRAICLHPLVCKGFNADFDGDQMAVHVP
2_Arabidopsis     424 GVAKSQIREKK---PIVWEILQEVMQGHPVLLNRAPTLHRLGIQSFQPILVEGRTICLHPLVCKGFNADFDGDQMAVHVP
3_Gossypium       424 GVAKSKIREKG---PIVWEILQEVMRGHPVLLNRAPTLHRLGIQAFQPILVEGRAICLHPLVCKGFNADFDGDQMAVHVP
5_Ricinus         398 GVAKSKIREKE---PIVWEILQEVMQGHPVLLNRAPTLHRLGIQAFQPILVEGRAICLHPLVCKGFNADFDGDQMAVHVP
6_Rosa            431 GVAKSKIREKE---PVVWEILQEVMQGHPVLLNRAPTLHRLGIQAFQPILVEGHAICLHPLVCKGFNADFDGDQMAVHVP
9_Cucumis         431 GVAKSKIREKE---PIVWEILQEVMQGHPVLLNRAPTLHRLGIQAFQPILVEGRAICLHPLVCKGFNADFDGDQMAVHVP
11_Nicotiana      431 GVAKSKIREKE---PIVWEILQEVMQGHPVLLNRAPTLHRLGIQAFQPVLVEGRAICLHPLVCKGFNADFDGDQMAVHVP
13_Syringa        431 GVAKSKIREKE---PIVWEILQEVMQGHPVLLNRAPTLHRLGIQAFQPVLVEGRAICLHPLVCKGFNADFDGDQMAVHVP
18_Liquidambar    424 GVAKSKIREKE---PIVWEILQEVMQGHPVLLNRAPTLHRLGIQAFQPVLVEGRAICLHPLVRKGFNADFDGDQMAVHVP
19_Papaver        424 GVAKSKIREKE---PIVWEILQEVMQGHPVLLNRAPTLHRLGIQAFQPILVEGRAICLHPLVCKGFNADFDGDQMAVHVP
20_Ananas         428 GIAKSKIREKE---PIVWEILQEVMQGHPVLLNRAPTLHRLGIQAFQPILVEGRAICLHPLVCKGFNADFDGDQMAVHVP
28_Liriodendron   425 GIAKSKIREKE---PIVWEILQEVMQGHPVLLNRAPTLHRLGIQAFQPILVEGRAICLHPLVRKGFNADFDGDQMAVHVP
30_Magnolia       425 GIAKSKIREKE---PIVWEILQEVMQGHPVLLNRAPTLHRLGIQAFQPILVEGRAICLHPLVRKGFNADFDGDQMAVHVP
32_Nymphaea       431 GLAKSKIREKE---PIVWEILQEVMQGHPVLLNRAPTLHRLGIQAFQPILVEGRAICLHPLVCKGFNADFDGDQMAVHVP
33_Amborella      424 GIAKSKIREKE---PIVWEILQKVMEGHPVLLNRAPTLHRLGIQAFQPILVEGRAICLHPLVRKGFNADFDGDQMAVHVP
35_Picea          439 RAAKSMIRDKG---PIVWEVLQEVMQGHPVLLNRAPTLHRLGIQAFQPILVEGRAIRSHPSVCGGFNADFDGDQMAVHVP
44_Ginkgo         431 RAAKSIIRDKE---PVIWKVLQEVLQGHPVSLNRAPTSHRLGIQAFQPILVEGRVIRLHPSVCGGFNADSDGDQMAVHVP
51_Physcomitrium  421 RAAKSMIQNKE---PIIWKILQEIMQGHPVLLNRAPTLHRLGIQAFQPILIKGRAIRLHPLVCGGFNADFDGDQMAVHIP

                                β'a12:I695-T793
                                β'b5: D686-T793
T. thermophilus   751 LSSFAQAEARIQMLSAHNLLSPASGEPLAKPSRDIILGLYYITQVRKE-K---------------------------KGA
E. coli           472 LTLEAQLEARALMMSTNNILSPANGEPIIVPSQDVVLGLYYMTRDCVNA----------------------------KGE
0_Nostoc          479 LSLESQAEARLLMLASNNILSPATGKPIITPSQDMVLGAYYLTAENPGAT---------------------------KGA
1_Litchi          501 LSLEAQAEARLLMFSHMNLLSPAIGDPISVPTQDMLIGLYVLTSRNLRGICANRYNPWNRRNYQNERIDDN---RYKYMK
2_Arabidopsis     501 LSLEAQAEARLLMFSHMNLLSPAIGDPISVPTQDMLIGLYVLTSGTRRGICANRYNPCNRKNYQNERIYE-TNY--KYTK
3_Gossypium       501 LSLEAQAEARLLMFSHMNLLSPAIGDPISVPTQDMLIGLYVLTSGNRRGICANRYNPWNRKSYQNERIDDN---NYKSTR
5_Ricinus         475 LSLEAQAEARLLMFSHMNLLSPAIGDPISVPTQDMLIGLYVLTSRNRRGICANRYNPCNHRNYQNERIYDNNNQ---YTK
6_Rosa            508 LSLEAQAEARLLMFSHMNLLSPAIGDPISVPTQDMLIGLYVLTSGNRRGICANRYNPYNRTNSKNERIA-DNNYKY--TK
9_Cucumis         508 LSLEAQAEARLLMFSHMNLLSSAIGDPISVPTQDMLIGLYVLTSGNRRGICANRYNPSNRKNHKNAKIY-NNNYKY--TK
11_Nicotiana      508 LSLEAQVEARLLMFSHMNLLSPAIGDPISVPTQDMLIGLYVLTSGNHRGICVNRYNPCNRRNYQNQKRSDNSHYKY--TK
13_Syringa        508 LSLEAQAEARLLMFSHMNLLSPAIGDPISIPTQDMLIGLYVLTSGNRRGICVNRYNPWNRRNYQNQR-SNNNNYKY--TK
18_Liquidambar    501 LSLEAQAEARLLMFSHMNLLSPAIGDPISVPTQDMLIGLYVLTSGNRRGICANRYNPCNRRNYQNERSD-DNNYKYTKEK
19_Papaver        501 LSLEAQAEARLLMFSHMNLLSPAIGDPISIPTQDMLMGLYVLTIGNRRGICANRYNPCNHLNYQNEKIDDN-NYKYTKEK
20_Ananas         505 LSLEAQAEARLLMFSHMNLLSPAIGDPISVPTQDILIGLYVLTIGNRRGICANMYNPYNCRNYQNQTV-DNNNYKYTKEK
28_Liriodendron   502 LSLEAQAEARLLMFSHMNLLSPAIGDPISVPTQDMLIGLYILTIGNRRGICSNRYNPCNRRNYQNETVDYN---KYTKEK
30_Magnolia       502 LSLEAQAEARLLMFSHMNLLSPAIGDPISVPTQDMLIGLYILTIGNRRGICSNRYNPCNRRNYQNETVDDN---KYTKEK
32_Nymphaea       508 LSLEAQAEARLLMFSHTNLLSPAIGDPISVPTQDMLMGLYVLTMGNRRGICANRYNPCNPRNHQNERI-DHSNYEYRKGK
33_Amborella      501 LSLEAQAEARLLMFSHMNLLSPAIGDPISVPTQDMLIGLYVLTIGNRRGICTNRYNPCNYRNYQNEIV-DDNNYKYTKEK
35_Picea          516 LSLEARAEARLLMFSETNLLSPAIGDPISIPTQDMLLGLYISTVGNSQGIYGNRYHPYHSE-K-----------KSFSCK
44_Ginkgo         508 LSLEAQAEARLLMFSHTNLLYPAIGDPISVPTQDMLLGLYILTVENNQGIYGNRYHPYNSN-K-----------KIFYCK
51_Physcomitrium  498 LSLEAQAEARLLMFSHTNLLSPATGDPVSVPSQDMLLGLYILTIKNHQGIYGNKNHPYKQNNN-----------KIFLNK

T. thermophilus   803 GLEFATPEEALAAHERGEVALNAPIKVAGRETSVGRLKYVFANPDEALLAVAH-------GI---VDLQDVVTVRYMGKR
E. coli           524 GMVLTGPKEAERLYRSGLASLHARVKVRITEYEK------DAN--GELVAKTS--------------------------L
0_Nostoc          532 GNYFSSLEDVIMAFQQDQIDLHAYIYVRFDGEIE------SDQPDTEPVKVTE--------------------------N
1_Litchi          578 NPFFCNSYDAIGAFRQKRINLDSPLWLRWR-LDQRVIVS--R---EAPIEVHYESLGTYHEIFGHYLIVRRVKKEILCIY
2_Arabidopsis     578 EPFFCNSYDAIGAYRQKKINLDSPLWLRWQ-LDQRVIAS--R---EVPIEVHYESFGNYHEIYAHYLIVRSVKKENFCIY
3_Gossypium       578 EPFFCNSYDAIGAYRQKRINLDSPLWLRWQ-LEQCVIAS--R---EAPIEVHYQSSGTYHEIYGHYLIVRSLKKKILCIY
5_Ricinus         552 ESFFSNSYDAIGAYRQKRINLDSPLWLRWQ-LDQRAIAS--R---EAPVEVHYESLGTYHEIYEHYLIVRNIKKEILCIY
6_Rosa            585 EPFFCNSYDAIGAYRQKRVNLDSPLWLRWR-LDQRVITS--R---ETPIEVHYESLGTSHEIYGHYVIVRSIKKEVLCIY
9_Cucumis         585 EPFFCNSYDAIGAYRQKRINLDSPLWLRWR-LDQRVIAS--R---EAPIEVHYESLGTHHEIYGYYLIVKSIKKEILCIY
11_Nicotiana      586 EPFFSNSYDAIGAYRQKRINLDSPLWLRWR-LDQRVIAS--R---ETPIEVHYESLGTFYEIYGHYLIVRSLKKQILFIY
13_Syringa        585 EPFFSNSYNAIGAYRQKRINLDSPLWLRWR-LDQRVIAS--R---ESPIEVHYESLGTYYEIYGHYLIVRSIKKELLFIY
18_Liquidambar    580 EPFFCNSYDAIGAYWQKRINLDSPLWLRWR-LDQRVIAL--R---EAPIEVHYESLGTYHEIYGHYLIVRSVKKETLCIY
19_Papaver        580 EPYFCSSYDALGAYRQKRINLDSPLWLRWR-LDQRVIGS--R---EVPIEVQYDSFGTYHEIYGHYLIVRSVKKETLCIY
20_Ananas         584 EPYFCSSYDALGAYRRKRISLDSPLWLRWR-LDQRVIGS--R---EVPIEVQYESLGTYHEIYGHYLIVGSVKKEIRCIY
28_Liriodendron   579 EPYFCSSYDALGAYRQKRINLDSPLWLRWR-LDQRVIAS--R---EVPIEVQYESLGTYHEIYGHYLIVRSVKKEILCIY
30_Magnolia       579 EPYFCSSYDALGAYRQKRINLDSPLWLRWR-LDQRVIAS--R---EVPIEVQYESLGTYHEIYGHYLIVRSVKKEILCIY
32_Nymphaea       587 EPYFCSSYDALGAYRQKGIDLYSTLWLRWR-LDQRVIASINR---EVPIEVQYESLGTYHEIYDHYRVVRSVKKGMLCIY
33_Amborella      580 EPYFCSSYDALGAYRQKRIDLYSPLWLRWR-LDQCVIASINR---EVPIEVQYESLGIYHEIHEHYRIVKSVKKEIVCIY
35_Picea          584 KPSFYSYDDVLRAYRQKRIDLYSPLWLRWGEVDLRIITSVNQ---EAPIEVQYESLGTFHEIHEHYRIRKGRMGEILNIY
44_Ginkgo         576 KLSFSSYDDALRAYRQKRIHLYSPLWLRWR-VDLRITTSVNR---EAPIEVQYESLGTFREIHEHYRIIRSMMGEILSIY
51_Physcomitrium  567 TPYFSSYDDVIKAYNQKKVRLHSALWLWWG-SKLRTITSINR---EKPIEVQYNSSGISFKIYEHYQLKKNKNEKNFSVY
```

```
T. thermophilus  873 LETSPGRILFARIVAEAVEDEKVAWELIQL-------DVPQEKNSLKDLVYQAFLRLGMEKTARLLDALKYYGFTFSTTS
E. coli          570 KDTTVGR--------------AILWMIVPKGLPYSIVNQALGKKAISKMLNTCYRILGLKPTVIFADQIMYTGFAYAARS
0_Nostoc         580 --EDGTR--------------TLLYKFR-------RVRQDAKGNVLSQYI---YT---TPGRVIYNNAIQ-EALAS----
1_Litchi         652 IRTTVGHISLYREIEEAIQGFCRACSYGT---------------------------------------------------
2_Arabidopsis    652 IRTTVGHISFYREIEEAIQGFSQACSYDT---------------------------------------------------
3_Gossypium      652 IRTTVGHISLYREIEEAIQGFFRAYSYDTQSYGI----------------------------------------------
5_Ricinus        626 IRTTVGHISLYREIEEAIQGFCQAGSDGI---------------------------------------------------
6_Rosa           659 VRTTVGHISLYREIEEAIQGFCRAYSYGT---------------------------------------------------
9_Cucumis        659 IRTTVGHISLYREIEEAIQGFCRACSYGT---------------------------------------------------
11_Nicotiana     660 IRTTVGHIALYREIEEAIQGFSRAYSSGT---------------------------------------------------
13_Syringa       659 IRTTVGHISLYREIEEAVQGFSQACSYGTELS------------------------------------------------
18_Liquidambar   654 IRTTVGHISLYREIEEALQGFYRACSYRT---------------------------------------------------
19_Papaver       654 IRTTVGHISFYREIEEAIQGFSQACSYDT---------------------------------------------------
20_Ananas        658 IRTTLGHISFYREIEEAIQGFCRAYSYTI---------------------------------------------------
28_Liriodendron  653 IRTTVGHISFYREIEEAIQGFCRAYLYDT---------------------------------------------------
30_Magnolia      653 IRTTVGHISFYREIEEAIQGFCRAYSYDT---------------------------------------------------
32_Nymphaea      663 IRTTVGHISFYREIEEAVQGFCRSYSYGT---------------------------------------------------
33_Amborella     656 IRTTVGHISFYREIEEAIQGFCRTY-------------------------------------------------------
35_Picea         661 IRTTVGRTRFNREMEEAIQGFAR-SEHPKKSLPALRI-------------------------------------------
44_Ginkgo        652 IRTTVGRIRFNREIEEAIQGFSRASEHPNKSLKAIRI-------------------------------------------
51_Physcomitrium 643 ICTTVGRIIFNQQIEEAIQGTLKASLFRNQSLPAITI-------------------------------------------

T. thermophilus  946 GITIGIDDAVIPEEKKQYLEEADRKLLQIEQAYEMGFLTD
E. coli          636 GASVGIDDMVIPEKKHEIISEAEAEVAEIQEQFQSGLVTA
0_Nostoc             ----------------------------------------
1_Litchi             ----------------------------------------
2_Arabidopsis        ----------------------------------------
3_Gossypium          ----------------------------------------
5_Ricinus            ----------------------------------------
6_Rosa               ----------------------------------------
9_Cucumis            ----------------------------------------
11_Nicotiana         ----------------------------------------
13_Syringa           ----------------------------------------
18_Liquidambar       ----------------------------------------
19_Papaver           ----------------------------------------
20_Ananas            ----------------------------------------
28_Liriodendron      ----------------------------------------
30_Magnolia          ----------------------------------------
32_Nymphaea          ----------------------------------------
33_Amborella         ----------------------------------------
35_Picea             ----------------------------------------
44_Ginkgo            ----------------------------------------
51_Physcomitrium     ----------------------------------------
```

**Fig. S6.** Sequence alignment of the β" subunits from PEP of angiosperms with those of the RNAPs from *E. coli*, *T. thermophilus* and Nostoc. The residues conserved more than 50 % are in red, those mutated in similar residues are in blue. The strictly conserved residues described by Lane & Darst (Lane & Darst, 2010) are highlighted in gray. The blue triangles show mutations observed among the strictly conserved residues described (Lane & Darst, 2010). The non-conservative mutations, at least three in a row in the β or β' domain in *E. coli* and *T. thermophilus*, are highlighted in green and displayed on the *E. coli* structure (PDB entry: 6GH5). Those colored in orange are nearby to the DNA, those in green are located at the surface of the subunits. The domains described for all-RNA polymerase (a) and the bRNAPs (b) are also given and highlighted in yellow and cyan respectively. The name of the RNAP domains are also given and highlighted in purple and green (Lane & Darst, 2010; Sutherland & Murakami, 2018).

```
                                                        β'a13:L914-E979
                                                   β'b6:K908-F1011
T. thermophilus  888 EAVEDEKVAWELIQLDV-------P-------QEKNSLKDLVYQAFLRLGMEKTARLLDALKYYGFTFSTTSGITIGIDD
E. coli          571 DTTVGRAILWMIVPKGL-------PYSIVNQALGKKAISKMLNTCYRILGLKPTVIFADQIMYTGFAYAARSGASVGIDD
0_Nostoc           1 ---------------------MTEKMIFRNRVVDKGQLRNLISWAFTHYGTARTAVMADKLKDLGFRYATRAGVSISVDD
1_Litchi           1 --------------MAER-----AGLVFHNKMIDGTAIKRLISRLIDHFGMAYTSHILDQVKTLGFQQATATSISLGIDD
2_Arabidopsis      1 --------------MAER-----ANLVFHNKVIDGTAIKRLISRLIDHFGMAYTSHILDQVKTLGFQQATATSISLGIDD
3_Gossypium        1 --------------MAER-----ANLVFHNKVIDGTAIKRLISRLIDHFGMAYTSHILDQVKALGFQQATATSISLGIDD
5_Ricinus          1 ----------MEVLMAKR-----ANLVFHNKVIDGTAIKRLISRLIDHFGMAYTSHILDQVKTLGFQQATATSISLGIDD
6_Rosa             1 --------------MAER-----ASLVFHNKVIDGTAIKRLISRLIDHFGMAYTSHILDQVKTLGFRQATATSISLGIDD
9_Cucumis          1 --------------MAER-----ADLVFHNKVIDGTAIKRLISRLIDHFGMAYTSHILDQLKTLGFQQATATSISLGIDD
11_Nicotiana       1 --------------MAER-----ANLVFHNKAINGTAMKRLISRLIDHFGMAYTSHILDQVKTLGFQQATATSISLGIDD
13_Syringa         1 ----------MEVLMAER-----TNLVFHNKVIDGTAMKRLISRLIDHFGMAYTSHILDQVKTLGFHQATATSISLGIDD
18_Liquidambar     1 ----------MEVLMAER-----ANLVFHNKVIDGTAMKRLISRLIDHFGMAYTSHILDQVKTLGFQQATATSISLGIDD
19_Papaver         1 --------------MAER-----ADLVFHNKAIDGTAIKRLISRLIDHFGMAYTSHILDQVKTLGFHQATATSISLGIDD
20_Ananas          1 --------------MAER-----ADLVFHNKVINGTAMKRLISRLIDHFGMGYTSHILDQVKTLGFHQATATSISLGIDD
28_Liriodendron    1 ----------MEVLMAER-----ADLVFHNKVIDATAMKRLISRLIDHFGMAYTSHILDQVKTLGFQQATATSISLGIDD
30_Magnolia        1 ----------MEVLMAER-----ADLVFHNKVIDATAMKRLISRLIDHFGMAYTSHILDQVKTLGFQQATATSISLGIDD
32_Nymphaea        1 ----------MEVLMAER-----ADLVFHNKVIDGTAMKRLISRLIDHFGIAYTSHILDQVKTLGFQQATATSISLGIDD
33_Amborella       1 --------------MAER-----AGLVFHNKVIDGTAIKRLISRLIDHFGMAYTSHILDQVKTLGFRQATATSISLGIDD
35_Picea           1 ------MKIWRFFLMKERTRLPFDNLPFYNKVMDKTAIKKLISRLIDHFGMTYTSHILDQLKTSGFQQATDTAISLGIDD
44_Ginkgo          1 --------------MTER-----AKLLFHNKVMNRIATKQLISRLIDHFGMTYTSHISDQLKASGFQQATDAAISLGIDD
51_Physcomitrium   1 ------------------------MLFYNKVMDRTAIKQLISRLITHFGITYTTYILDQLKTVGFKQATQAAISLGIDD

                                   Secondary channel rim-helices
             β'a13:L984-E979          β'a14:T984-F1011
                          β'b6:K908-F1011
T. thermophilus  954 AVIPEEKKQYLEEADRKLLQIEQAYEMGFLTDRERYDQILQLWTETTEKVTQAVFKNFE------------ENYPFNPLY
E. coli          644 MVIPEKKHEIISEAEAEVAEIQEQFQSGLVTAGERYNKVIDIWAAANDRVSKAMMDNLQTETVINRDGQEEKQVSFNSIY
0_Nostoc          60 LMVPPTKRSLLEAAEEEIRATEARYQRGEITEVERFQKVIDTWNGTSEALKDEVVVHFK------------KTNPLNSVY
1_Litchi          62 LLTIPSKGWLVQDAEQQSLILEKHHHYGNVHAVEKLRQSIEIWYATSEYLRQEMNPNFR------------MTDPLNPVH
2_Arabidopsis     62 LLTIPSKGWLVQDAEQQSWILEKHHHYGNVHAVEKLRQSIEIWYATSEYLRQEMNPNFR------------MTDPFNPVH
3_Gossypium       62 LLTIPSKGWLVQDAEQQSLILEKHHHFGNVHAVEKLRQSIEIWYATSEYLRQEMNPNFR------------MTDPFNPVH
5_Ricinus         66 LLTIPSKGWLVQDAEQQSLILEKHYHYGNVHAVEKLRQSIEIWYATSEYLRQEMNLNFR------------MTEPFNPVH
6_Rosa            62 LLTIPSKGWLVQDAEQQSLILEKHHHYGNVHAVEKLRQSIEIWYATSEYLRQEMNPNFR------------MTDPFNPVH
9_Cucumis         62 LLTIPSKGWLVQDAEQQSLILEKHHHYGNVHAVEKLRQSIEIWYATSEYLRQEMNPNFR------------MTDPFNPVH
11_Nicotiana      62 LLTIPSKGWLVQDAEQQSLILEKHHHYGNVHAVEKLRQSIEIWYATSEYLRQEMNPNFR------------MTDPFNPVH
13_Syringa        66 LLTIPSKRWLVQDAEQQSLILEKHHHYGNVHAVEKLRQSIEIWYATSEYLRQEMNPNFR------------MTDPFNPVH
18_Liquidambar    66 LLTIPSKGWLVQDAEQQSFILEKHHHYGNVHAVEKLRQSIEIWYATSEYLRQEMNPNFR------------MTDPFNPVH
19_Papaver        62 LLTIPSKGWLVQDAEQQSLILEKHHHYGNVHAVEKLRQSIEIWYATSEYLRQEMNPNFR------------MTDPLNPVH
20_Ananas         62 LLTIPSKGWLVQDAEQQSFILEKHHHYGNVHAVEKLRQSIEIWYATSEYLRQEMNPNFR------------MTDPSNPVY
28_Liriodendron   66 LLTIPSKGWLVQDAEQQSLILEKHHHYGNVHAVEKLRQSIEIWYATSEYLRQEMHPNFR------------MTDPSNPVH
30_Magnolia       66 LLTIPSKGWLVQDAEQQSLILEKHHHYGNVHAVEKLRQSIEIWYATSEYLRQEMHPNFR------------MTDPSNPVH
32_Nymphaea       66 LLTTPSKRWLVQDAEQQSLILEKHHHYGNVHAVEKLRQSIEIWYATSEYLRQEMNLNFK------------MTDPSNPVH
33_Amborella      62 LLTIPSKGWLVQDAEQQSLILEKHHHYGNVHAVEKLRQSIEIWYATSEYLRQEMNPNFR------------MTDPFNPVH
35_Picea          75 LLTAPSKAWLVQDAEQQGSVSEKQNHYGNVHAVEKLRQSIEIWYATSEYLRKEMNPNFS------------MTDPLNPVH
44_Ginkgo         62 LLTAPSRGWLVRDAEQQGSISEKHHHYGNVYAVEKLRQSIETWYATSEYLRQEMNPNFG------------MTDPSNPVH
51_Physcomitrium  56 LLTAPSKSWLIQDAEQQGYISEKHYRYGNVHAVEKLRQLIETWYATSEYLKQEMNPNFR------------MTDPLNPVH

                                                                Bridge helix
                                       β'a15:N1018-G1113
                                       β'b7:N1018-G1113
T. thermophilus 1022 VMAQSGARGNPQQIRQLCGLRGLMQKPSGETFEVPVRSSFREGLTVLEYFISSHGARKGGADTALRTADSGYLTRKLVDV
E. coli          724 MMADSGARGSAAQIRQLAGMRGLMAKPDGSIIETPITANFREGLNVLQYFISTHGARKGLADTALKTANSGYLTRRLVDV
0_Nostoc         128 MMAFSGARGNISQVRQLVGMRGLMADPQGEIIDLPIKTNFREGLTVTEYIISSYGARKGLVDTALRTADSGYLTRRLVDV
1_Litchi         130 IMSFSGARGNASQVHQLVGMRGLMSDPQGQMIDLPIQSNLREGLSLTEYIISCYGARKGVVDTAVRTSDAGYLTRRLVEV
2_Arabidopsis    130 MMSFSGARGNASQVHQLVGMRGLMSDPQGQMIDLPIQSNLREGLSLTEYIISCYGARKGVVDTAVRTSDAGYLTRRLVEV
3_Gossypium      130 IMSFSGARGNASQVHQLVGMRGLMSDPQGQMIDLPIQSNLREGLSLTEYIISCYGARKGVVDTAVRTSDAGYLTRRLVEV
5_Ricinus        134 IMSFSGARGNVSQVHQLVGMRGLMSDPQGQMIDLPIQSNLREGLSLTEYIISCYGARKGVVDTAVRTSDAGYLTRRLVEV
6_Rosa           130 MMSFSGARGNASQVHQLVGMRGLMSDPQGQMIDLPIQSNLREGLSLTEYIISCYGARKGVVDTAVRTSDAGYLTRRLVEV
9_Cucumis        130 IMSFSGARGNASQVHQLVGMRGLMSDPQGQMIDLPIQSNLREGLSLTEYIISCYGARKGVVDTAVRTSDAGYLTRRLVEV
11_Nicotiana     130 IMSFSGARGNASQVHQLVGMRGLMSDPQGQMIDLPIQSNLREGLSLTEYIISCYGARKGVVDTAVRTSDAGYLTRRLVEV
13_Syringa       134 IMSFSGARGNASQVHQLVGMRGLMSDPQGQMIDLPIQSNLREGLSLTEYIISCYGARKGVVDTAVRTSDAGYLTRRLVEV
18_Liquidambar   134 IMSFSGARGNASQVHQLVGMRGLMSDPQGQMIDLPIQSNLREGLSLTEYIISCYGARKGVVDTAVRTSDAGYLTRRLVEV
19_Papaver       130 IMSFSGARGNASQVHQLVGMRGLMSDPQGQMIDLPIQSNLREGLSLTEYIISCYGARKGVVDTAVRTSDAGYLTRRLVEV
20_Ananas        130 LMSFSGARGNASQIHQLVGMRGLMSDPQGQMIDLPIQSNLREGLSLTEYIISCYGARKGVVDTAVRTSDAGYLTRRLVEV
28_Liriodendron  134 IMSFSGARGNASQVHQLVGMRGLMSDPQGQMIDLPIQSNLREGLSLTEYIISCYGARKGVVDTAVRTSDAGYLTRRLVEV
30_Magnolia      134 IMSFSGARGNASQVHQLVGMRGLMSDPQGQMIDLPIQSNLREGLSLTEYIISCYGARKGVVDTAVRTSDAGYLTRRLVEV
32_Nymphaea      134 IMSYSGARGNASQVHQLVGMRGLMSDPQGQMIDLPIQSNLREGLSLTEYTISCYGARKGVVDTAVRTSDAGYLTRRLVEV
33_Amborella     130 MMSFSGARGNASQVHQLVGMRGLMSDPQGQMIDLPIQSNLREGLSLTEYIISCYGARKGVVDTAVRTSDAGYLTRRLVEV
35_Picea         143 VMSFSGARGSTSQVHQLVGMRGLMSDPQGQIIDLPIRRNLREGLSLTEYIISCYGARKGVVDTAVRTADAGYLTRRLVEV
44_Ginkgo        130 MMSFSGARGNTSQVHQLVGMRGLVSDPQGQIIDLPIQRNFREGLSLTEYIISCYGARKGVVDTAVRTSDAGYLTRRLVEV
51_Physcomitrium 124 MMSFSGARGSTSQVHQLVGMRGLMSDPQGQIIDLPIQSNFREGLSLTEYIISCYGARKGVVDTAVRTSDAGYLTRRLVEV
                                                                      ▲  ▲ ▲
```

```
T. thermophilus 1102 THEIVVREADCGTTNYISV-PLFQPDEVTRSLRLRKRADIEAGLYGRVLAREVEVLGVR---LEEGRYLSMDDVHLLIKA
E. coli          804 AQDLVVTEDDCGTHEGIMMTPVIEGGDVKEPLRDR--------VLGRVTAEDVLKPGTADILVPRNTLLHE----QWCDL
0_Nostoc         208 SQDVIIREFDCGTTRGIPIRPMTEGAK---TLIPL-----ANRLMGRVIGEDVVHPVTKEVIAPRNTPISDDLAKEI---
1_Litchi         210 VQHIVVRRTDCGTIRGISVSPQN--RMMSERVF-------SQTLMGRVLADDIYI--GPRCLAIRNQDIGIGLVNRF---
2_Arabidopsis    210 VQHIVVRRTDCGTIRGISVSPRNKNRMMSERIF-------IQTLIGRVLADDIYI--GSRCVAFRNQDLGIGLVNRL---
3_Gossypium      210 VQHIVVRRTDCGTTRGISVSPQK--RTLPERIF-------IQTLIGRVLADDIYM--GPRCIAIRNQDIGLGLVDRF---
5_Ricinus        214 VQHIVVRRTDCGTTRGISVSPQN--GMMSERIF-------IQTLIGRVLADNIYM--GLRCIAIRNQDIGIRLANRF---
6_Rosa           210 VQHIVVRRTDCGTVRGISVSPRN--GMMPERIF-------IQTLIGRVLADDIYI--GPRCIAVRNQDIGIGLVNRF---
9_Cucumis        210 VQHIVVRRTDCGTIRGILVSPGN--RMIPERIF-------IQTLIGRVLADDIYM--GPRCIGVRNQDIGIGLINRF---
11_Nicotiana     210 VQHIVVRRTDCGTARGISVSPRN--GMMPERIF-------IQTLIGRVLADDIYM--GPRCIATRNQDIGIGLVNRF---
13_Syringa       214 VQHIVVRRTDCGTSRGISVSPRN--GMMPERIF-------IQTLMGRVLADDIYT--GTRCIASRNQDVGIGLVNRF---
18_Liquidambar   214 VQHIVVRRTDCGTTRGISVSSRN--GMMPERIF-------IQTLIGRVLADDIYM--GPRCIAIRNQDIGIGLVNRF---
19_Papaver       210 VQHIVVRRTDCGTIRGISVSPRN--GMMTERIF-------IQTLIGRVLADDIYM--GSRCIAIRNQDIGIGLVNRF---
20_Ananas        210 VQHIIVRRRDCGTIRGISVSPQN--GM-TEKIF-------VQTLIGRVLADDIYI--GLRCIATRNQDIGIGLVNRF---
28_Liriodendron  214 VQHIVVRRTDCGTIRGISVSPRN--GM-TEKIL-------IQTLIGRVLADDIYM--GLRCIATRNQDIGIGLVNRF---
30_Magnolia      214 VQHIVVRRTDCGTIRGISVSPRN--GM-TEKIW-------IQTLIGRVLADDIYM--GLRCIATRNQDIGIGLVNRF---
32_Nymphaea      214 VQHIVVRRTDCGSTRGISVSLRK--GM-TERIF-------IQTLIGRVLANDVYL--GLRCIATRNQDIGIGLVNRF---
33_Amborella     210 VQHIVVRRTDCGNIRGISVSSRN--GMMSERIF-------IQTLIGRVLADDIYI--GPRCIAVRNQDIGIGLVNRF---
35_Picea         223 VQHIVVRRKDCGTIQGIFVSPIRGRERDRNEIVVR-----TQILIGRVLADDVYI--NRRCIATRNQDIGVGLANQL---
44_Ginkgo        210 VQHIVVRRADCGTIRGISVSPIRGRERIKKEFVL-------QTLIGRVLADDVHI--NKRCIATRNQDIGVGLADQL---
51_Physcomitrium 204 VQHIVVRKVDCGTSENIFVTPLQNNY-------KK-----NNKLIGRILADNIYI--NGRCIAIRNQDITTNLVISL---

                                                                      Trigger loop-helix1 Trigger loop
                                                                    β'a16:R1213-A1247
                                                      β'b8:V1186-A1247
T. thermophilus 1178 AEAGEIQEVPVRSPLTCQTRYGVCQKCYGYDLSMARPVSIGEAVGIVAAQSIGEPGTQLTMRTFHTGGVAGAA-------
E. coli          872 LEENSVDAVKVRSVVSCDTDFGVCAHCYGRDLARGHIINKGEAIGVIAAQSIGEPGTQLTMRTFHIGGAASRAAAESSIQ
0_Nostoc         277 -GRSGVGEVVVRSPLTCEAARSVCQHCYGWSLAHAKMVDLGEAVGIIAAQSIGEPGTQLTMRTFHTGGVFTGEVAQQVRS
1_Litchi         276 -ITFRTQAISIRTPFTCRSTSWICRLCYGRSPTHGDLVELGEAVGIISGQSIGEPGTQLTLRTFHTGGVFTGGTAEHVRA
2_Arabidopsis    278 -ITFGTQSISIRTPFTCRSTSWICRLCYGRSPTHGDLVELGEAVGIIAGQSIGEPGTQLTLRTFHTGGVFTGGTAEHVRA
3_Gossypium      276 -RAFRTQPISIRTPFTCRSTSWICRLCYGRSPTHGDLVELGEAVGIIAGQSIGEPGTQLTLRTFHTGGVFTGGTAEHVRA
5_Ricinus        280 -ITFRTQTISIRTPFTCRSTSWICRLCYGRSPTHGDLVELGEAVGIIAGQSIGEPGTQLTLRTFHTGGVFTGGTAEHVRA
6_Rosa           276 -ITFQTQPIPIRTPFTCKSTSWICRLCYGRSPTHGDLVELGEAVGIIAGQSIGEPGTQLTLRTFHTGGVFTGGTAEHVRA
9_Cucumis        276 -ITFQTQPISIRTPFTCRSTSWICRLCYGRSPTHGDLVELGEAVGIIAGQSIGEPGTQLTLRTFHTGGVFTGGTAEHIRA
11_Nicotiana     276 -ITFRAQPISIRTPFTCRSTSWICRLCYGRSPTHGDLVELGEAVGIIAGQSIGEPGTQLTLRTFHTGGVFTGGTAEHVRA
13_Syringa       280 -ITFRAQPISIRTPFTCRSASWICRLCYGRSPTHGDLVELGEAVGIIAGQSIGEPGTQLTLRTFHTGGVFTGGTAEHVRA
18_Liquidambar   280 -ITFRAQPISIRTPFTCRSTSWICRLCYGRSPTHGDLVELGEAVGIIAGQSIGEPGTQLTLRTFHTGGVFTGGTAEHVRA
19_Papaver       276 -ITFRAQPIYIRTPFTCRSTSWICRLCYGRSSTHGDLVELGEAVGIIAGQSIGEPGTQLTLRTFHTGGVFTGGTAEHVRA
20_Ananas        275 -ITFRAQPIYIRTPFTCRSTSWICQLCYGRSPTHGDLVELGEAVGIIAGQSIGEPGTQLTLRTFHTGGVFTGGTAEHVRA
28_Liriodendron  279 -ITFRAQSIYIRTPFICRSTSWICRLCYGRSPTHGDLVELGEAVGIIAGQSIGEPGTQLTLRTFHTGGVFTGGTAEHVRA
30_Magnolia      279 -ITFRAQSIYIRTPFICRSTSWICRLCYGRSPTHGDLVELGEAVGIIAGQSIGEPGTQLTLRTFHTGGVFTGGTAEHVRA
32_Nymphaea      279 -MTSRAQPIYIRTPFTCRSASWICRLCYGRSPTHGDLVELGEAVGIIAGQSIGEPGTQLTLRTFHTGGVFTGGTAEHVRA
33_Amborella     276 -ITFQTQPISIRTPFTCKSTSWICRLCYGRSPTHGDLVELGEAVGIIAGQSIGEPGTQLTLRTFHTGGVFTGGTAEHVRA
35_Picea         293 -INLRTQPIYIRTPFTCKSISRICQLCYGRSTTHNHLIELGEAVGIIAGQSIGEPGTQLTLRTFHTGGVFTGDIAEHVRA
44_Ginkgo        278 -RTLRTQPIYIRTPPTCKSLSRICRLCYGRSPTHSNLIELGEAVGIIAGQSIGEPGTQLTLRTFHTGGVFTGDIAEHVRA
51_Physcomitrium 267 -INFQRKGIFIRSPLICKSMLWICQLCYGWSLTHGNLIELGEAVGIIAGQSIGEPGTQLTLRTFHTGGVFTGDIAEHIRT
                                                                                  ▲  ▲

T. thermophilus 1251 --------------------------------------------------------------------------------
E. coli          952 VKNKGSIKL-----------------------------------------------------------------------
0_Nostoc         356 -KIDGTVKIPRKLRTRQYRTRHGEDALYVEANGVIILEPKKEGDATPANQEVQLTQGSTLYVFDGNQVKQGQLLAEVALG
1_Litchi         355 -PSNGKIKFNEDLV-HPTRTRHGHPAFLCYIDLYVIIESEDI------MHKVTIPPKSFLLVQNDQYVESEQVIAEIRAG
2_Arabidopsis    357 -PYNGKIKFNEDLV-HPTRTRHGHPAFLCYIDLSVIIESEDI------IHSVTIPPKSFLLVQNDQYVESEQVIAEIREG
3_Gossypium      355 -PFNGKIKFNEDLV-HPTRTRHGHPAFLCYRDLYVIIESEDI------IHKVTIPPKSFLLVQNDQYVESEQVIAEIRAG
5_Ricinus        359 -PSNGKIKFNEDLV-HPIRTRHGHPAFLCYIDLYVTIESHDI------IHNATIPPKSFLLVQNNQYVESEQVIAEIRAG
6_Rosa           355 -PSNGKIKFNEDLV-HPTRTRHGHPAFLCYIDLYVTIESENI------IHNVTIPPKSLILVQNDQYVESEQVVAEIRAG
9_Cucumis        355 -SSNGKIKFNENLV-HPTRTRHGHPAFLCYIDLYVTIESEDI------IHNVTIPPKSLLLVQNDQYVESEQVIAEIRAG
11_Nicotiana     355 -PSNGKIKFNEDLV-HPTRTRHGHPAFLCSIDLYVTIESEDI------LHNVNIPPKSLLLVQNDQYVESEQVIAEIRAG
13_Syringa       359 -PSNGKIKFNEDLV-HPTRTRHGHPAFLCSIDLYVTIESEDI------LHNVNIPSKSFLLVQNDQYVESEQVIAEIRAG
18_Liquidambar   359 -PSNGKIKFNEDLV-HPIRTRHGHPAFLCYIDLYVTVESEDI------LHNVNIPQKSFLLVQNDQYVESEQVIAEIRAG
19_Papaver       355 -PSNGKIKFNEDLV-HPTRTRHGHPAFLCYIDLFVTIESQDI------IHNVNIPPKSFLLVQNDQYVESEQVIAEIRAG
20_Ananas        354 -PSNGKIKFNEDLV-HPTRTRHGHPAFLCSIDLYVTIESRDI------IHNVTIPPKSLILVQNDQYVESEQVIAEIRAG
28_Liriodendron  358 -PSNGKIKFNECLV-HPTRTRHGHPAFLCYIDLYVTIESQDI------IHNVNIPPKSFLLVQNDQYVESEQVIAEIRAG
30_Magnolia      358 -PSNGKIKFNEDLV-HPTRTRHGHPAFLCYIDLYVTIESQDI------LHNVNIPPKSFLLVQNDQYVESEQVIAEIRAG
32_Nymphaea      358 -PSNGKIKFNEDLV-HPTRTRHGHPAFLCYVDLYVTIESQDI------IHSVNIPPKSFLLVQNDQYVESEQVIAEIRAG
33_Amborella     355 -PSNGKIKFNEDLA-HPTRTRHGHPAFLCSIDLDVIIESEDI------IHNVTIPPKSLILVQNDQYVESEQVIAEIRAR
35_Picea         372 -PFNGKIEFNDNLV-YPTRTCNGHPAYLCHNNLSITIHGQDQ------VKNLTIPPQSLLLVQNDQYVESEQIIAEVRAR
44_Ginkgo        357 -PFNGKIQFNENLV-HPTRTRHGHPASICHNELSITIDGQDQ------VHSLTIPSQSLLLVQNDQYVESEQVIAEARAR
51_Physcomitrium 346 -PFNGIIQFDTNSV-YPTRTRHGHPAWICNNNLSVVIKSKKK------LHNLVIPTQSLLLVQSNQYVESKQVIAEVRAK
```

```
T. thermophilus 1251 --------------------------------------------------------------------------------
E. coli          961 -------------------------------------SNVKSVVNSSGKLVITS--------------------------
0_Nostoc         435 GRTTRTNTEKAVKDVASDLAGEVQFAEVVPEQKTDRQGNTTTTAARGGLIWILSGEVYNLPPGAELVVKNGDAIASNGVL
1_Litchi         427 TY-TLNFKERVRKHIYSDSEGEMHWSTDVYHAPEFTYSNVH-LLPKTSHLWILSGGSCGSGVVSFSLYKDQDQLSIHYRS
2_Arabidopsis    429 TY-TFHFKERVRKYIYSDSEGEMHWSTDVSHAPEFTYSNVH-LLPKTSHLWILSGGSCGSSLIRFSIHKDQDQMNIPFLS
3_Gossypium      427 TY-TLNLKERVRKHIYSDSEGEMHWSTDVYHSPEFTYSNVH-LLPKTSHLWILSGGSYKFSVVPFSLHKDQDQINIHYLS
5_Ricinus        431 TY-TLNFKEKVRKHIYSDSEGEMHWSTDVYHAPEFTYSNVH-LLPKTSHLWILSGNSCRSSIVPFSLHKDQDQMNVHSLS
6_Rosa           427 AY-TFNFKERVRKHIYSDSEGEMHWSTDVYHAPEFTYSNVH-LLPKTSHLWILSGGSCRFSAVPPSLHKDQDQTNVHSLS
9_Cucumis        427 TY-TLNLKERVRKHIYSDSEGEMHWSTDVYHAPEFTYSNVH-LLPKTSHLWILSGGSCGCSVVPFSLYKDQDQINVHSLC
11_Nicotiana     427 IS-TLNFKEKVRKHIYSDSDGEMHWSTDVYHAPEFTYGNVH-LLPKTSHLWILLGRPCRSSLVYLSIHKDQDQMNAHFLS
13_Syringa       431 TS-TLNFKEKVRKHIYSDSDGEMHWSTDVYHAPEFTYGNVH-LLPKTSHLWILLGGPCRSSLVSLSLHKDQDQINAHSRS
18_Liquidambar   431 TY-TFNFKERVRKHIYSDSEGEMHWSTDVYHAPEFTYGNVH-LLPKTSHLWILSGGSCRSSVVPFSLHKDQDQMNVHSLS
19_Papaver       427 TS-TFNFKERVRKHIYSDLEGEMHWSTDVYHAPEYTYGNVH-LLPKTSHLWILSGGLCRSSIVPFSLGKDQDQTNVHSLF
20_Ananas        426 TS-TFHFKERVRKHIYSESEGEMHWSTDVYHAPEYTYGNVH-LLPKTSHLWILAGGPCRSSIVSFSLHKDQDQMNVYSLS
28_Liriodendron  430 TS-TFNFKERVRKHIYSDSEGEMHWSTDVYHAPEYRYGNVH-LLPKTSHLWILSGGPCRSSIVPFSLHKDQDQMNVHSLS
30_Magnolia      430 TS-TFNFKERARKHIYSDSEGEMHWSTDVYHAPEYRYGNVH-LLPKTSHLWILSGGPCRSSIVTFSLHKDQDQMNVHSLS
32_Nymphaea      430 TS-TFHFKERVRKHIYSDSEGEMHWSTGVYHAPEYTHGNVH-FLPKTSHLWILSGGPCKSSLVPFSLHKDQDQMNVQSLS
33_Amborella     427 TS-TFNFKEKVRKHIYSDSEGEMHWSTDVYHAPDFTYSNVH-LLPKTSHLWILSGSSYRSSVVPFSIHKDQDQTNVYFLS
35_Picea         444 TS---SFKEKVRKNIYSDLEGEMHWSTNVCHAPEYVHGNVH-PILRTGYLWILSGGIYGSGVVPFPFHKHQDQVDVQPFV
44_Ginkgo        429 TS---PSKEKVMKHIYSDLEGEMHWSTNVCHAPENVHGNVH-LILRTSYLWVLSGGLYESGVVPFPLYKDQDQVNIQFPL
51_Physcomitrium 418 TS---PFKEKVQKYIYSNLSGEMHWSSKVQHSSEYIHSNVH-LLRKTGHIWILAGNFDKDNKFSFIFYQNQDKLDNKLPI
```

```
T. thermophilus 1251 --------------------------------------------------------------------------------
E. coli          978 ---------------------------------------------------------------RNTELKLIDEFGRTKES
0_Nostoc         515 AETKLTT------LHGGVVRL-------------PEATPGK--------STREIEIITASVVLDQATVTVQSS--QGRNN
1_Litchi         505 VERRYISSLS---VNNDQVRHQLVSSDFSDNK--EDGISDY-SGFNRIIGIGHCNLIHAAILHENSD--LLAK--RRRNR
2_Arabidopsis    507 AERKSISSLS---VNNDQVSQKFFSSDFADPK--KLGIYDY-SELNGNLGTSHYNLIYSAIFHENSD--LLAK--RRRNR
3_Gossypium      505 AERRYISRFS---VNNDQVRHNLFSSDFSDKK--EERIYDY-SELNRIIGTGHCDFIYSAILHENAD--LLAK--RRRNR
5_Ricinus        509 IKRRYISSPSVNSVNNDQVKPKFFSSDFSGKK--PSRIPYY-SELNRIVCTGHCNLIYPAILYENSD--LLAK--RRRNK
6_Rosa           505 VEGRYFSSLS---VNNDQVKHKFFGLNLSGKK--ESCIPDY-SELNRIIYTSHCNLIFPPIRHDN-F--LLTK--RRRNR
9_Cucumis        505 VERRYISSLS---VNNDKVGQKFYGPDLSGKN--ESGIPDY-SELNPILCTGQSNLTYPAIFHGNSD--LLAK--RRRNG
11_Nicotiana     505 GKRRYTSNLS---VTNDQARQKLFSSDFSGKK--EDRIPDY-SDLNRIICAGQYNLVYSPILHENSD--LLSK--RRRNK
13_Syringa       509 VKRRYTSNLS---GTNDPERQKLFSSYFYGKKKXEDRISDY-SDLNRIICNGRCNLIYPTILHQNSD--LFSK--RRRNR
18_Liquidambar   509 VERRYISNLS---VTNDQVRHKLFSSDISGKK--EGRIPDY-SELNRIICSGHCNLIYPAILRENSD--LLAK--RRRNR
19_Papaver       505 AKQRYTPSLS---VTNDQVKQKFCSSESSGTG--GRGVLDY-SGPDRIICNGHCNFIYPPILHESSD--LLAK--RRRNR
20_Ananas        504 VEGRYISNPS---MTNDQVRHKLLDT--SGKK--DRKILDY-SRLDRIISNGHWNFIYPSILQENPD--FLAK--KRRNR
28_Liriodendron  508 VERRYISDLS---VTNDRVRHKLFSSDPSGKK--KERILDY-SGPDRIVSNGHWNFLYPAILHENSD--LLAK--RRRNR
30_Magnolia      508 VERRYISDLS---VTNDRVRHKLFSSDPSSKK--GKGILDY-SGPDRIISNGHWNFLYPATLHENSD--LLAK--RRRNR
32_Nymphaea      508 VQERSISDFS---VNNNRVKHKLFGSDPLARK--GRRISDYAAGLERVISNGDGDFIYPAILRENSY--LLAK--RRRNR
33_Amborella     505 AEGKNISSRSVNTVNNDQGKGKFFSSDFSGKK--ESTIPDY-SEFNRIIDRDHWNLIFPSILHKNYDLFLLAK--RRRNR
35_Picea         520 AKHQSLFDSY---V--DQVEHRSGDSNCYGKE---EQIFSY-SETDRTISNEHRDSIYVTLSPKNYN--MKGK--RQMNR
44_Ginkgo        505 AKHKSLSDSS---VNQDRVKHKSVDSNFSGKE---EKISGY-SGIDRIMSNEHWDSIYSTIPFDNCK--ILGK--KQRNR
51_Physcomitrium 494 AKQ----------------------------------TLNY-FQLKEHFLNNFWNSIYSSIILYNYR--FLEK--K-NNK
```

```
T. thermophilus 1251 --------------------------------------------------------------------------------
E. coli          995 YKVPYG------------------------AV------------------------------------------------
0_Nostoc         566 YLVSTGNNQVFN------------LRATPGTKVQNGQVVAELIDDRYRTTTGGFLKFAGVEVQKKGKA------------
1_Litchi         575 FLIPFQSIQEQEKELMPH--SGISIEIPIKGVFRKNSIFAYFDDPRYRRKNSGITKYGTIGAHSIVKKEDLIEYRGRGGK
2_Arabidopsis    577 FLIPFQSIQEQEKEFIPQ--SGISVEIPINGIFRRNSIFAFFDDPRYRRKSSGILKYGTLKADSIIQKEDMIEYRG--VQ
3_Gossypium      575 FIIPFQLIQDQEKELMLHSHSGISMEIPINGIFRRKSILAFFDDPRYRRKSSGITKYGTLGAHSIVKREDVIEYRG--VK
5_Ricinus        582 FIIPFQSIQEQEKKLMTRS-SAISIEIPLNGIFRRNSVFAYFDDPQYRRKSSGITKYGAIGVHSIVKKEDLIEYRG--VK
6_Rosa           574 FIIPFQSIQEQEKERMPR--PDISIEIPINGIFRRNSILAYFDDPQYRRKSSGITKYGTVGLHSILKKEDLIEYRG--VK
9_Cucumis        575 FIIQFESLQEREKELRPP--SGISIEIPINGLFRRNSILAFFDDPQYRRNSSGITKYGTIGVHSILKKEDLIEYRG--VK
11_Nicotiana     575 FIIPLHSIQELENELMPC--SGISIEIPVNGIFRRNSILAYFDDPRYRRKSSGIIKYGTVETHSVIKKEDLLEYRG--VK
13_Syringa       581 FIIPLQSIQERENELMPR--SGISIEIPPNGIFRRNSILAYFDDPRYRRKSSGITKYGTIEMHSIVKKEDLIEYRG--VK
18_Liquidambar   579 FIIPFQSIQEREKEQMPHSNSGISIEIPINGIFRRNSILAYFDDPRYRRKSSGITKYGTIEVHSIVKKEDLIECRG--VK
19_Papaver       575 LIIPFQSNQERDKERIPR--SGISIEIPINGIFRRNSILAYFDDPRYRRNSSGITKYETLEMHSIVKKEDFIEYRR--AK
20_Ananas        572 FIIPLQYDQEREKELIPC--FGISIEIPINGILRRNSILAYFDDPRYRRSSSGITKYGTIEVDSIIKKEDLIEYRG--AK
28_Liriodendron  578 FIIPFQYDQEREKELMPR--SGISIEIPINGILRRDTILAYFDDPRYRRSSSGITKYGTIEVDSIVKKEDLIEYRG--AK
30_Magnolia      578 FIIPFQYDQEREKELMPR--SGISIEIPINGILRRNSILAYFDDPRYRRSSSGITKYGTIEVDSIVKKEDLIEYRG--AK
32_Nymphaea      579 FIIPFQYDPEREKELTPHSSTSITVEIPANGILRRNSILAYFDDPRYRRSSSGITKYGIIEVDSIVKKEGLIEYRR--PK
33_Amborella     580 FIIPFQWIQERENELMLR--SSISIEIPINGVFRKNSILAYFDDPQYRRKSSGITKYGAIGLHSIFKKEDLIEYVG--IK
35_Picea         587 FIVPLQCDKEWGKRIISF--PDAILRIPKSGVLQRNSIFGY---------------------------------------
44_Ginkgo        574 LIVPLRYDKEREKRRIPC--PNSILRIPRNFLFQRNHILAVLDDPQYRVDSSGILKYGNIRIDSIEKKDDFLEDQG--SR
51_Physcomitrium 534 ----------YEKKL----LFQFMLKLPKNGILKQNDIFAIFNDPKYRIKNSGIIKYGNIKVDLINKKNDIFEDQK--TK
```

```
T. thermophilus 1251 --------------------------------------------------------------------------------
E. coli         1003 --------------------------------------------------------------------------------
0_Nostoc         622 --KLGYEV-VQGGTLLWIPEETHEVNKD-ISLLLVEDGQFVEAGTEVVK--DIFCQNSGVIEVTQKNDILREVVVKPGEL
1_Litchi         653 KIKPKYQ--MKFDRFFFIPEEVHTLPES--SYVMVRNNSLIGVDTRITL--NRRSQVGGLVRVERKKKR-IELKIFSGDI
2_Arabidopsis    653 KIKTKYE--MKVDRFFFIPEEVHILPES--SAIMVQNYSIIGVDTRLTL--NIRSQVGGLIRVEKKKKR-IELKIFSGDI
3_Gossypium      653 KVKPKYQ--MKVDRFFFIPEEVHILSES--SSIMVRNNSIIGVDTPITL--NTRSQVGGLVRVERKKKR-IELKIFSGNI
5_Ricinus        659 EFKPKYQ--MKVDRFFFIPEEVYILPES--SSLMVRNNSIIGVDTPITL--NTRSRVGGLVRVERKKKK-IELKIFSGDI
6_Rosa           650 EFKPKYQ--TKVDRFFFIPEEVHILPES--SSIMVRNNSIIGIDTRITL--NTRSRVGGLVRIERKKKR-IELKIFSGDI
9_Cucumis        651 DFKPKYQMQMKVDRFFFIPEEVHILPES--SSIMVRNNSIIGVATRLTL--SIRSRVGGLVRVEKKKKR-IELKIFSGDI
11_Nicotiana     651 EFRPKYQ--MKVDRFFFIPEEVHILPGS--SSIMVRNNSIVGVDTQITL--NLRSRVGGLVRVERKKKR-IELKIFSGDI
13_Syringa       657 AFRPKYQ--MKVDRFFFIPEEVHILPGS--SSIMVRNNSLIGVDTQITL--NIRSRVGGFVRVERKKKR-IELQIFSGDI
18_Liquidambar   657 EFKPRYQ--MKVDRFFFIPEEAHILPGS--SSIMVRNNSIIGVDTQITL--NTRSRVGGLVRVERKKKR-IELKIFSGDI
19_Papaver       651 EFRQKYQ--KKVDRFFFIPEEVHILSGS--SSIMVRNNSIIGIDTRITL--NIRSRVGGLVRVERKKKR-IELKIFSGDI
20_Ananas        648 EFSPKYQ--TEVDQFFFILEEVHILPGS--SLIMVRNNSIIGVDTRLALNINTRSRVRGLVRVERKKKY-IELKIFSGDI
28_Liriodendron  654 EFRPKYQ--MKVDRFFFIPEEVHILPGS--SPIMVRNNSIIGVDTRIAL--NTRSRVGGLVRVERKKKK-IELKIFSGDI
30_Magnolia      654 EFRPKYQ--MKVDRFFFIPEEVHILPGS--SSIMVRNNSIIGVDTRIAL--NTRSRVGGLVRVERKKKR-IELKIFSGDI
32_Nymphaea      657 ESRPKYQ--MKVDRFFVIPEEVHILPGS--SSIMVRNNSIIGVDTRITF--NTRSQIGGLVRIEKKKKK-IELKIFSGGI
33_Amborella     656 ELKPKYQTKYYWNKYT-----NHF---------------------------KYKKPSRRIGPSGEKKKR-IELKIFSGEI
35_Picea         626 --------------------------------------------------------------------------------
44_Ginkgo        650 GSRPKYE--IEGGRFLFIPEEVHILHES--SSIMVRNGSIIRTGTQITF--NIESQVGGLVRIERMRKK-IEVRILPGDI
51_Physcomitrium 598 TVRPRYKI-LKEGNFFLLPEEVYILDQSSFSSILVKNNSFIKAGTKITF--NISSKITGFVKIKKKFNN-FKIKILPGSI
```

```
T. thermophilus 1251 --------------------------------------------------------------------------------
E. coli         1003 --------------------------------------------------------------------------------
0_Nostoc         696 LMVDDPEAVMGRDNTFVQPGEEFQGT--------VATELRYIQYVE-TPEGPALLSRPVVEFAVPNNPDVPSTTS----V
1_Litchi         726 HFPGEADKISRHSGILIPPETGKKKLKESTGESKKLKKWIYVQRITLTKKKYFVLVRPVVTYEIAD---GINLATLFPQD
2_Arabidopsis    726 HFPDKTDKISRHSGILIPPGRGKKNSK----ESKKFKNWIYVQRITPTKKKFFVLVRPVATYEIAD---SINLATLFPQD
3_Gossypium      726 YFPGERDKISRHSGILIPPGTGKTNSK----ESKKLKNWIYVQRITPTKKKYFVLVRPVTPYEIPD---GLNLATLFPQD
5_Ricinus        732 HFPGETDKISRHSGILIPPGMVKTNSK----ESKKQKNWIYIQRIAPTRKKYFVLVRLVIIYEIAN---GINLETLFPRD
6_Rosa           723 HFPGEMDKIFRHNGILIPPGT---NSK----ESKKRNNWIYVQWITPTKKKYFVLVRPVIIYEIAD---GINLATLFPQD
9_Cucumis        726 HFPGEMDKISRHNGILIPPERVKKNSK----KSKKSKNWIYVQWITPTKKKYFVFVRPVIIYELAD---GINLVKLFPQD
11_Nicotiana     724 HFPGETDKISRHTGVLIPPGTGKRNSK----ESKKVKNWIYVQRITPSKKKFFVLVRPVVTYEITD---GINLATLFPPD
13_Syringa       730 HFPGETDKISRHSGVLIPPGTGNSNSK----ESKKLKNWIYVQRITPSKKKYFVLVRPVVTYEITD---GINLVTLFPPD
18_Liquidambar   730 HFPGETNKIARHSGILIPPGTGKTNSK----ESKKLKNWIYVQRITPTKKKHFVLVRPVVTYEIAD---GINLATLFPQD
19_Papaver       724 HFPGETDKISWHSGILIPPGTGKKNAG----DSKKLKNWIYVQRITPIKKKFFVLVRPVVTYEIAD---GINLATLFPHD
20_Ananas        723 HFPGETDKISRHSGIFIPPETEKKNSK----ESKKWKNWIYVQRITPTKKKYFVSVRPVVTYEISD---GINLATLFPRD
28_Liriodendron  727 HFPGETDKISRHSGILIPPGTGKKNSK----ESKKWKNWIYVQRITPTKKKYFVSVRPVVTYEIAD---GINLGTLFPQD
30_Magnolia      727 HFTGETDKISRHSGILIPPGTGKRNSK----ESKKWKNWIYVQRITPTKKKYFVSVRPVVTYEIAD---GINLGTLFPQD
32_Nymphaea      730 HFPGETDKISRHIGILIPPGARKKMDKGSQGKNWEGKNWVYVQRITPIKKKYFVSVRPVVTYEIAD---GINLVTLFPGD
33_Amborella     703 QFPVEMDKIFRHSGILIPPGRVKKKIK----ESKKLKNWIYVQWITPTKKKYFVLVRPVIIYEIAD---GINLETLFPQD
35_Picea         626 ---------------------------------------------------------SNVEYGIPD---GPIMATSFSLD
44_Ginkgo        723 YFPGEIHEISRHNGTLIPPGKIIFD-------EFQSVNWIYFQWITPHKEKPFVPVRSAAEYGIHD---GSNRTAPFYLD
51_Physcomitrium 674 YYPKEKQKNFKQNGILIPPGEKIF-------EQFRAKNWIYLEWIVLSKDNSFFLIRPAIEYKIIFNDNPLTLPIPFYLD
```

```
T. thermophilus 1251 --------------------------------------------------------------------------------
E. coli         1003 ---------------LAKGDGEQVA----------GGETVANWDPHTMPVIT---------------EVSGFVRFTDMID
0_Nostoc         763 SQQTGRSIQLRAVQRLPYKDSERVKSVE--GVELLRTQLVLEIEQEGEQQDHNASPLAADIELVQDTEDPEVQRLQLVIL
1_Litchi         803 PLREKDNMQLRVVNYILYGKGKPTRGISDTSIQLVRTCLVLNWGQDKKS-SSAEEVRTSFVEVSTNGMIRDFLRIDLVQS
2_Arabidopsis    799 LFREKDNIQLRVFNYILYGNGKPTRGISDTSIQLVRTCLVLNWDKN----SSLEEVRAFFVEVSTKGLIQDFIRIGLVKS
3_Gossypium      799 PFQEKDNMQLRAVNYILYGNGKPTRRISDTSIQLVRTCLVLSWDQDNKS-SFAEEVCASFVEVRTNGLIRDFLRIDLVKS
5_Ricinus        805 LLQEKDNLKLRVVNYILSGNGKPIRGISDTSIQLVRTCLVLNWDQEKKS-SSIEEARASFVEVNTNGLIRDFLRINLVKS
6_Rosa           793 PLRERDNLELRVVNYILYGNGKPIRGISGTSIQLVRTCLILNWDKNKKS-SSIEEAHASFVEVSANGLIQDFLRINLVKS
9_Cucumis        799 LLQERDNLELRVVNYILYGNGKPIRGISGTSIQLVRTCLLLNWDRDKKS-SSIEDARASFVEVSTNGLVRNFLRIDLGKS
11_Nicotiana     797 PLQERDNVQLRIVNYILYGNGKPIRGISDTSIQLVRTCLVLNWNQDKKS-SSCEEARASFVEIRTNGLIRHFLRINLVKS
13_Syringa       803 LLQERDNVQLRVVNYILYGNGKPIRGISDTDIQLVRTCLVLNWDQDKKKSSSSEEARASFVEIRTNGLIRHFLRIDLVKS
18_Liquidambar   803 LLQERDNMKLRVVNYILYGNGKPIRGISDTSIQLVRTCLVLNWDQDKKSA-SSGEAHASFVEVRTNGLIRNFLRINLVES
19_Papaver       797 LLQERDNVQLRVVNYILYGNGKPIRGIYHTSIQLVRTCLVLNWNQEKKG-SSIEEVQASFVEVRVNNLIRYFIRMDLVKS
20_Ananas        796 ILQEKDNVQLRVVNYILYGNGKSIRGIYHTSIQLVRTCLVLNWDQEQNG--FIEEVHASFVEVRANDLIRDFIRIELVKS
28_Liriodendron  800 LLQERDNVQLRVVNYILYGNGKPIRGIYHTSIQLVRTCLVLNWDQDRNG--SIEEVHASFVEVGTNDLIRDFIRIDLVKS
30_Magnolia      800 LLQERDNVQLRVVNYILYGNGKPIRGIYHTSIQLVRTCLVLNWDQDRNG--SIEEVHASFVEVGANDLIRDFIRIDLVKS
32_Nymphaea      807 MLQEKDNLRLQVVNYILYGDGKPIRGISHTSIQLVRTCLVLNWDQDKKG--SIEKVQASSAEVRANDLIRYFIRIDLVKS
33_Amborella     776 LLQEKDNLELRVVNYILYGNGKPILGISGTSIQLVRTCLMLNWDQDNKS-SSSEEAHVYFVEVSTTGLIRDFLRINLAKS
35_Picea         646 LSREGDNLQIQVSNSSSYEDGERIQVMSDTSIPLVQTCLGFDWEQIDS---IESEAYASLTSVRTNKIVSNMIQISLIKY
44_Ginkgo        793 LLGEEDNLQVQVGNYILYGDGEQIQVISDTSIQLVRTCSVLNWEQKDS---ME-EAYAFLTEVRINEVVRNFLQISLMKY
51_Physcomitrium 747 LLKEQKKIKIQTVKYILYEDSEEVEINPDTDIQLIQTCLILNWETK----VFIKEAHISFIKIRINKIIKNFFQINLIEN
```

```
T. thermophilus 1251 --------------------------------------------------------------------------------
E. coli         1043 GQTITR-QTDE--LTG--------------------------------------------------LSSLVVLDSAERT-
0_Nostoc         841 ESLVIRRDITADATQG--------STQTTLEVQDGLTIAPGSVVARTQILSKEGGIVRGVQKGTENVRRCLVLRET----
1_Litchi         882 HISYMR-KRNDPSSSG--LISDNGSDRTNINP--FYSLYF--KARVQQSLSQNQRTLHTLLNRNKKCQSLIILSSSNCFR
2_Arabidopsis    875 HISYIR-KRNNSPDSGLI-------SADHMNP--FYSISPK-SGILQQSLRQNHGTIRMFLNRNKESQSLLILSSSNCFR
3_Gossypium      878 HIFYIR-KRNDPSGSE--LISDNRSDRTNKNP--FYSIYS--NARIQQSFSQNHGTIHTLLNRNKESQSLIILSASNCFR
5_Ricinus        884 HISYISRKRNDPSGSG--PISNNGANHTNINP--FYPIYF--KTRIQQSLKQNQGTISTLLNRNKECQSLIILSSSNCFR
6_Rosa           872 HTSYIR-KRNDPLGSG--LISDNRSDRTNINP--FYSIYS--KERIQQSLRQNQGTFRTLLNRNKESQSLIILSSYNCFQ
9_Cucumis        878 DTAYMR-KRKDPSGSG--LIFNNESDRTNINP--FFSIYS--KTRVPQSPSQNQGTIRTLFNRNKERQSLIILSASNCLQ
11_Nicotiana     876 PISYIG-KRNDPSGSG--LLSDNGSDCTNINP--FSSIYSYSKAKIQQSINQPQGTIHTLLNRNKECQSLIILSAANCSR
13_Syringa       883 PISYIG-KRNNPSGSG--LLSDNGSDCTNINP--FSSIYS--KARIQHSLNQNQGTIHTLLNRNTGFQSLIILSSSNCFR
18_Liquidambar   882 PISYTG-KRNDPSGSG--WISDNGSDRTNINP--FYSTYS--KERIQQSLSQNQGTIRTLLNRNKECQSLRILSSSNCSR
19_Papaver       876 PILYTR-KRNDRGGAGLIWIPDNGSDRTNLNP--FSF---SSKARIQQTFTQHQGTIRALVNRNKESQPLIILSSSNCFQ
20_Ananas        874 TISYTG-KRYDRASSG--LIPDNGLDRTNINP--FYSKAK------IQSLSQHQGTIGTLLNRNKECQSLIILSSSNCSR
28_Liriodendron  878 PISYIG-KRDDTTGSG--LIPDNESDRTNINT--FYSKT-----R-IQSLTQHQGTIRTFLNRNKECQSFLILSSSDCSR
30_Magnolia      878 PISYIG-KRNDTAGSG--LIPDNESDRTNINT--FYSKT-----R-IQSLTQHQGTICTFLNRNKECQSFLILSSSDCSR
32_Nymphaea      885 PILYTG-KRNDGSGS---VIPDTGSYCANTNL--FSSKVK------MKSLSQHQGTVRTFLNRNKEGQSLIVFSSSNCSR
33_Amborella     855 NICYIR-KRNDPLGSG--LISDNRSDCT--NP--FYSIYS--KEKIQQSLRQNQGTIHTLLNRSKESQSLIILSSSNCFQ
35_Picea         723 PLFFMGR-RDNKASSN--LMFHNKLDHT--NL--FYSN------GERQLISKHQGTICSLYNGEEDSGSFMVLSPSDCFR
44_Ginkgo        869 PG---GK-RKNVTGSK--FLFHNRSDQT--NT--FSSN------RGSQFFSKHQGTIRTLPNEEKEGGSFAVLSPSDCSR
51_Physcomitrium 823 INLMNKK-KNNNIILN--YLFKKKR----------YII-NQKDCEKILLLSKTWGIIRTPSNKNQEKSFFLILSPFNLFQ
```

```
T. thermophilus 1251 --------------------------------------------------------------------------------
E. coli         1069 ------------------------------------------------------------------------------AG
0_Nostoc         909 ---------------------------------------------------------------DLITVNTSTQPKVKM--
1_Litchi         955 MGPFNDI-KYHNVIKQSIHI------------QKGSLIPIRNSLGPLGT-VLQIANFYSFYY--LITYNQISVT--KYWK
2_Arabidopsis    944 MGPFNHV-KHHNVINQSI--------------KKNTLITIKNSSGPLGT-ATPISNFYSFLP--LLTYNQISLI--KYFQ
3_Gossypium      951 MGPFNDV-KYHNVIKQSI--------------KKDPLIPIKNLLGPLGT-APKIANFYSSFYP-LITHNQTSVA--KYFE
5_Ricinus        958 MDPFNDV-KHHNVIKESI--------------KRDPIIPIRNSLGPLGT-ALQIANLYLFYHLNLITHNRISVT--KYLK
6_Rosa           945 MSPFNDV-KYYDGIKESIKR------------DRDSLIQITNLLGPLGT-ASQIDLFYSFYHL--LTHNHISVTKYFYLQ
9_Cucumis        951 MDLFNDVKDY-NVIKESS--------------KKDPLISIRNSLGPLGA-APQIVNFYSFYD--LITHNPISLT--KYLQ
11_Nicotiana     951 MGPFKDV-KYHSVIKKSI--------------KKDPLIPIRNSLGPLGT-SLPIENFYSSYH--LITHNQILVT--NYLQ
13_Syringa       956 MGPFNDVIKYHNVIKESIKI------------TKDPLIPLKNSLGPFGT-AFTIANFFSFYH--LITHNQILVN--NYLQ
18_Liquidambar   955 MGPFNDV-KYHNVIKESI--------------KRDPLIPIRNSLGPLGT-ALQIANFYSFYH--LITHNKILVT--KYLQ
19_Papaver       950 MGPFNSVKYNDGVTKEST--------------KRDLRISILNSLGPLGI-VPKFVNFS---SYHLITHNQILVK--KYLL
20_Ananas        943 IGPFNSS-KYNNLTK-----------------ESDPLIPIRDSLGLLGAIVPKIANFYSSYH--LITHNQIVLK--KYLL
28_Liriodendron  947 IGPFNGS-KSHKVTKESI--------------KEDPMIPIRNSLGPLGT-VPKIANFYSSYY--LITHNQILLN--KYLL
30_Magnolia      947 IGPFNGS-KSHKVTKESI--------------KEDPMIPIRNSLGPLGT-VSKIANFYSSYY--LITHNQILLN--KYLL
32_Nymphaea      953 IN----VSKYHNVTKESIKE------------KEDTPIPILNLLGPLGT-VPKIHNFSPSYH--SITHNEILLNKYLILD
33_Amborella     926 MSPFKDV-QYSNGIKESI--------------KVEPLIPIRNSLGPLGT-SSQIENFYY----LLKTHNQISVT--KYLQ
35_Picea         790 IVLFNDSKCYDTV-NKSN--------------REDPMRKIIEFSGLLGHLH-SITNRFPSS--HFLTYKKVLSKKHS--I
44_Ginkgo        933 TVLFSGSKYYDTV-KRSI--------------QEDPMMQIIELSGLLGNLH-SIANRFPSP--HLITYNKVLSNKHS--I
51_Physcomitrium 889 TILFDKTKQNLKIENNVEKLFTYEPKKIIKTFNIEKRKNFVEFLGLLGYLQN-ITKSFQLFSCKKFSDKSI---PINFSI
```

```
T. thermophilus 1251 --------------------------------------------------------------------------------
E. coli         1071 GKDLRPALKIV--------DAQG---------------------------------------------------------
0_Nostoc         924 GDLLVAGT---EVATGIFTEESGQVTNVK---------------------------------------KLG-VKSEELGV
1_Litchi        1017 LDNLKQTFQIC---KFYLMDENGRIYNPDPDSKIVLNPFNLNWYFLHH--------NYCEE--MSTIISLGQFICENVCI
2_Arabidopsis   1004 LDNLKYIFQKI---NSYLIDENGIILNLDPYSNVVLNPFKLNWYFLHQNYHHNYCE----E--TSTIISLGQFFCENVCI
3_Gossypium     1012 LDNLKQAFQVL---NYYLIAENGRIYNFDPCRNIFLNAVNLNWYFPHHHYHH----NYCEE--TSTIISLGQFICENVCI
5_Ricinus       1020 LDNLKQTFRVL---KYYLMDENGRVVNPDPCSNSVLNPFNLNWYFLHHNYHHNYCHNYCEE--SFTIISLGQFICENVCM
6_Rosa          1009 LDNLKQTFQVF---KYYLMDENGRISNSDPCSSILLNPFNLNWHFLDH--------NYCEE--TSTIISLGQFICENLCI
9_Cucumis       1011 LDNLKQTFQVL---KYYLMDENGGIFNSDPCSNIVFNTFNLNWHFLHHNYHNNYCEET--P--TRTRISLGHFFFENVCI
11_Nicotiana    1011 LDNLKQTFQVIK-FKYYLMDENGKIFNPDPCRNIILNPFNLNWYFLHH--------NYCEE--TSKIISLGQFICENVCI
13_Syringa      1019 LDNLKQTFQVI---KYYLMDENEKIYNPEPGSNIILNPFNLNWYFLHH--------NYCQE--TSTIISLGQFICENVCI
18_Liquidambar  1015 LDNLKQTFQVL---NYYLMDENGRIYNPDPCSNIILNPFNLNWYFLHH--------NYCEE--TSTIISLGQFICENVCI
19_Papaver      1010 LDNLKQTFTF-QGLKYYLLDETGRIYNPNLGSHIILNPLNLNWFFLQH--------DYCEE--RATIINLGQFICENVCI
20_Ananas       1001 LDNLKQIFQVLQVLKYCLIDENRRIYNPDPCSNIILNPFHLNWCFFHH--------DYCEE--TSTKISLGQFICENVCL
28_Liriodendron 1007 LDNLKQTFQVL---KYYLMDENGRIYNPNLHSNIIFNPFDLNWCFLRH--------DYCEE--TSTIISLGQFICENVCI
30_Magnolia     1007 LDNLKQTSQVL---KYYLMDENGRIYNPDPRSNIIFNPFDLNWCFLRH--------DYCEE--TSTIISLGQFICENVCI
32_Nymphaea     1014 NNNPKQTFQLL---KYFLVDENGRISNANPCSDIIFNLFGS--CFLPH--------DYCKGTSTTRIISLGQFICENVCL
33_Amborella     984 LDNFTQPFQVL---QYYLMDENGIVYNSDPCSNTRLNPFNLNWHFFHNNYDNNYYQK------KSPIISLGRFFCENVCI
35_Picea         850 FHN---SFNTFQVPKYYFMDENTRISHFDPCRNIISNLLGPNWCSSSS--------EFCKK--IFPVVSPGQLIPESVCI
44_Ginkgo        993 SDN---SGKVSQVSKCYFMGGNTGILNFDSCRNIIFNLFNSNWCSPLS--------NFCKK--KLPAVSLGQLIRESVCI
51_Physcomitrium 965 IDNLKKKIK---ISKWFFLNENKKVQKFFLTQNTILSL--LNWSFPIF--------DLAKK--KTQLFNLGHFFCDGLSI
```

```
                                                                               Trigger loop helix2

                                                                               β'b9:D1251-V1281
T. thermophilus 1251 ---------------------------------------------------------------DITQGLPRVIELFEARR
E. coli         1086 ----------NDVLI------PGTDMPAQYFLPGKAIVQLEDGVQISSGDTLARIPQESGGTKDITGGLPRVADLFEARR
0_Nostoc         961 NSETPNSS-----LQ-TQNYAITIRLGRPYRVSPGAVLQIEDGDLVQRGDNLVLLVFERAKTGDIIQGLPRIEELLEARK
1_Litchi        1084 ATNGPHLK-SGQVLIVQ-VGSVVIRSAKPYLATPGATVHGHYGEILYEGDTLVTFIYEKSRSGDITQGLPKVEQVLEVRS
2_Arabidopsis   1075 AKKEPHLK-SGQVLIVQ-RDSAVIRSAKPYLATPGAKVHGHYSEILYEGDTLVTFIYEKSRSGDITQGLPKVEQVLEVRS
3_Gossypium     1083 AKSGPRLK-SGQVFIVQ-ADSIVIRSAKPYLATPGATVHGHYGETLYEGDTLVTFIYEKSRSGDITQGLPKVEQVLEVRS
5_Ricinus       1095 AKNGPHLK-SGQVIIIH-IGSVVIRSAKPYLATPGATVHGHYGEILYEGDTLVTFIYEKSRSGDITQGLPKVEQVLEVRS
6_Rosa          1076 AKKGSPLK-SGQVIVVQ-LDSLVIRSAKPYLATPGATVHGHYGEILSEGDTLVTFIYEKSRSGDITQGLPKVEQVLEVRS
9_Cucumis       1084 AKNRPHLK-SGQIIVE-VDSVVIRSAKPYLATSGATVHRHYGEMLYEGDTLVTFIYEKSRSGDITQGLPKVEQVLEVRS
11_Nicotiana    1080 AKNGPPLK-SGQVILVQ-VDSIVIRSAKPYLATPGATVHGHYGETLYEGDTLVTFIYEKSRSGDITQGLPKVEQVLEVRS
13_Syringa      1086 AKNTPHLK-SGQVILVQ-VDSVVIRSAKPYLATPGATVHGHYGEILYEGDTLITFIYEKSRSGDITQGLPKVEQVLEVRS
18_Liquidambar  1082 AKNGPHLK-SGQVLIVQ-VDSVVIRSAKPYLATTGATVHGHYGEILYEGDTLVTFIYEKSRSGDITQGLPKVEQVLEVRS
19_Papaver      1079 SKYGPRLK-AGQVLIIR-VGSLVIRSAKPYLATPGATVHGHYGETLSEGDTLVTFIYEKSRSGDITQGLPKVEQVLEVRS
20_Ananas       1071 FKYEPHVKKSGQILIVN-VDSLVIRSAKPYLATPGATVHGHYGKILYEGDTLVTFIYEKSRSGDITQGLPKVEQILEVRS
28_Liriodendron 1074 SKYGPHIK-SGQVLIVH-VDSLVIRSAKPHLATPGATVHGHYGEILSEGDTLVTFIYEKSRSGDITQGLPKVEQVLEVRS
30_Magnolia     1074 SKYGPHIK-SGQVLIVH-VDSLVIRSAKPHLATPGATVHGHYGEILYEGDTLVTFIYEKSRSGDITQGLPKVEQVLEVRS
32_Nymphaea     1081 SKHRTRIK-SGQVIMVY-LDSFIIRSAKPYLATRGATVHGDYGEIFYEGDTLVTFIYEKSRSGDITHGLPKVEQVLEVRS
33_Amborella    1055 VKHGPHLK-SGQVIIVQ-IDSVVIRLAKPYLATPGATVHGHYGEKLSEGDTLVTFIYEKSRSGDITQGLPKVEQVLEVRS
35_Picea         917 SEDEPLPE-SGQIIAVD-EESLVIRSAKPYLATRKATVHGHYGEIIDKGDTLITLIYERFKSSDIIQGLPKVEQLSEARL
44_Ginkgo       1060 SEDKPLSG-SGQIIAVH-EEYLVIRSAKPYLATRRAAVHGHYGEFLDEGDTLITLIYERSKSGDTIQGLPKVEQLSEARS
51_Physcomitrium1030 AEYPTFSE-SGQIIAIYDDLSLVIRLAKPYLATGGAIIHNNYGEIVKEGDILITLIYERLKSGDIIQGLPKVEQLLEARL


                   β'b9: D1251-V1281                                    β'b10:V1313-N1404
T. thermophilus 1268 PKAKAVISEIDGVVRIEET--EEKLSV-FVE-S-EGFSKEYKLPKEARLLVKDGDYVEAGQPLTRGAIDPHQLLEAKGP-
E. coli         1150 PKEPAILAEISGIVSFGKE--TKGKRRLVITPVDGSDPYEEMIPKWRQLNVFEGERVERGDVISDGPEAPHDILRLRGV-
0_Nostoc        1035 PKEACILARRAGEVKVVYGDGDEAIAIKVV--ESNGVVTDYPLGPGQNLIVPDGSHISAGQPLTDGPSNPHEILEIFFSL
1_Litchi        1162 IDSISMNLEKRV------E-------------GWNA--------------------------------------RITRIL
2_Arabidopsis   1153 IDSISLNLEKRI------K-------------GWNK--------------------------------------CITRIL
3_Gossypium     1161 IDSISMNLEKRI------E-------------GWNE--------------------------------------CITRIL
5_Ricinus       1173 IDSISINLEKRV------G-------------GWNE--------------------------------------CIPRIL
6_Rosa          1154 IDSISMNLEKRV------E-------------GWNE--------------------------------------CITRIL
9_Cucumis       1162 IDSISMSLEKRI------E-------------GWNE--------------------------------------RITRIL
11_Nicotiana    1158 VDSISMNLEKRI------E-------------GWNK--------------------------------------CITRIL
13_Syringa      1164 IDSISMNLEKRV------E-------------GWNE--------------------------------------RITRIL
18_Liquidambar  1160 IDSISMNLEKRV------E-------------GWND--------------------------------------RITRIL
19_Papaver      1157 LDSISMNLEKRV------E-------------GWNE--------------------------------------RITRIL
20_Ananas       1150 IDSLSMNLERRV------E-------------GWNE--------------------------------------RIPRIL
28_Liriodendron 1152 IDSISMNLEKRI------E-------------GWNE--------------------------------------HITRIL
30_Magnolia     1152 IDSISMNLEKRI------E-------------GWNE--------------------------------------RITRIL
32_Nymphaea     1159 IDSISMNLEKRV------E-------------GWNE--------------------------------------HITGIL
33_Amborella    1133 IDSISMNLEKRV------E-------------GWND--------------------------------------LLTRIL
35_Picea         995 NNSISMNLKESF------E-------------NWTG--------------------------------------DMTRFL
44_Ginkgo       1138 INPIPRNLEESF------E-------------DWNE--------------------------------------DMTRSL
51_Physcomitrium1109 TNPVSINLEKGF------G-------------EWNK--------------------------------------DMTNFF


                          β'a17:L1348-K1354      β'a18:D1365-L1389     β'a19:L1395-E1401
                                            β'b10:V1313-N1404
T. thermophilus 1342 --------------EAVERYLVEEIQKVYRAQGVKLHDKHIEIVVRQMMKYVEVTDPG-DSRLLEGQVLEKWDVEALNER
E. coli         1227 --------------HAVTRYIVNEVQDVYRLQGVKINDKHIEVIVRQMLRKATIVNAG-SSDFLEGEQVEYSRVKIANRE
0_Nostoc        1113 GSEDGVYACASHALQKVQTFLVNEVQMVYQSQGIDISDKHIEVIVRQMTNKVRIDD-GGDTTMLPGELVELRQVEQVNEA
1_Litchi        1185 GIPWGFLIGAELTIVQSRISLVNKIQKVYRSQGVQIHNRHIEIIVRQITSKVLVSEDGMSNVFLPGELIGLLRAERTGRA
2_Arabidopsis   1176 GIPWGFLIGAELTIVQSRISLVNKIQKVYRSQGVQIHNRHIEIIVRQITSKVLVSEEGMSNVFLPGELIGLLRAERTGRA
3_Gossypium     1184 GIPWGFVIGAELTIVQSRLSLVNKIQKVYRSQGVQIHNRHIEIIVRQITSKVLVSEDGMSNVFLPGELIGLLRAERTGRA
5_Ricinus       1196 GIPWGFLIGTELTIVQSRISLVNKIQRVYRSQGVQIHNRHIEIIVRQITSKVLVSEDGMSNVFSPGELIGLLRAERTGRA
6_Rosa          1177 GIPWGFLIGAELTIAQSRISLVNKIQKVYRSQGVQIHNRHIEIIVRQITSKVLVSEDGMSNVFSPGELIGLLRAERTGRA
9_Cucumis       1185 GIPWGFLIGSELTIVQSRISLVNKIQKVYRSQGVEIHNRHIEIIVRQITSKVLVSEDGMSNVFSPGELIGLLRAERTGRA
11_Nicotiana    1181 GIPWGFLIGAELTIAQSRISLVNKIQQVYRSQGVQIHNRHLEIIVRQITSKVLVSEDGMSNVFSPGELIGLLRAERMGRA
13_Syringa      1187 GMPWGFLIGAELTIVQSRISLVNKIQKVYRSQGVQIHNRHIEIIVRQITSKVLVSEDGMSNVFSPGELIGLLRAERMGRA
18_Liquidambar  1183 GIPWGFLIGAELTIVQSRISLVNKIQKVYRSQGVQIHNRHIEIIVRQITSKVLVSEDGMSNVFSPGELIGLLRAERTGRA
19_Papaver      1180 GIPWGFLIGAELTIAQSRISLVNKIQKVYRSQGVQIHNRHIEIIVRQITSKVLVSEDGMSNVFLPGELIGLLRAERTGRA
20_Ananas       1173 GIPWGFLIGAELTIAQSCISLVNKIQKVYRSQGVQIHNRHIEIIVRQVTSKVLVSEDGMSNVFSPGELIGLLRAERAGRA
28_Liriodendron 1175 GIPWGFLIGAELTIAQSRISLVNKIQKVYRSQGVQIHNRHIEIIVRQITSKVLVSEDGMSNVFSPGELIGLLRAERTGRA
30_Magnolia     1175 GIPWGFLIGAELTIAQSRISLVNKIQKVYRSQGVQIHNRHIEIIVRQITSKVLVSEDGMSNVFSPGELIGLLRAERTGRA
32_Nymphaea     1182 GIPWGFLIGAELTIAQSRISLVNKIQKVYRSQGVQIHNKHIEIIVRQITSKVLVSEDGMSNVFSPGELIGLLRAERAGRA
33_Amborella    1156 GIPWGFLIGAELTIVQSRISLVNKIQKVYRSQGVQIHNRHIEIIVRQITSKVVVSEDGMSNVFLPGELIGLLRAERTGRA
35_Picea        1018 GSLWGLFISARITMEQSQIHLVNQIQKVYRSQGVRICDKHIEIIVRQMTSKVLISEDGMANVFSPGELIGLSRAQRMDRA
44_Ginkgo       1161 GSLWGLFISARITMEQSQIHLVNQIQKVYRSQGVRIYDKHIEIIVRQMTSKVFISGDGMADVFSPGELIELSRAQRMNRA
51_Physcomitrium1132 GSLWGYFLSAQISMEQSQVNLVNQIQKVYRSQGVNISDKHIEIIVRQMTSKVFTLEDGMTNGFLPGELIEFARAKRMNRA
```

```
                                                  Clamp            Switch 5
                                                      β'a20:F1440-G1477
                                        β'b11:V1415-G1477
T. thermophilus 1407 LIAEGKTPVAWKPLLMGVTKSALSTKSWLSAASFQNTTHVLTEAAIAGKKDELIGLKENVILGRLIPAGTGSDFVRFTQV
E. coli         1292 LEANGKVGATYSRDLLGITKASLATESFISAASFQETTRVLTEAAVAGKRDELRGLKENVIVGRLIPAGTGYAYHQDRMR
0_Nostoc        1192 MAITGGARAQYTPVLLGITKASLNTDSFISAASFQETTRVLTEAAIEGKSDWLRGLKENVIIGRLIPAGTGYNTYEETSA
1_Litchi        1265 LE----EAIRYRAILLGITRASLNTQSFISEASFQETARVLAKAALRGRIDWLKGLKENVVLGGMIPVGTGF-KGLVHCS
2_Arabidopsis   1256 LE----EAICYRAVLLGITRASLNTQSFISEASFQETARVLAKAALRGRIDWLKGLKENVVLGGVIPAGTGFNKGLVHCS
3_Gossypium     1264 LE----EAICYRAVLLGITRASLNTQSFISEASFQETARVLAKAALRGRIDWLKGLKENVVLGGMIPAGTGF-KGLVHRS
5_Ricinus       1276 LE----EAICYRAILLGITRASLNTQSFISEASFQETARVLAKAALRGRIDWLKGLKENVVLGGMIPVGTGF-KGLVQGS
6_Rosa          1257 LE----EAICYRAILLGITKASLNTQSFISEASFQETARVLAKAALRGRIDWLKGLKENVVLGGMIPVGTGF-KGFVPRS
9_Cucumis       1265 LE----EAICYRAVLLGITKASLNTQSFISEASFQETARVLAKAALRGRIDWLRGLKENVVLGGMIPVGTGF-RELAHRS
11_Nicotiana    1261 LE----EAICYRVVLLGITRASLNTQSFISEASFQETARVLAKAALRGRIDWLKGLKENVVLGGVIPVGTGF-KGLVHPS
13_Syringa      1267 LE----EAVCYRAVLLGITRASLNTQSFISEASFQETARVLAKAALRGRIDWLKGLKENVVLGGMIPLGTGL-KGLVPPS
18_Liquidambar  1263 LE----EAICYRAILLGITKASLNTQSFISEASFQETARVLAKAALRGRIDWLKGLKENVVLGGMIPVGTGF-KGLVHHS
19_Papaver      1260 LE----EAICYRAVLLGITRASLNTQSFISEASFQETARVLAKAALRGRIDWLKGLKENVVLGGMIPVGTGF-KGLVYHS
20_Ananas       1253 LD----ESICYRAILLGITRASLNTQSFISEASFQETARVLAKAALRGRIDWLKGLKENVVLGGIIPVGTGF-QKFVHRS
28_Liriodendron 1255 LE----EGICYRAILLGITRASLNTQSFISEASFQETARVLAKAALRGRIDWLKGLKENVVLGGMIPVGTGF-KGLVHRS
30_Magnolia     1255 LE----EAICYRAILLGITRASLNTQSFISEASFQETARVLAKAALRGRIDWLKGLKENVVLGGMIPVGTGF-KGLVHRS
32_Nymphaea     1262 LE----EAICYRAVLLGITRASLNTQSFISEASFQETARVLAKAALRGRIDWLKGLKENVVLGGMIPVGTGF-KRFVHRS
33_Amborella    1236 LE----EAICYRAILLGITKASLNTQSFISEASFQETARVLAKAALRGRIDWLKGLKENVVLGGMIPVGTGF-KGLVHCS
35_Picea        1098 LE----EAIYYQTMLLGITRASLNTQSFISEASFQETARVLAKAALQGRIDWLKGLKENVILGGIIPAGTGQ-H--IHRS
44_Ginkgo       1241 LE----EAIYYRTVLLGITRASLDTQSFISGASFQETARVLAKAALRGRIDWLKGLKENVILGGIIPAGTGF-KRFLRHS
51_Physcomitrium1212 LE----EVIPYKPVLLGITKASLNTQSFISEASFQETTRVLAKAALRGRIDWLKGLKENVILGGIIPTGTGCEEVLWQIT

T. thermophilus 1487 VDQKTLKAIEEARK-E-AVEAKER---------PAARRGVK-----REQP-GKQA-----------------------
E. coli         1372 RR-----AAGEAPAAP-QVTAEDA---------SASLAELL-----NAGLGGSDNE----------------------
0_Nostoc        1272 IDDY-ATDI-----SSSVLDEVDDPLDMVLDDRTARTYNLDAPTLGEPSYGSRRAERSILDDDDDLIADEVADEEDYEDD
1_Litchi        1340 R-QHNNILLETQKNT--LFGGV---RDILLHHRELFDFCIS-----------KTLR-----DTSEQSL-----------
2_Arabidopsis   1332 R-QHTNIILEKKTKNLALFEGDMR--DILFYHREFCDSSIS-----------KSDFSRI--------------------
3_Gossypium     1339 R-QHNNILLETKKKN--FFGGEMR--DIFFHHRELFDSCIS-----------NNLH-----DTSGRSF-----------
5_Ricinus       1351 R-QYKNIPLKTKKNN--LFGGEFRDRDILFHHRELFYSCIS-----------KNFY-----DTSEQSF-----------
6_Rosa          1332 R-QHNNISLETKNKS--LFEGEMR--DILVHHRELFDFCIS-----------KNLH-----DTSEQSF-----------
9_Cucumis       1340 R-QHNNIPLEPPPKK--IFEGEMR--DILFHHKELFDFFIS-----------TNLH-----DTSEQAF-----------
11_Nicotiana    1336 K-QHNNIPLETKKKN--LFEGEMR--DILFHHKKLFDSCLS-----------KNFH-----DIPEQSF-----------
13_Syringa      1342 K-QDSNSPLETKKNN--LFEGEMR--DILFHHRKLFDSCLS-----------KNFH-----DTSEQSF-----------
18_Liquidambar  1338 R-QHNNIPLETKKKN--LFEGEMR--DILFHHRELFHSCIS-----------KNFH-----DISEQSF-----------
19_Papaver      1335 R-QHSNIPFEIKKNN--LFRGGFR--DILFQHKELFDSYIP-----------KNIH-----DPSEQLF-----------
20_Ananas       1328 R-QDKNIYLEIKRKN--LFELEMR--DILLHHRELFCSCAT-----------NNFHETNLHETSEQSF-----------
28_Liriodendron 1330 R-QHNNIPLEIKKKN--LFEGEMR--DILFHHRELLSSCIP-----------KNFH-----DTSEQSF-----------
30_Magnolia     1330 R-QHNNIPLEIKKKN--LFEGEMR--DILFHHRELLSSCIP-----------KNFH-----DTSEQSF-----------
32_Nymphaea     1337 R-EYNNIPLEIQKKN--FFGGEMR--DILFHHRELFCSCIP-----------KPK---SFHNTSEQPF-----------
33_Amborella    1311 R-KHNNIPLEPKNKN--LFEWEMR--DILFHHRELFSFCIS-----------KNGTSSLFFTLKKKKRE-----EVWGEM
35_Picea        1171 G-KRNGMDPRMGNRN--LFSKKVK--DIFFHYHKVSFFSIQ-----------ENSHNILKQPFK---------------
44_Ginkgo       1316 E-ERNKIDSRTGNKN--LFNNKVK--DIFSHHGKVSVSPIK-----------DNYHNTLKQPLCENSVD-----K-----
51_Physcomitrium1288 LEKQKNILLKKNK--SKLFHNKVK--DIFLYK-KLSISFTS-----------EKIHKNY--------------------

T. thermophilus      --------------------------------------------------------------------------------
E. coli              --------------------------------------------------------------------------------
0_Nostoc        1346 DEDEDDFDDE----------------------------------------------------------------------
1_Litchi        1386 ----RGFNKS----------------------------------------------------------------------
2_Arabidopsis        --------------------------------------------------------------------------------
3_Gossypium     1386 ----IGIEFND--S------------------------------------------------------------------
5_Ricinus       1400 ----IGFNDS----------------------------------------------------------------------
6_Rosa          1379 ----FGFNDS----------------------------------------------------------------------
9_Cucumis       1387 ----LGFNDS----------------------------------------------------------------------
11_Nicotiana    1383 ----IGFNDS----------------------------------------------------------------------
13_Syringa      1389 ----IGFNDS----------------------------------------------------------------------
18_Liquidambar  1385 ----MGFNDS----------------------------------------------------------------------
19_Papaver      1382 ----TGFNDS----------------------------------------------------------------------
20_Ananas       1380 ----MRFNDS----------------------------------------------------------------------
28_Liriodendron 1377 ----TGFNDS----------------------------------------------------------------------
30_Magnolia     1377 ----TGFNDS----------------------------------------------------------------------
32_Nymphaea     1386 ----YTMGSNP--ISGFIIS------------------------------------------------------------
33_Amborella    1370 TRRYWNINLEEMMEAGVHFGHGTKKWNPRMAPYISAKRKGIHIINLTRTARFLSEACDLVFDAASRGKQFLIVGTKNKAA
35_Picea             --------------------------------------------------------------------------------
44_Ginkgo            --------------------------------------------------------------------------------
51_Physcomitrium     --------------------------------------------------------------------------------
```

```
T. thermophilus          --------------------------------------------------------------------------------
E. coli                  --------------------------------------------------------------------------------
0_Nostoc                 --------------------------------------------------------------------------------
1_Litchi                 --------------------------------------------------------------------------------
2_Arabidopsis            --------------------------------------------------------------------------------
3_Gossypium              --------------------------------------------------------------------------------
5_Ricinus                --------------------------------------------------------------------------------
6_Rosa                   --------------------------------------------------------------------------------
9_Cucumis                --------------------------------------------------------------------------------
11_Nicotiana             --------------------------------------------------------------------------------
13_Syringa               --------------------------------------------------------------------------------
18_Liquidambar           --------------------------------------------------------------------------------
19_Papaver               --------------------------------------------------------------------------------
20_Ananas                --------------------------------------------------------------------------------
28_Liriodendron          --------------------------------------------------------------------------------
30_Magnolia              --------------------------------------------------------------------------------
32_Nymphaea              --------------------------------------------------------------------------------
33_Amborella        1450 DSVAGAAIKARCHYVNKKWLGGMLTNWYTTETRLHKFRDLRTEQKTGRLNRLPKRDAAVLKRQLSHLQTYLGGIKYMTGL
35_Picea                 --------------------------------------------------------------------------------
44_Ginkgo                --------------------------------------------------------------------------------
51_Physcomitrium         --------------------------------------------------------------------------------

T. thermophilus          --------------------------------------------------------------------------------
E. coli                  --------------------------------------------------------------------------------
0_Nostoc                 --------------------------------------------------------------------------------
1_Litchi                 --------------------------------------------------------------------------------
2_Arabidopsis            --------------------------------------------------------------------------------
3_Gossypium              --------------------------------------------------------------------------------
5_Ricinus                --------------------------------------------------------------------------------
6_Rosa                   --------------------------------------------------------------------------------
9_Cucumis                --------------------------------------------------------------------------------
11_Nicotiana             --------------------------------------------------------------------------------
13_Syringa               --------------------------------------------------------------------------------
18_Liquidambar           --------------------------------------------------------------------------------
19_Papaver               --------------------------------------------------------------------------------
20_Ananas                --------------------------------------------------------------------------------
28_Liriodendron          --------------------------------------------------------------------------------
30_Magnolia              --------------------------------------------------------------------------------
32_Nymphaea              --------------------------------------------------------------------------------
33_Amborella        1530 PDIVIIVDQQEEYTALRECITLGIPTICLIDTNCDPDLADISIPANDDAIASIRLILNKLVFAILLYDISGVEVGQHFYW
35_Picea                 --------------------------------------------------------------------------------
44_Ginkgo                --------------------------------------------------------------------------------
51_Physcomitrium         --------------------------------------------------------------------------------

T. thermophilus          --------------------------------------------------------------------------------
E. coli                  --------------------------------------------------------------------------------
0_Nostoc                 --------------------------------------------------------------------------------
1_Litchi                 --------------------------------------------------------------------------------
2_Arabidopsis            --------------------------------------------------------------------------------
3_Gossypium              --------------------------------------------------------------------------------
5_Ricinus                --------------------------------------------------------------------------------
6_Rosa                   --------------------------------------------------------------------------------
9_Cucumis                --------------------------------------------------------------------------------
11_Nicotiana             --------------------------------------------------------------------------------
13_Syringa               --------------------------------------------------------------------------------
18_Liquidambar           --------------------------------------------------------------------------------
19_Papaver               --------------------------------------------------------------------------------
20_Ananas                --------------------------------------------------------------------------------
28_Liriodendron          --------------------------------------------------------------------------------
30_Magnolia              --------------------------------------------------------------------------------
32_Nymphaea              --------------------------------------------------------------------------------
33_Amborella        1610 QIGGFQVHAQVLITSWVVIAILLGSAILAVRNPQTIPTDGQNFFEYVLEFIRDVSKTQIGEEYGPWVPFIGTLFLFIFVS
35_Picea                 --------------------------------------------------------------------------------
44_Ginkgo                --------------------------------------------------------------------------------
51_Physcomitrium         --------------------------------------------------------------------------------
```

```
T. thermophilus      --------------------------------------------
E. coli              --------------------------------------------
0_Nostoc             --------------------------------------------
1_Litchi             --------------------------------------------
2_Arabidopsis        --------------------------------------------
3_Gossypium          --------------------------------------------
5_Ricinus            --------------------------------------------
6_Rosa               --------------------------------------------
9_Cucumis            --------------------------------------------
11_Nicotiana         --------------------------------------------
13_Syringa           --------------------------------------------
18_Liquidambar       --------------------------------------------
19_Papaver           --------------------------------------------
20_Ananas            --------------------------------------------
28_Liriodendron      --------------------------------------------
30_Magnolia          --------------------------------------------
32_Nymphaea          --------------------------------------------
33_Amborella    1690 NWSGALLPWKIIELPHGELAAPTNDINTTVALALLTRFHKTLIT
35_Picea             --------------------------------------------
44_Ginkgo            --------------------------------------------
51_Physcomitrium     --------------------------------------------
```

**Fig. S7:** view of the catalytic core from the *E. coli* RNAP (PDB entry: 3LU0 (Opalka *et al*., 2010)) manually fitted into the envelope of PEP using Chimera (Pettersen *et al*., 2004).

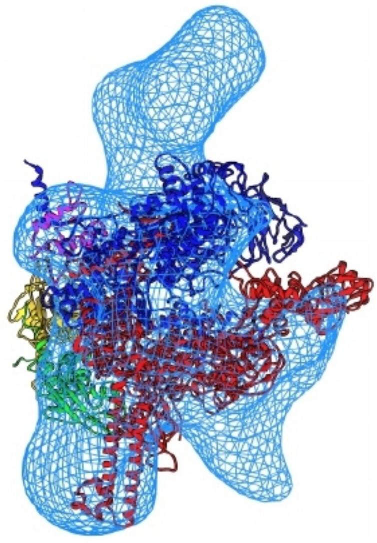

**Figures S8a and S8b** overall shape of the a) human RNA polymerase II (EMDB entry: EMD-2194; Kassube *et al*., 2013) and b) yeast RNA polymerase III (EMDB entry: EMD-1753; Vanini *et al*., 2010) solved at 25 and 21 Å respectively.

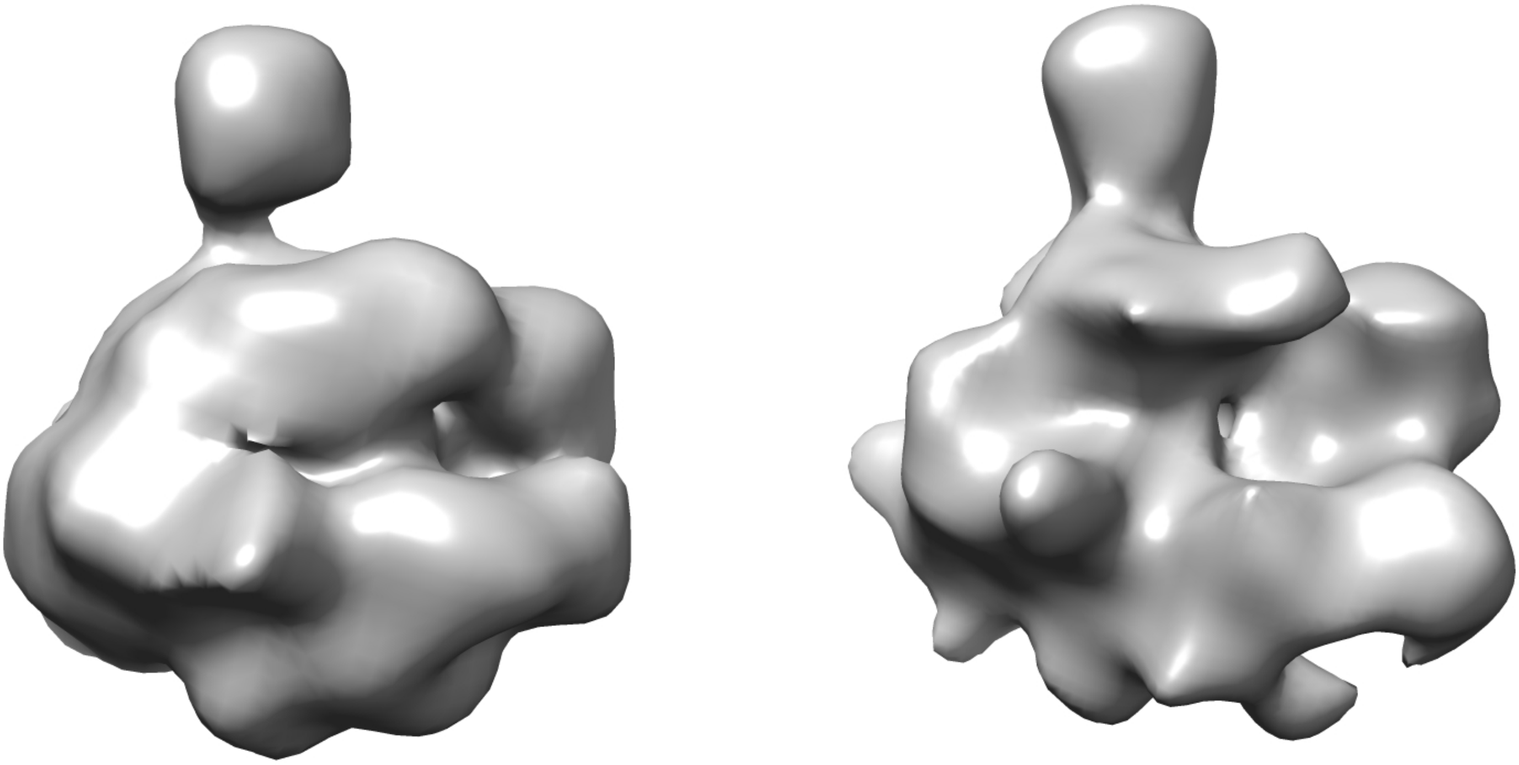

***Figure S9 :*** FSC curve for the PEP 3D reconstruction calculated between two independent half maps (gold standard FSC). The dotted line represents the FSC=0.143 cutoff used to determine the resolution.

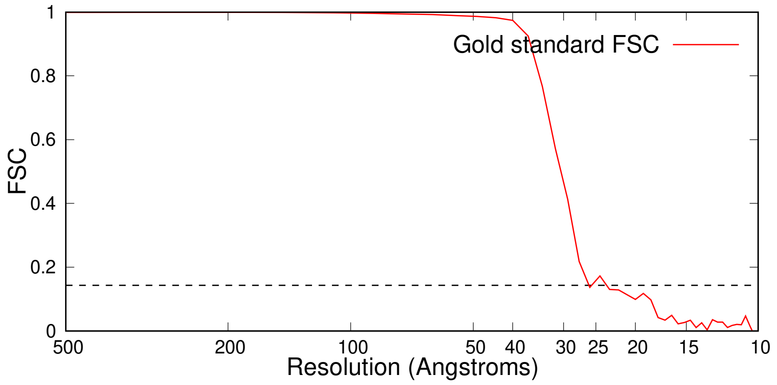
